# Epstein-Barr virus inactivates the transcriptome and disrupts the chromatin architecture of its host cell in the first phase of lytic reactivation

**DOI:** 10.1101/573659

**Authors:** Alexander Buschle, Paulina Mrozek-Gorska, Stefan Krebs, Helmut Blum, Filippo M. Cernilogar, Gunnar Schotta, Dagmar Pich, Tobias Straub, Wolfgang Hammerschmidt

## Abstract

Epstein-Barr virus (EBV), a herpes virus also termed HHV 4 and the first identified human tumor virus, establishes a stable long-term latent infection in human B cells, its preferred host. Upon induction of EBV’s lytic phase the latently infected cells turn into a virus factory, a process, that is governed by EBV. In the lytic, productive phase all herpesviruses ensure the efficient induction of all lytic viral genes to produce progeny, but certain of these genes also repress the ensuing antiviral responses of the virally infected host cells, regulate their apoptotic death or control the cellular transcriptome. We now find that EBV causes previously unknown massive and global alterations in the chromatin of its host cell upon induction of the viral lytic phase and prior to the onset of viral DNA replication. The viral initiator protein of the lytic cycle, BZLF1, binds to >10^5^ binding sites with different sequence motifs in cellular chromatin and in a concentration dependent manner. Concomitant with DNA binding, silent chromatin opens locally as shown by ATAC-seq experiments, while previously wide-open cellular chromatin becomes inaccessible on a global scale within hours. While viral transcripts increase drastically, the induction of the lytic phase results in a massive reduction of cellular transcripts and a loss of chromatin-chromatin interactions of cellular promoters with their distal regulatory elements as shown in Capture-C experiments. Our data document that EBV’s lytic cycle induces discrete early processes that disrupt the architecture of host cellular chromatin and repress the cellular epigenome and transcriptome likely supporting the efficient *de novo* synthesis of this herpesvirus.

## INTRODUCTION

Viruses exploit their hosts at the cellular or organismic level to support viral propagation and spread. Towards this end they also manipulate the infected cellular host with their own toolkit to suppress various antiviral defense mechanisms. For example, viruses can inhibit several levels of interferon responses (Qian et al., 2017 and references therein), they counteract cellular cytidine deaminases with potent antiviral activities (Zheng et al., 2012), reduce cellular immunity with viral micro RNAs (Albanese et al., 2017), or even mimic histone tails to interfere with antiviral responses of the infected cells (Marazzi et al., 2012).

The manipulation of the host’s antiviral programs is especially important for herpes viruses. Commonly, they turn the infected cell into a virus factory, but they can also initiate their temporal coexistence in certain cells to establish long-lasting, non-productive, latent infections. During latency, the herpesviral genomic DNA acquires a genuine cellular and highly repressive chromatin signature, which blocks transcription of most viral genes. Herpes viruses can escape from the latent phase and reactivate virus production. In this step, herpes viruses instruct their cellular host to remove the repressive epigenetic signature from viral chromatin to allow and activate massive viral transcription of all lytic viral genes within hours. During lytic reactivation, the host cell must also be manipulated to provide chemical energy, macromolecules, and nuclear space for the so-called viral replication compartments (Quinlan et al., 1984; Monier et al., 2000) or amplification factories (Chiu et al., 2013). Additionally, the cell’s transcription machinery needs to be redirected to support an efficient virus *de novo* transcription within a few hours after reactivation. Many molecular details of these fundamental processes are unknown.

A ubiquitous human herpes virus is Epstein-Barr virus (EBV, HHV4), which infects about 95 % of the human population. B lymphocytes are the preferred target cells in which EBV establishes a latent infection initially. EBV reactivates from this latent state with the help of the viral BZLF1 protein, which is expressed upon differentiation of EBV-infected memory B cells to plasma cells (Laichalk and Thorley-Lawson, 2005) and induces the switch from latency to EBV’s lytic phase (Countryman and Miller, 1985; Takada et al., 1986). BZLF1, which is also called EB1, ZEBRA, zta, or Z, is known to act as an essential viral transcription and replication factor (Takada et al., 1986; Countryman and Miller, 1985; Schepers et al., 1993). BZLF1 binds two classes of DNA binding motifs (ZREs) in the viral genome, one of which needs to be methylated to be bound efficiently (Bergbauer et al., 2010; Bhende et al., 2004). These sites are termed CpG-ZREs or meZREs and are mainly positioned in promoters of important lytic viral genes (Bergbauer et al., 2010; Kalla et al., 2010). Upon initial infection during EBV’s pre-latent phase (Jochum et al., 2012), BZLF1 is transiently expressed (Wen et al., 2007; Kalla et al., 2010), but it does not bind its many viral CpG-ZREs, because the incoming viral DNA is free of methylated CpG dinucleotides (Kalla et al., 2012). As a consequence, the virus cannot activate its lytic promoters in the pre-latent phase but it rather initiates its latent program, which leads to the restricted expression of the few latent viral genes, only. CpG methylation of viral DNA is a slow process in newly infected primary B cells that takes several weeks to completion (Woellmer et al., 2012), but CpG methylation further supports the epigenetic silencing of all viral lytic genes (Kenney and Mertz, 2014). It is likely that EBV uses this strategy to prevent the premature expression of its lytic genes in newly infected cells, which would induce a massive antiviral immune response of EBV-specific T cells and would readily eliminate the virally infected cells (Kalla and Hammerschmidt, 2012).

BZLF1 is a homo-dimeric bZIP transcription factor. Its DNA binding domain shows a strong homology to the cellular AP-1 protein family including the Jun, Fos, Fra, and ATF subfamilies. AP-1 binding motifs and ZREs are related sequences of about seven nucleotides (Lieberman et al., 1990; Chang et al., 1990). Similar to BZLF1, c-Jun/c-Fos heterodimers can also bind to and regulate cellular genes via methylated DNA motifs, which are related to CpG-ZRE motifs of BZLF1 (Gustems et al., 2014). Collectively, these findings suggest that BZLF1 may widely influence cellular transcription since several hundred thousand motifs of members of the AP-1 family are known in the genome of mammalian cells (Zhou et al., 2005), some of which are located in enhancers (Lee et al., 1987; Chavanas et al., 2008; Zanconato et al., 2015; Phanstiel et al., 2017; Vierbuchen et al., 2017). AP-1 family members participate in regulating genes involved in cellular proliferation, differentiation, and apoptosis (Shaulian and Karin, 2001; Shaulian and Karin, 2002; Ameyar et al., 2003; Eferl and Wagner, 2003), pathways which EBV also manipulates in its pre-latent, latent and lytic phases.

Much is known about the impact of BZLF1 on the viral genome after induction of EBV’s lytic phase and the ensuing alterations in viral chromatin (Schaeffner et al., 2019; Woellmer et al., 2012), but if and how herpesviruses in general and EBV in particular manipulate the chromatin of the host cell during the early hours of lytic, productive infection is not known in detail. Therefore, we examined the global modification of the host genome in the lytic, pre-replicative phase of EBV. We chose the Raji cell line as our preferred model for two reasons: (i) in Raji cells, EBV’s latent phase is very tightly controlled, but, upon induction the majority of the cells readily enter EBV’s lytic phase. The lytic phase, however, is incomplete and does not support the amplification the viral DNA because Raji cells carry a defective EBV genome (Polack et al., 1984; Hatfull et al., 1988; Decaussin et al., 1995).Thus, Raji cells (ii) allow studying the very early modifications in cellular chromatin in the pre-replicative phase when the cells transit from viral latency to lytic reactivation but prior to the formation of replication compartments or amplification factories. In EBV-infected cells they become microscopically visible as early as 16 to 24 hours after lytic phase induction and are characterized by the local accumulation of EBV DNA and the occlusion of cellular chromatin constituents such as histones (Chiu et al., 2013). This report also indicated that the nuclear architecture of the lytically induced cells undergoes substantial modifications prior to the formation of the amplification factories, because the nuclei show an initial ‘honeycombed’ structure suggestive of an early reorganization of the cellular chromatin or nuclear 3D architecture.

BZLF1 and certain members of cellular transcription factor family AP-1 are structurally and functionally similar. It seems therefore plausible that BZLF1 may take part in manipulating the chromatin and transcriptome of its cellular host during viral reactivation and the subsequent lytic phase. While our focus was on detecting global changes within the host cell upon induction of the lytic cycle we also considered the possible role of BZLF1 in inducing modifications in the host cell with respect to the cell’s transcriptome, chromatin accessibility, and chromatin-chromatin interactions.

We discovered that the induction of EBV’s lytic phase induces global alterations in the host cell chromatin. We found (i) a massive reduction of cellular transcripts indicative of the virus-induced host shut-off, (ii) a prevalent reduction or even collapse of chromatin-chromatin interactions concomitant with a (iii) general loss of chromatin accessibility, and (iv) a localized opening of chromatin at the majority of about 10^5^ cellular BZLF1 binding sites. These findings document that the host cell is subject to global nuclear changes already in the pre-replicative phase of the virus’ lytic cycle.

## MATERIAL AND METHODS

### Eukaryotic cell lines

The cell lines B95-8 (EBV-positive) (Miller and Lipman, 1973), DG75 (EBV-negative) (Ben-Bassat et al., 1977), and Raji (EBV-positive) (Pulvertaft, 1964) and their derivatives were cultured in RPMI 1640 medium (Life Technologies) supplemented with 1 mM sodium pyruvate (Life Technologies), 100 μg/ml streptomycin, 100 units/ml penicillin (Life Technologies), and 8 % fetal bovine serum (FBS) (BioSell) in an atmosphere with 5 % CO_2_, 95 % humidity at 37 °C. 1 mg/ml puromycin (Invitrogen) was added to the cells to select for the maintenance of oriP plasmids with two conditional doxycycline inducible BZLF1 alleles: iBZLF1 (p4816) and iBZLF1 (p5694) encode the wild-type BZLF1 and the activation domain (AD)-truncated alleles, respectively. On average, the cells were kept at a density of 5×10^5^ cells/ml. HEK 293 cells (Graham et al., 1977) were kept at a confluency of about 70 %. HEK 293 cells that produce the EBV strain wt/B95.8 (2089) upon induction were cultivated with 100 ng/ml hygromycin as described (Delecluse et al., 1998).

### Plasmids

The doxycycline inducible BZLF1 expressing plasmids p4816 and p5694, which encode full-length iBZLF1 and the activation domain (AD)-truncated iBZLF1, respectively, are described in Woellmer et al. (Woellmer et al., 2012). The two plasmids also express the green fluorescent protein (GFP) upon doxycycline addition (Bornkamm et al., 2005). The BZLF1 plasmid p5406 expresses full-length BZLF1 with an amino-terminal tandem step-tag as described (Schäffner, 2015).

### Quantitative BZLF1 expression analysis

BZLF1 protein was purified from HEK 293 cells transiently transfected with the BZLF1-step-tag expressing plasmid p5406 for 48 h. Cells were lysed in RIPA buffer (1 % NP-40, 0.1 % SDS, 0.5 % sodium deoxycholate, 150 mM NaCl, 50 mM Tris-HCl pH 8.0, 1 × proteinase inhibitor cocktail (Roche)), incubated (20 min), and sheared on ice with a Branson sonifier 250-D (10 sec on/ 50 sec off, 1 min, 20 % amplitude). The lysates were centrifuged (15 min, 16,000 g, 4 °C) and the supernatants were purified with IBA Strep-Tactin sepharose beads (IBA, 2-1201-002) on Poly-Prep Chromatography Columns (731-1550, BioRad) with IBA’s buffer’s W (washing) and E (elution). The concentration of purified protein was determined by Coomassie staining of 14 % gels after SDS-PAGE with BSA standards obtained from the Pierce BCA protein assay kit (23209, Thermo Fisher Scientific). The purified BZLF1 protein was used as a quantitative standard and reference for subsequent Western blot analyses.

For BZLF1 expression level analysis 2.5×10^6^ DG75, B95-8 cells, non-induced Raji iBZLF1 cells, or Raji iBZLF1 cells induced for 6 h were lysed in RIPA buffer and sheared with the BioRuptor (Diagenode) four times (5 min, 30 on/off, high) in ice cold water. 5x Laemmli loading buffer containing 15 % DTT (1 M) and 0.75 % beta-mercaptoethanol were added to the lysates. Aliquots of these lysates corresponding to different cell numbers were loaded onto 14 % SDS-PAGE gels and analyzed by Western blot immune detection. BZLF1 was detected with the BZ1 antibody (Young et al., 1991), which was used at a 1:50 dilution in combination with a secondary anti-mouse HRP antibody (1:10,000, Cell Signaling #7076S). ECL Select (GE Healthcare) was used for signal detection, which were quantified with a Fusion FX (VILBER) system. Calculation and visualization of BZLF1 dimers was done with R (Team, 2018).

### Intracellular BZLF1 staining

For the intracellular staining of BZLF1 Fix & Perm permeabilization Kit (Invitrogen) was used. 1×10^6^ cells were centrifuged (300 g, 10 min) and washed with 100 μl PBS. 100 μl of Kit reagent A were added (15 min, in the dark). Samples were washed with 2 ml wash buffer (PBS, 50 % FBS, 0.1 % NaN_3_) before 100 μl Kit reagent B was added. 1 μl BZ1 antibody coupled with Alexa647 was added before vortexing (2 sec) and incubation (20 min, RT, in the dark). Samples were washed with 2 ml wash buffer and taken up in 0.5 ml wash buffer afterwards for FACS analysis (BD Fortessa).

### Virus titration

The 2089.2.22.7 version of HEK 293 cells stably transduced with the maxi-EBV plasmid p2089 releases the EBV strain wt/B95.8 (2089) upon BZLF1 expression (Delecluse et al., 1998). This cell line was transfected with the inducible iBZLF1 expression plasmid p4816 and selected with 500 ng/ml puromycin. 4×10^5^ HEK 293 2089 cells were induced with 25, 100, or 200 ng doxycycline/ml for three days or the cells were left uninduced as a control. The supernatants were filtered through a filter with a pore size of 1.2 μm and 25, 50, or 100 μl were used to infect 10^5^ parental Raji cells. GFP positive Raji cells (Green Raji units) were identified via FACS analysis as described in detail (Steinbrück et al., 2015). The virus concentrations were calculated and visualized with R.

### Next generation ChIP-sequencing

For immunoprecipitation two samples with 1×10^8^ cells each were adjusted to a concentration of 5×10^5^ cells/ml in fresh medium and were left non-induced or were induced with a final concentration of 100 ng/ml doxycycline (Sigma-Aldrich) for 15 h. Nuclei were extracted with hypotonic buffer (10 mM KCl, 340 mM Sucrose, 1.5 mM MgCl2, 10 mM Hepes pH 7.9) containing 10 % proteinase inhibitor cocktail (PIC, Roche) and lysed in RIPA buffer + 1x PIC. The chromatin was shared in a BioRuptor on wet ice (4 cycles, 5 min each, 30 on/off, high). The chromatin was immunoprecipitated with the BZ1 antibody (Young et al., 1991), 10 % input was used as a control. The precipitates were washed with different salt concentrations and the proteins were digested with proteinase K. For Raji cells, the library preparations and sequencing (Illumina, paired-end, 150 nt) were performed by Vertis Biotechnology AG (Freising, Germany). The results were mapped with bowtie2 2.2.6 (Langmead and Salzberg, 2012), formats were transformed with samtools 1.0 (Li et al., 2009) and bedtools v2.25.0 (Quinlan and Hall, 2010) and the peak calling of the samples versus input was done by MACS2 2.1.0 (Feng et al., 2012; Zhang et al., 2008). The overlapping peaks of two replicates were merged with bedtools intersect and the DREME algorithm (Bailey, 2011) and the MEME-Chip 4.10.1 suit (Bailey et al., 2015) were used for motif identification. Raji 15 h DREME motifs (Fig. 2A) were merged artificially. The original, individual motifs and sub-motifs are shown in Supplementary Figure S3B.

### ATAC-sequencing

10^6^ Raji cells with either inducible BZLF1 plasmid (iBZLF1, 4816) or AD-truncated BZLF1 (5694) were treated with 100 ng/ml doxycycline for 15 h or left untreated. The cells were FACS sorted for living cells (untreated) and living, induced cells (15 h induced), controlled with trypan blue, and prepared for NGS. Omni-ATAC was performed as previously described (Corces et al., 2017) with minor modifications. Briefly, 50,000 FACS sorted cells were washed in PBS, resuspended in 50 μl of ATAC-seq resuspension buffer (RSB: 10□mM Tris-HCl, pH 7.4, 10□mM NaCl, and 3□mM MgCl_2_) containing 0.1% NP40, 0.1% Tween-20 and 0.01% digitonin (Promega) and were incubated on ice for 3□min. Following lysis, 1□ml of ATAC-seq RSB containing 0.1% Tween-20 was added, and nuclei were collected at 500 g (4 °C, 10□min). Pelleted nuclei were resuspended in 50□μl of transposition mix (25□μl 2□×□TD buffer, 2.5□μl Tagment DNA enzyme (Illumina Nextera DNA Library Preparation Kit, cat. FC-121-1030), 16.5□μl PBS, 0.5□μl 1 % digitonin, 0.5□μl 10 % Tween-20, and 5.25□μl water) and incubated at 37□°C for 30□min in a thermomixer shaking at 1,000 rpm. DNA was purified using Qiagen PCR clean-up MinElute kit (Qiagen). The transposed DNA was subsequently amplified in 50 μl reactions with custom primers as described (Buenrostro et al., 2013). After four cycles libraries were then monitored with qPCR: 5 μl PCR sample in a 15□μl reaction with the same primers. qPCR output was monitored for the ΔRN; 0.25 ΔRN cycle number was used to estimate the number of additional cycles of the PCR reaction needed for the remaining PCR samples. Amplified libraries were purified with the Qiagen PCR clean-up MinElute kit (Qiagen) and size selected for fragments less than 600 bp using the Agencourt AMPure XP beads (Beckman Coulter). Libraries were quality controlled by Qubit and Agilent DNA Bioanalyzer analysis. High throughput sequencing was performed by the Laboratory for Functional Genome Analysis (LAFUGA) of the Ludwig-Maximilian-University, Munich on an Illumina Hi-Seq 1500 using 50□nt single-end reads.

The data were mapped on the hg19 genome with Bowtie 1.1.2. The HOMER 4.9 software was used to calculate tag directories. The data for the metaplots and heatmaps were calculated with HOMER’s annotatePeaks.pl tool on data from three independent experiments and visualized with R (3.5.1) (Team, 2018). For heatmaps visualization the data.table (https://cran.r-project.org/web/packages/data.table/index.html) and superheat packages (Barter and Yu, 2017) were used.

### Next generation RNA-sequencing

For RNA sequencing parental Raji cells and their derivates carrying the conditional expression plasmids encoding full-length iBZLF1 or AD-truncated iBZLF1 were employed. At an initial cell concentration of 5×10^5^ cells/ml, a total of 2×10^7^ non-induced cells or 4×10^7^ cells induced with 100 ng/ml doxycycline for 6 h were analyzed. Doxycycline leads to the co-expression of BZLF1 and the truncated human NGF-receptor. To limit the analysis to BZLF1 expressing cells, only, we sorted NGF-R-positive cells with magnetic beads (MACS, Miltenyi Biotec) and the primary anti-NGF-R antibody (HB8737-1, IgG1) and a secondary anti-mouse IgG (1:10, Miltenyi Biotech) antibody. 7.5×10^5^ cells were lysed in 1 ml Trizol (Thermo Fisher Scientific), snap frozen in liquid nitrogen and stored at −80 °C.

Prior to RNA isolation identical molar amounts of ERCC spike-in control RNAs (Ambion) were added to 3×10^5^ non-induced or doxycycline induced Raji iBZLF1 (0h/6h) cells. The subsequent processes and steps were identical for all cells. RNA was extracted with the Direct-zol RNA MiniPrep Kit (Zymo Reseach). The RNA concentration and quality were controlled with Nanodrop (Thermo Fisher Scientific), Quibit (Thermo Fisher Scientific), and Bioanalyzer (Agilent), and the RNAs were treated with dsDNase (Thermo Fisher Scientific). The Encore Complete RNA-Seq Library System kit (NuGEN), which uses not-so-random hexamer primers for rRNA depletion, was used for library preparation. The samples were sequenced on an Illumina HiSeq 1500 instrument (100 nt, single-end reads).

For bioinformatic analysis the samples were mapped with Tophat2 (Kim et al., 2013) on a Galaxy server (Afgan et al., 2016) to the Hg19 genome, locally with HiSat2 2.0.1 to EBV-Raji (KF717093), and reads were assigned to annotated genes with HTSeq-count 0.6.1p1 (Anders et al., 2015). The counts were compared between induced (6 h) and non-induced (0 h) cells with the R package DESeq2 1.12.3 (Love et al., 2014). Samples with added ERCC spike-in RNAs were normalized to these. The visualization was done with R including the packages ‘Extrafont’ 0.17 (Chang, 2014) and ‘RColorBrewer’ 1.1-2 (Neuwirth, 2014).

### Promoter analysis

Bedtools intersect was used to identify peaks in the data sets, which are located within the promoters (−1kb/+5kb relative to TSS (UCSC, RefSeq Genes, Hg19)) of regulated and non-regulated genes. R was used to remove duplicates, calculate and visualize the number of BZLF1 peaks or motifs within the promoters. Genes with fewer than 20 reads were excluded from the analysis.

### Capture-C

For Capture-C analysis, 10^7^ uninduced Raji iBZLF1 cells or cells induced for different time periods (6 h, 15 h) with 100 ng/ml doxycycline were used. Cells were washed (PBS), filtered for single cells, and fixed (1% formaldehyde, 10 % fetal bovine serum in PBS). The fixation was quenched on ice with 1 M glycine before cells were homogenized two times with a 20G syringe in 3C lysis buffer (10 mM TrisHCl (pH 7.5), 10 mM NaCl, 0.2 % NP-40 in ddH2O + PIC). Cells were washed and taken up in 1.2x CutSmart buffer (NEB). SDS was added to a concentration of 0.1 % and the cells were incubated (65 °C, 40 min) while shaking (1,200 rpm). Triton X-100 was added to a final concentration of 4 % and the cells were incubated (37 °C, 1 h, 1,200 rpm). The cell suspension was divided into six aliquots of 200 μl each and incubated with 80 units DpnII at 37 °C while shaking (1,200 rpm) overnight followed by another 400 units of DpnII for 4 h. The reaction was stopped with SDS at a final concentration of 0.1 %. Samples were joined and diluted with 1.25x 3C ligation buffer (62.5 mM Tris-HCl (pH 7.5), 12.5 mM MgCl2, 1.25 mM ATP, 12.5 mM DTT in ddH_2_O) and Triton X-100 (37 °C, 1 h, shaking ever 10 min) to a final volume of 1.5 ml. 100 units T4 DNA Ligase (Affymetrix) was used for DNA fragment ligation (16 °C for 4.5 h and RT for 45 min) followed by proteinase K treatment at 65 °C for 1 h to revert crosslinking. RNAs were degraded with RNase A (300 μg, 37 °C, 1 h). DNA was extracted with organic solvents (2/3 v/v phenol-chloroform and 1/3 v/v butanol), precipitated (EtOH, 100 %), washed (EtOH 70 %), and rehydrated (TE buffer).

1.5 μg DNA was sheared with a Covaris M-series instrument (Covaris, peak incident power: 50 watts; duty factor 20 %; cycles/burst: 200 counts, duration 200 sec). Samples were cleaned with Agencourt AMPure beads (Beckman Coulter). The library preparation and sequence capture were done as described in the SureSelectXT2 Target Enrichment System for Illumina Paired-End Multiplexed Sequencing manual (Agilent, Version: E1, June 2015). The desired fragments were captured with Dynabeads (Thermo Fisher Scientific). Finally, the samples were sequenced on a HiSeq 1500 (Illumina, paired-end reads, 100 nt read length).

Overlapping paired-end reads were joined using the flash tool (Magoc and Salzberg, 2011). The sequences were DpnII digested in silico with the dpnII2E.pl script (kindly provided by James Davies, Oxford, UK) and mapped with Bowtie 1.1.0 (Langmead et al., 2009). The perl script dpngenome3_1.pl (kindly provided by James Davies, Oxford, UK) was used to digest the hg19 genome in silico with DpnII. Interactions between distant DNA fragments were identified with the CCAnalyser2.pl script (kindly provided by James Davies, Oxford, UK). The data were averaged between triplicates, normalized by the total number of interactions for all captured fragments per time point, and visualized with R. Additionally, ChIP-seq and RNA-seq results were added to the visualization.

## RESULTS

The aim of this study was to investigate the alterations of the chromatin structure and the transcriptome of latently infected B cells when BZLF1 is expressed and EBV’s lytic cycle starts. First, we determined the physiological levels of BZLF1 protein in cells that naturally support virus *de novo* synthesis. Next, we stably introduced two conditional oriP plasmids into Raji cells. The two plasmids express regulated levels of full-length BZLF1 or BZLF1 devoid of its transactivation domain upon addition of doxycycline. The established human B cell line Raji is latently infected with EBV, does not support EBV’s lytic phase spontaneously but enters it rapidly and synchronously upon the induced expression of full-length BZLF1. We used this cell line (and parental controls) throughout our studies and investigated the cells in their non-induced state and upon addition of doxycycline for up to 15 hours covering the pre-replicative lytic phase of reactivated EBV. Indistinguishable from full-length BZLF1, BZLF1 without its transactivation domain binds to its many cognate sequence motifs, but cells that express it do not enter EBV’s lytic phase (Woellmer et al., 2012; Schaeffner et al., 2019). We therefore used Raji cells engineered to express the conditional AD-truncated BZLF1 for certain control experiments.

### Lytic viral gene expression requires high levels of BZLF1

When latently EBV infected memory B cells come in contact with their cognate antigens the viral BZLF1 gene is activated and virus synthesis is induced (Laichalk and Thorley-Lawson, 2005). *In vivo*, only single memory B cells respond to this trigger, which makes it technically impossible to determine the protein levels of BZLF1 required to induce and/or support the lytic phase of EBV’s life cycle in these cells. Therefore, we turned to the B95-8 cell line because a small fraction within its population spontaneously enters the lytic phase and releases infectious virions. We first determined this fraction of cells in the lytic cycle to be 2.8 % on average by intracellular FACS staining with an Alexa 647 fluorophore coupled monoclonal antibody directed against BZLF1.

We then obtained whole cell protein lysates from a known number of B95-8 cells and compared the BZLF1 signal intensities by Western blot immunodetection with known molar amounts of a truncated BZLF1 protein that was recombinantly expressed in HEK 293 cells and purified to high homogeneity (Supplementary Figs. S1 and S2). We also engineered Raji cells to carry a conditional BZLF1 allele on the oriP-based p4816 plasmid (Woellmer et al., 2012), added doxycycline for 6 hours (Raji 6h iBZLF1), and applied protein lysates of non-induced Raji cells (Raji 0h iBZLF1) to load onto the same gel. Depending on the cell line and its state of induction protein equivalents that corresponded to different cell numbers were loaded and quantified. Supplementary Figure 1A provides three independent replicates. Based on these considerations, we estimated that each B95-8 cell that was found to express BZLF1 by intracellular FACS staining contained 1.3×10^6^ ±0.2×10^6^ BZLF1 dimers on average. BZLF1 levels in Raji cells induced for 6 h with doxycycline were about 3.5-fold higher (Supplementary Fig. 1B) while the population of non-induced Raji cells still expressed detectable levels of BZLF1 that were lower by a factor of about 42 (Supplementary Fig. 1A, B).

We were concerned that the conditional expression plasmid p4816 is leaky and might express steady state levels of BZLF1 even in the absence of doxycycline that are sufficient to induce EBV’s lytic phase in latently infected cells. Since Raji cells are incapable of fully supporting *de novo* virus synthesis (Polack et al., 1984) we turned to a derivative of HEK 293 2089 cells that releases the EBV strain wt/B95.8 (2089) upon BZLF1 expression (Delecluse et al., 1998). We stably introduced the p4816 plasmid into these cells (HEK 293 2089 iBZLF1) and evaluated the concentration of EBV virions in the cells’ supernatant in the absence and presence of different concentrations of doxycycline (Supplementary Fig. 1C). The supernatants were used to infect Raji cells that turn GFP-positive upon infection with the EBV strain wt/B95.8 (2089). Virus amounts were expressed as Green Raji units (GRU) as described (Delecluse et al., 1998; Steinbrück et al., 2015). As expected, only HEK 293 2089 iBZLF1 cells released infectious wt/B95.8 (2089) EBV in a dose-dependent fashion after adding increasing concentrations of doxycycline but not the parental HEK 293 2089 cells (Supplementary Fig. 1C). Without doxycycline the supernatants of HEK 293 2089 iBZLF1 cells contained very low numbers of virus similar to supernatants obtained from the parental HEK 293 2089 cells (Supplementary Fig. 1C). In all subsequent experiments a doxycycline concentration of 100 ng/ml was used in cells with the conditional expression plasmid p4816 (iBZLF1) to reach BZLF1 levels comparable to lytically induced B95-8 cells (Supplementary Fig. 1A, B).

In RNA-seq experiments using our conditional Raji iBZLF1 cell model, we compared viral lytic genes prior to and after induction of BZLF1 to test how the expression of BZLF1 influences their expression. Before induction, lytic genes were expressed only at very low levels, while latent genes were strongly expressed (Supplementary Tab. S1). Upon adding doxycycline for 6 hours, Raji iBZLF1 cells readily expressed many viral genes that clearly mark the onset of EBV’s lytic phase. Figure 1 shows the log2 fold differential expression of viral genes comparing non-induced and induced Raji iBZLF1 cells after ERCC spike-in normalization (see below for experimental details). Very few viral genes were downregulated such as EBNA1 and EBNA2 among others latent genes (Supplementary Tab. S1). Five of the six early lytic genes known to be essential for lytic amplification of viral DNA (Fixman et al., 1992) were upregulated 60 to more than 250-fold: BBLF3-BBLF2, BBLF4, BALF5, BMLF1-BSLF1, and BMRF1 (highlighted genes in Fig. 1). BALF2, which is the sixth gene on this list and know to be deleted in Raji EBV DNA (Decaussin et al., 1995). Additional genes that contribute to the lytic replication of viral DNA were strongly induced: BKRF3, BRLF1, BHLF1, and BMLF1 as well as BZLF1 (Fixman et al., 1995) (Fig. 1). The most strongly upregulated gene, BDLF3.5, was induced almost 300-fold. Supplementary Table S1 provides a list of viral genes and their regulation.

**Figure 1.**
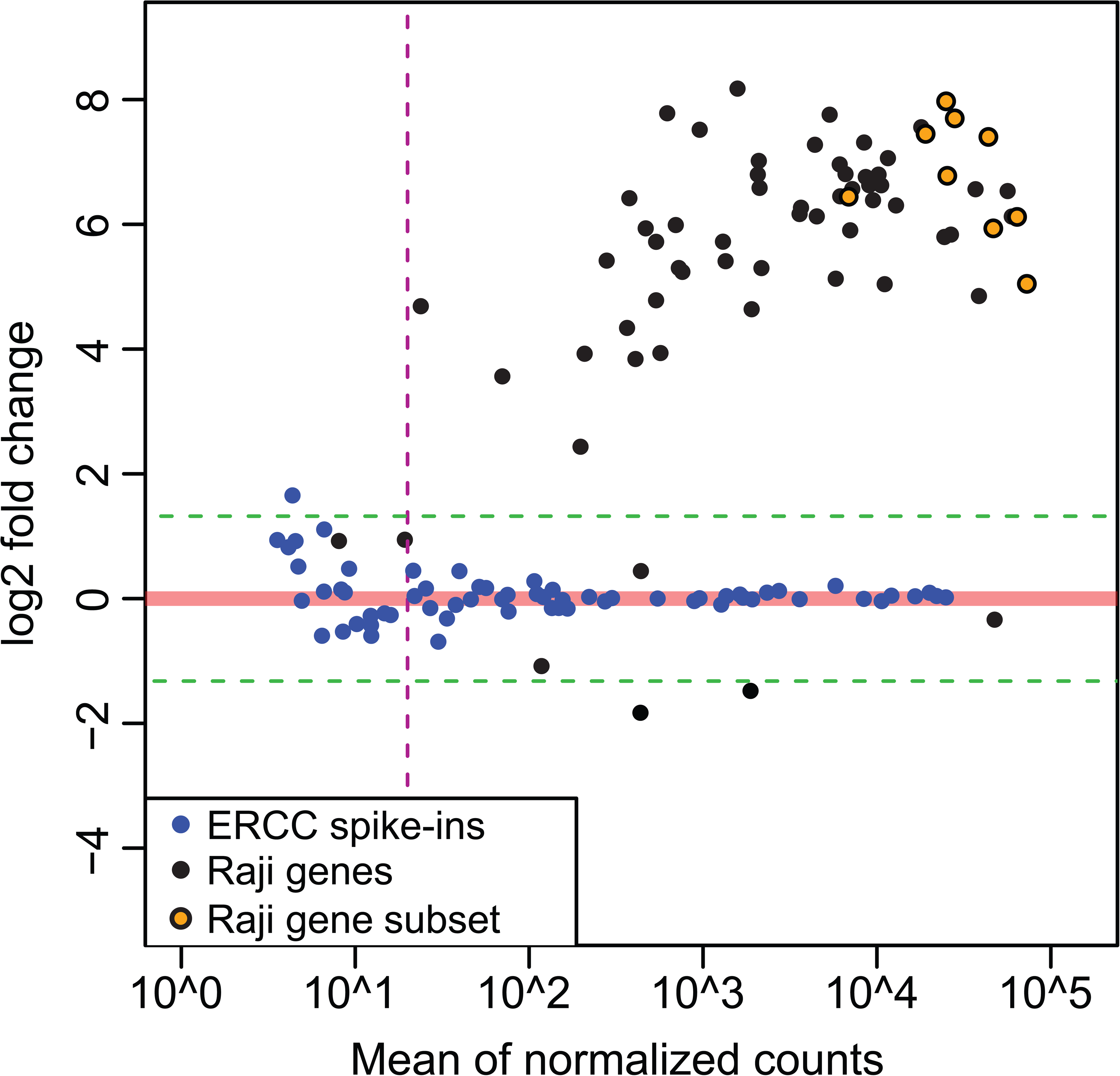
Expression of lytic viral genes in Raji cells upon induced expression of BZLF1. The induction of BZLF1 in Raji iBZLF1 cells for 6 hours resulted in the regulation of viral genes in the range of 0.3 fold (EBNA2) to 300 fold (BDLF3.5). The x-axis shows the mean of the sequencing reads after ERCC normalization, the y-axis shows the log2-fold change of gene expression in non-induced cells compared with cells induced with doxycycline for 6 hours. The blue dots indicate the detected 63 ERCC spike-in RNA reads, which were used to normalize the data set. Only genes with more than 20 sequencing reads (indicated by the purple dashed vertical line) that were either up-regulated (61 genes, >2.5 fold, green dashed horizontal line) or down-regulated (2 genes, <0.4 fold, green dashed horizontal line) were taken into consideration. Viral genes outside this range are greyed out, while genes that are essential, important or known to support viral DNA replication are indicated as orange dots and are described in detail in the text. Supplementary Table S1 provides the complete list of BZLF1 regulated viral genes.

### BZLF1 binds a high number of DNA motifs in cellular chromatin

The binding sites and sequence motifs of BZLF1 were already identified and further analyzed in the viral genome (Bergbauer et al., 2010; Dickerson et al., 2009; Flower et al., 2011). BZLF1 is a member of the cellular AP-1 family of transcription factors (Farrell et al., 1989), which are estimated to bind about 500,000 sites in human DNA (Zhou et al., 2005). We therefore hypothesized that BZLF1 might also bind to many accessible sites in the chromatin of human B cells. To address this possibility, we used our conditional iBZLF1 Raji cell model to identify BZLF1 binding sites in a genome-wide ChIP-seq approach.

We analyzed three different experimental conditions using the BZ1 monoclonal antibody directed against the dimerization domain of BZLF1 (Young et al., 1991) in our ChIP-seq analysis: Raji iBZLF1 cells in their (i) non-induced state and (ii) after induction of full-length BZLF1 with 100 ng/ml doxycycline for 15 hours, as well as (iii) DG75 cells free of EBV as a negative control.

In the control DG75 cells we found few mappable reads after ChIP-seq that rarely accumulated to peaks. This finding suggested that our ChIP-seq protocol delivers data with a very low background, because DG75 cells do not express BZLF1.

When BZLF1 is expressed at low levels in Raji iBZLF1 cells in the absence of doxycycline (Supplementary Fig. 1) we found about 30,000 ChIP peaks, of which 70 % (Fig. 2A) contained the known BZLF1 binding motif TGWGCGA (Fig. 2C) also termed meZRE (Bergbauer et al., 2010; Woellmer et al., 2012). BZLF1 preferentially binds this motif when its CpG dinucleotide contains methylated 5’-cytosine residues (Bergbauer et al., 2010). When the cells were induced with doxycycline for 15 hours, more than half of the peaks with this motif were preserved (Fig. 2B), but the total number of peaks increased dramatically (Fig. 2A, B). Now, about 146,000 ChIP peaks were identified, of which 92% (Fig. 2A) contained one or more copies of the less precisely defined consensus motif TGWGYVA (Fig. 2C). At low BZLF1 levels this motif was not among the highly ranked motifs. Supplementary Figure S3 contains detailed information about different motif compositions, their frequencies and p-values comparing the non-induced and induced states in iBZLF1 Raji cells.

**Figure 2.**
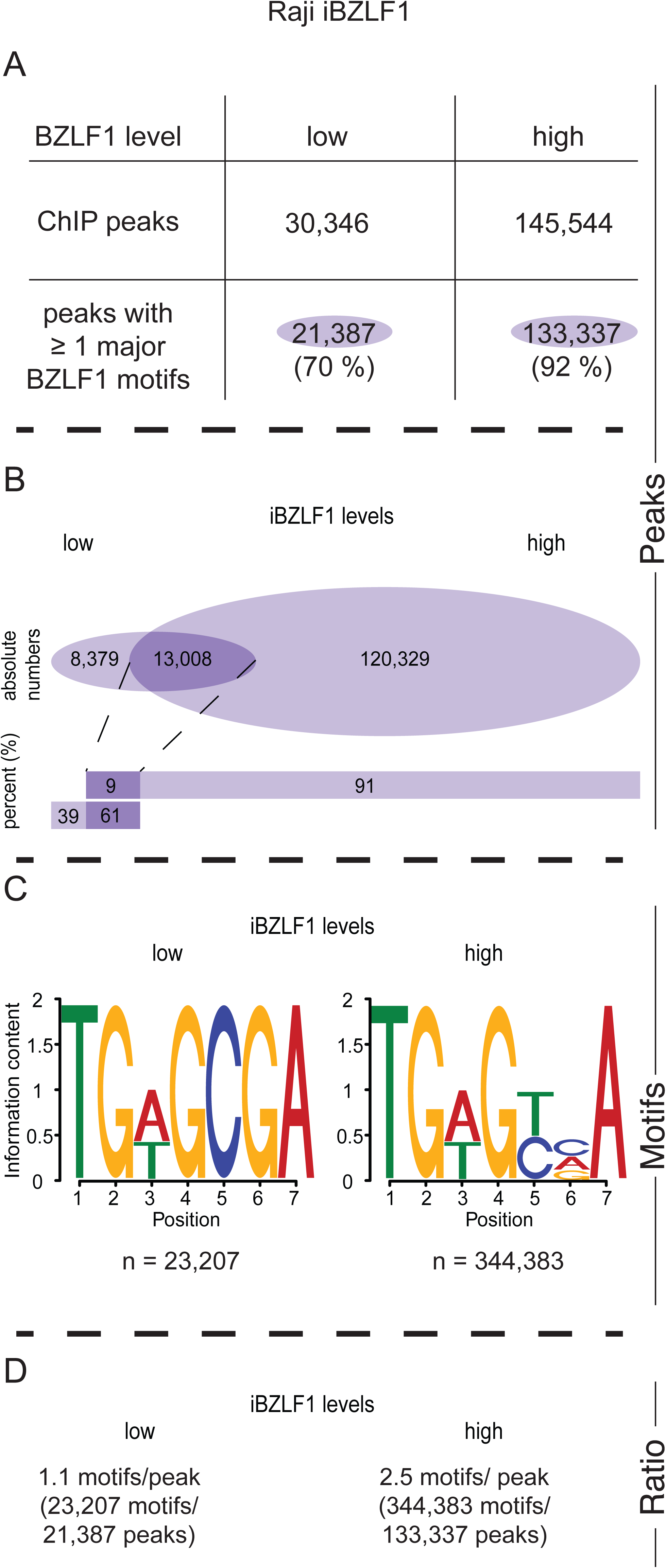
Identification of ChIP-seq peaks and BZLF1 binding motifs in chromatin of Raji iBZLF1 cells. **(A)** Peaks identified with the BZ1 antibody directed against BZLF1 at low and high levels (i.e. non-induced and induced levels, respectively) of BZLF1 in EBV-positive Raji iBZLF1 cells. **(B)** The intersections of the ‘low’ and ‘high’ peak sets with at least one identified motif indicate that the majority of low-level peaks are maintained when BZLF1 is highly induced. The abundance of peaks increases up to 5-fold at high BZLF1 levels compared with the peak number at low BZLF1 levels. **(C)** At low BZLF1 levels the known BZLF1 binding motif TGWGCGA predominates in individual peaks. At high BZLF1 levels the less specific TGWGYVA motif was identified as the major motif, which encompasses the previously identified meZRE motif TGWGCGA (Bergbauer et al., 2010). The Supplementary Figure S3 shows the detailed statistics and the relative abundance of frequent motif compositions. **(D)** The average frequencies of the number of BZLF1 motifs per ChIP-seq peak is provided.

In a minor fraction of called peaks the MEME-Chip suite (Bailey et al., 2015) failed to identify a common sequence motif for unknown reasons. We therefore turned to a visual inspection of the DNA sequences within the peak regions of this ChIP fraction. After inspection of peaks, in which no other motif could be identified previously, we found an additional and related sequence motif TGWGYVT. This motif is very similar to the meZRE motif TGWGYVA except its last nucleotide residue. We continued our analysis with a computational search for this possible motif in the sets of called ChIP peaks. Data shown in Supplementary Figure S4A indicate that 58 % of all peaks in Raji iBZLF1 cells contained the motif TGWGYVT when BZLF1 levels were high (Supplementary Fig. S4A). Taking both motifs with terminal A or T residues into consideration, 96 % of all peaks contained the consensus motif TGWGYVW at high BZLF1 levels. Detailed information about the motif can be found in Supplementary Figure S5. We also found that the majority of peaks contains both the motifs ending with A or T (Supplementary Fig. S6).

In total, 146,000 peaks in induced Raji iBZLF1 cells contained 580,000 TGWGYVW motifs of which 344,000 terminated with A and 236,00 with T (Fig. 2C, Supplementary Figs. S3, S4C, S5). On average each peak was found to contain about 2.6 BZLF1 binding motifs (Fig. 2D, Supplementary Fig. S4D). The analysis also revealed a clear hierarchy of the BZLF1 sequence motifs as shown in Figure 3. The highest affinity at low BZLF1 levels appears to correlate with the motif TGWGCGA, whereas the weakest binding is associated with the motif TGWGYVT to which BZLF1 binds also less frequently at high BZLF1 levels compared with the TGWGYVA motif (Fig. 3).

**Figure 3.**
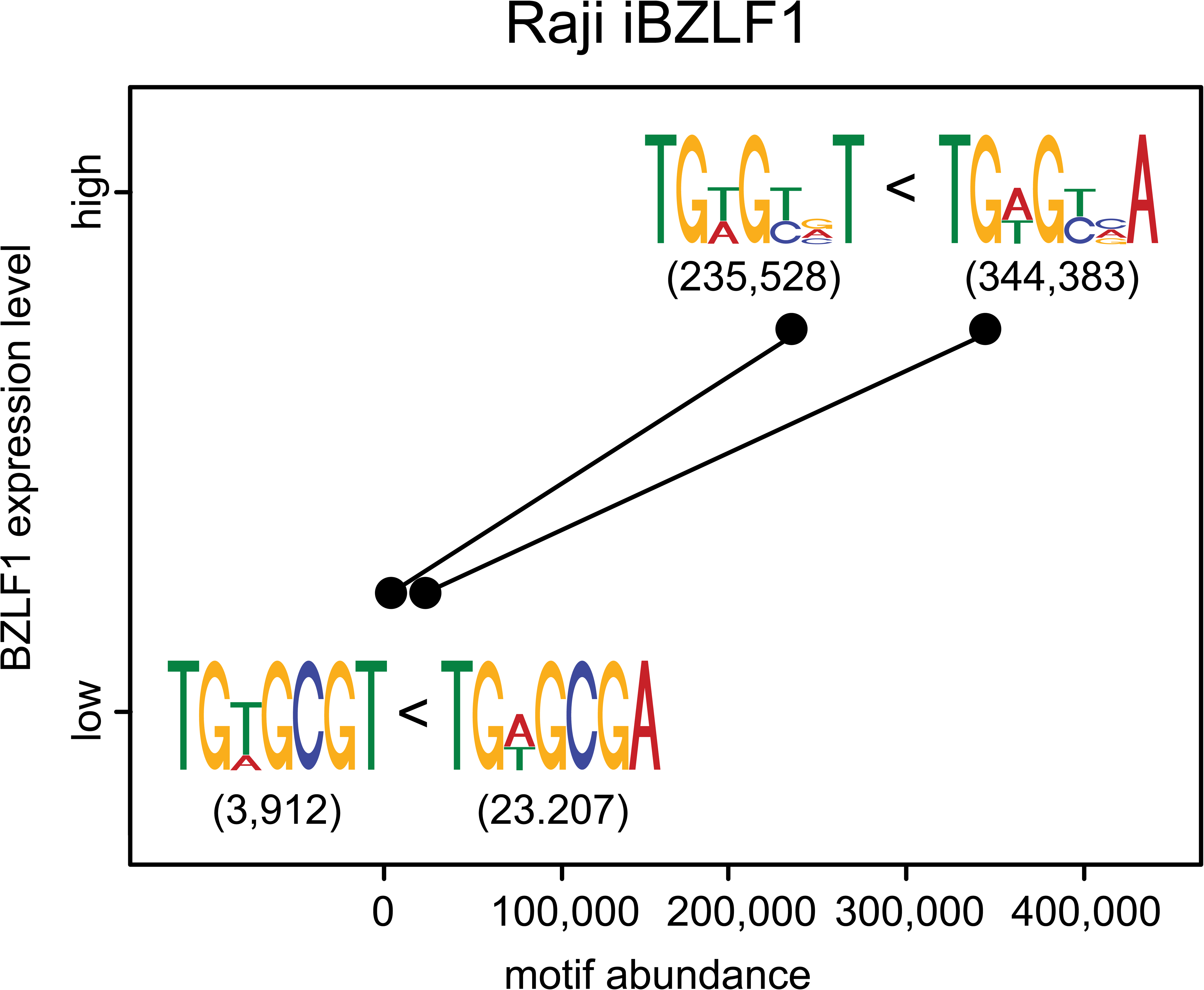
Motif abundance as a function of BZLF1 levels in Raji iBZLF1 cells. The number of the four BZLF1 binding motifs found in ChIP-seq peaks is plotted at low and high expression levels of BZLF1. At both levels the number of motifs ending with the residue A exceeds the number of motifs ending with T. The finding suggests that BZLF1’s ranked motif preference is TGWGCGA > TGWGCGT > TGWGYVA > TGWGYVT.

### High BZLF1 levels induce open chromatin at cellular BZLF1 binding sites but a genome-wide loss of open chromatin

Next, we investigated the chromatin of EBV’s host cells and studied the consequences of inducing EBV’s lytic cycle and the binding of BZLF1 to cellular DNA. From our work in press (http://biorxiv.org/cgi/content/short/317354v1) (Schaeffner et al., 2019) we knew that BZLF1 is a viral pioneer factor, binds to nucleosomal DNA, recruits chromatin remodelers such as INO80, and induces the local opening of repressed viral chromatin. To extend this insight, we analyzed the chromatin accessibility of non-induced and induced iBZLF1 Raji cells by ATAC-seq. In these experiments, Raji cells engineered to carry a conditionally expressed BZLF1 gene lacking its transactivation-domain (AD-truncated BZLF1) were used as a negative control.

We combined the ATAC-seq data with the BZLF1 peaks identified at high levels (Fig. 2A) to analyze the local chromatin accessibility as a function of BZLF1 binding. Supplementary Figure S7 illustrates the bioinformatic approach. Average chromatin accessibilities along all BZLF1 peaks are shown in Figure 4A. Raw signals and input signals in heatmap representation are shown in Supplementary Figures S8 and S9, respectively.

**Figure 4.**
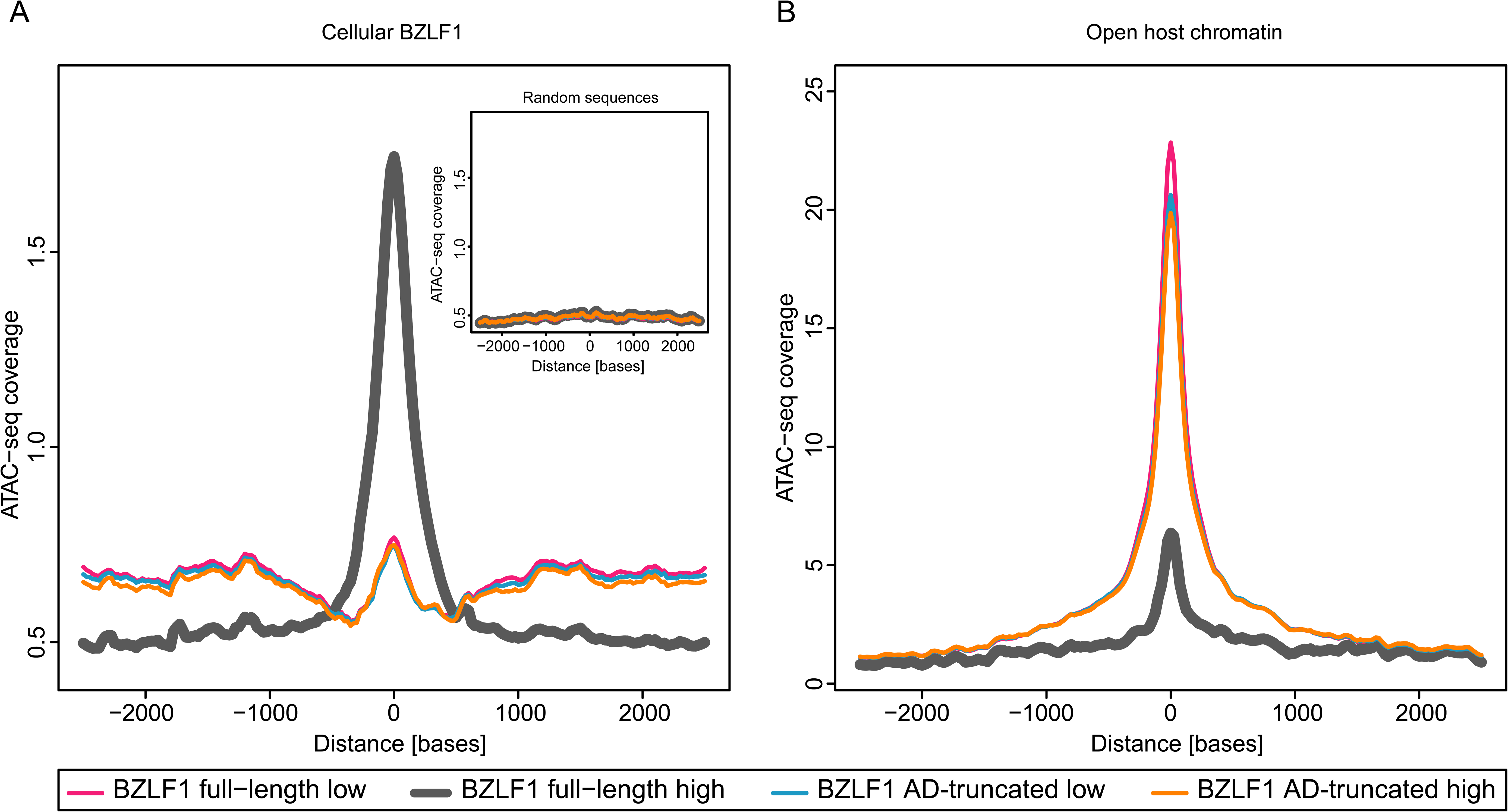
Changes in cellular chromatin accessibility after induction of EBV’s lytic cycle by BZLF1. **(A)** The metaplot summarizes the chromatin accessibility at the 145,000 BZLF1 binding sites (Fig. 2) prior to and after induction of full-length or AD-truncated BZLF1 in Raji cell chromatin. The average ATAC-seq coverages in the four different Raji cell samples are plotted according to the nucleotide coordinates of the centers of the 145,000 BZLF1 peaks. In non-induced Raji iBZLF1 cells (BZLF1 full-length, low) the average ATAC-seq coverage is congruent with the coverage found in induced and non-induced Raji cells that carry the conditional AD-truncated BZLF1 allele. At induced BZLF1 levels (full-length, high) the average ATAC-seq coverage is substantially increased indicating a gain in chromatin accessibility. The inset provides the ATAC-seq coverage of 1,145,000 randomly sampled sequences in the chromatin of Raji iBZLF1 cells expressing full-length BZLF1 at high levels after doxycycline-mediated induction. **(B)** The metaplot summarizes the ATAC-seq coverage at the 81,000 called peaks of open host chromatin identified prior to the induction of EBV’s lytic phase. After induced expression of full-length BZLF1 the chromatin accessibility is strongly reduced indicating that open host chromatin becomes globally inaccessible upon induction of EBV’s lytic phase. The ATAC-seq coverage is barely affected compared with non-induced cells when the AD-truncated BZLF1 protein is expressed. The data summarize three independent biological replicates.

We analyzed the average coverage of ATAC-seq reads at these sites in the two Raji cell lines that express full-length or the AD-truncated BZLF1 protein in their non-induced and induced states. The visualization in Figure 4A documents the opening of silent, compact chromatin at cellular BZLF1 binding sites that occurs at induced high levels of BZLF1, only. An example is shown in Supplementary Figure S10A and heatmaps are provided in Supplementary Figure S11A. The average peak of chromatin opening co-locates exactly with the peak center of the >10^5^ cellular BZLF1 binding sites in induced cells that express full-length BZLF1. A truncated BZLF1 protein without its transcriptional activation domain does not induce chromatin remodeling (Fig. 4A, Supplementary Fig. S11, panel A4), although it binds inactive chromatin as efficiently as full-length BZLF1 (Bergbauer et al., 2010). This finding clearly supports our view that BZLF1’s activation domain recruits chromatin remodelers such as INO80 to silent chromatin (http://biorxiv.org/cgi/content/short/317354v1) (Schaeffner et al., 2019). As a consequence, the previously repressed chromatin becomes accessible and is presumably activated (Fig. 4A, Supplementary Fig. S11, panel A2) or is kept open (Supplementary Fig. 10B). Accessibilities at random locations (n=1,455,440) not linked to BZLF1 binding sites were not affected (inset in Fig. 4A, Supplementary Fig. S11B).

Raji cell chromatin in the latent phase demonstrates a high number of accessible, open regions that are clearly identifiable by ATAC-seq (Supplementary Fig, S10C). We found about 81,000 peaks of accessible cellular chromatin in latent Raji iBZLF1 cells using the MACS2 peak caller and calculated the average ATAC-seq coverage of open chromatin prior to and after induction of EBV’s lytic cycle. As shown in Figure 4B, in corresponding heatmaps (Supplementary Fig. S11, panel C), and in a representative IGV browser image (Robinson et al., 2011; Thorvaldsdóttir et al., 2013) shown in Supplementary Figure S10C, we found a strong general reduction of open Raji cell chromatin upon induction of EBV’s lytic phase. Reduction of chromatin accessibility was evident at the majority of ATAC-seq peaks previously identified to be accessible during latency (Fig, 4B, Supplementary Fig. 10C).

The data indicate that the induced expression of BZLF1 together with the ensuing activation of EBV’s lytic phase modulate the accessibility of cellular chromatin globally. On the contrary, the very many open chromatin loci in latently infected Raji cells become mostly non-accessible whereas chromatin loci bound by BZLF1 open up site-specifically in the very early phase of lytic reactivation.

### The induction of EBV’s lytic cycle drastically reduces cellular transcripts in Raji cells

We investigated the consequences of the induction of EBV’s lytic phase and BZLF1’s binding to cellular chromatin with respect to transcriptional regulation. RNA-seq experiments were performed in three different experimental layouts:

(i) Steady state RNA transcript levels of parental Raji cells were analyzed and compared with the same cells incubated with doxycycline for six hours (Fig. 5A). Doxycycline did not regulate any genes according to our threshold criteria, which are provided in the figure. The violin plot shown in Figure 5A indicates that the log2 fold change of 95 % of the expressed genes, subsequently termed ‘population spread’, was located in a very narrow range in parental Raji cells (Fig. 5A, bottom panel).

**Figure 5.**
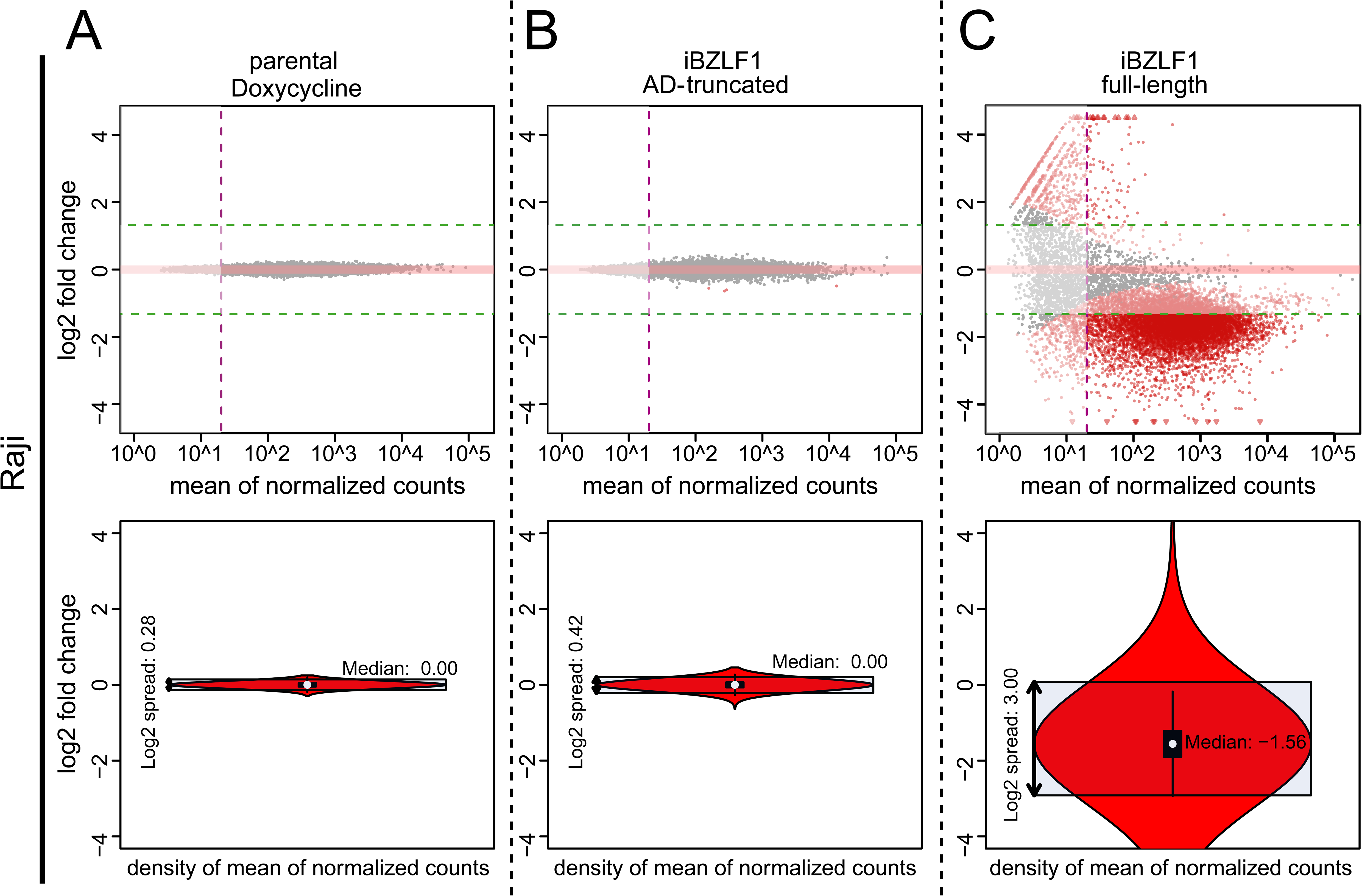
Gene regulation in Raji cells upon doxycycline-induced expression of full-length BZLF1 compared with a BZLF1 variant lacking its transcriptional activation domain. Three different cell lines (parental Raji cells [panel A], Raji iBZLF1 AD-truncated [panel B], and Raji iBZLF1 [panel C]) were induced with doxycycline for 6 hours and compared with their non-induced counterparts. The analyses are based on the hg19 reference genome and three replicates of each condition and cell line. Viral genes and *NGFR* (used as a doxycycline-regulated reporter gene) were excluded from the analyses. **(A)** The transcriptomes of parental Raji cells were analyzed by comparing their untreated versus doxycycline treated (6 hours) states. No gene with more than 20 sequencing reads (vertical purple line) was found to be up-(>2.5x) or down-regulated (<0.4x) in the MA plots (upper and lower green horizontal lines, respectively). The median is centered at zero in the violin plot (lower panel). The distance between the quantiles that encompass 95 % of all data points describes the spread of the gene populations which was determined to be 2^0^.^28^. **(B)** RNA expression in Raji cells with a conditional activation-domain (AD)-truncated BZLF1 allele upon doxycycline induction for 6 hours. No gene was considered up-or down-regulated. The violin plot reveals the very narrow spread of the gene population (2^0^.^42^) comparable to the results shown in panel A. **(C)** RNA expression in Raji cells upon doxycycline induction of full-length wild-type BZLF1 (iBZLF1) after data normalization according to ERCC spike-in RNA reads. The MA plot shows a strong global reduction of cellular mRNA transcripts 6 hours after induction of BZLF1. 91 genes were upregulated (upper horizontal green line), while transcripts of 7,174 genes were reduced by a factor of at least 0.4 (lower horizontal green line). The MA violin plot shows that the median of the gene population is reduced by a factor of almost three (2^−1^.^56^) indicating a global reduction of mRNA steady state levels. The distance between the 2.5 and 97.5 quantiles shows a spread of the gene population of 2^3^.

(ii) We repeated the experiment with Raji iBZLF1 AD-truncated cells engineered to express a truncated version of BZLF1 without its transcriptional activation domain (Woellmer et al., 2012). Upon adding doxycycline for 6 hours, no gene was identified to be regulated (Fig. 5B). The population spread in Raji iBZLF1 AD-truncated cells slightly exceeded the range found in the parental Raji cells (Fig. 5B).

(iii) Raji iBZLF1 cells engineered to express the full-length BZLF1 protein were induced by adding doxycycline for six hours (Supplementary Fig. 1) and were analyzed by RNA-seq. We used the ERCC RNA spike-in mix to be prepared for global changes in the cellular transcriptomes, to analyze the basic performance metric of the RNA-seq libraries, and to normalize the data during the steps of subsequent bioinformatic analyses according to this standard. The quantitative detection of the external spike-in RNAs demonstrated the linearity, quality, and reproducibility of the experimental approach (Supplementary Fig. S12).

In Raji iBZLF1 cells we found 91 cellular genes with increased transcript levels, while 7,174 showed a decrease (Fig. 5C). The population spread was much higher when compared to the controls and the median of the population was shifted to a strong negative value indicating that the majority of cellular transcripts was substantially reduced on average. Individual triplicates are shown as boxplots in Supplementary Figure S13.

The top 1,000 most strongly downregulated genes in Raji iBZLF1 cells were used in a KEGG pathway analysis, which identified critical metabolic pathways and pathways in immune cells involved in host cell immune responses, antigen and cytokine signaling (Supplementary Fig. S14).

The results indicate that the short-term induced expression of full-length BZLF1 and subsequent induction of EBV’s lytic cycle led to a dramatic and global drop of transcript levels in Raji iBZLF1 cells. Fewer than hundred genes were found upregulated upon BZLF1 expression, an unexpected finding, because BZLF1 is a known transcriptional activator in the viral genome which also binds to the cellular genome frequently (Fig. 2, Supplementary Fig. S4).

### BZLF1 binding sites in proximity to TSS do not correlate with the magnitude of gene regulation

Since BZLF1 binds more than 145,000 sites in the cellular chromatin of Raji iBZLF1 cells (Fig. 2, Supplementary Fig. S3), it seemed plausible that BZLF1 regulates promoters of cellular genes similar to the many viral promoters of lytic EBV genes (Bergbauer et al., 2010; Kalla et al., 2010; Woellmer et al., 2012). To address this question, we looked for peaks identified by ChIP-seq within promoter regions of genes found to be regulated in our RNA-seq experiments (Fig. 5C). We limited our search to a region 5 kb upstream to 1 kb downstream of the TSSs (Fig. 6A). About half of the upregulated genes comprised one or more peaks within the defined limits of their TTSs (Fig. 6A). In Raji iBZLF1 cells, the promoter regions of 63 % downregulated genes did not contain recognizable peaks attributed to BZLF1 binding (Fig. 6A).

**Figure 6.**
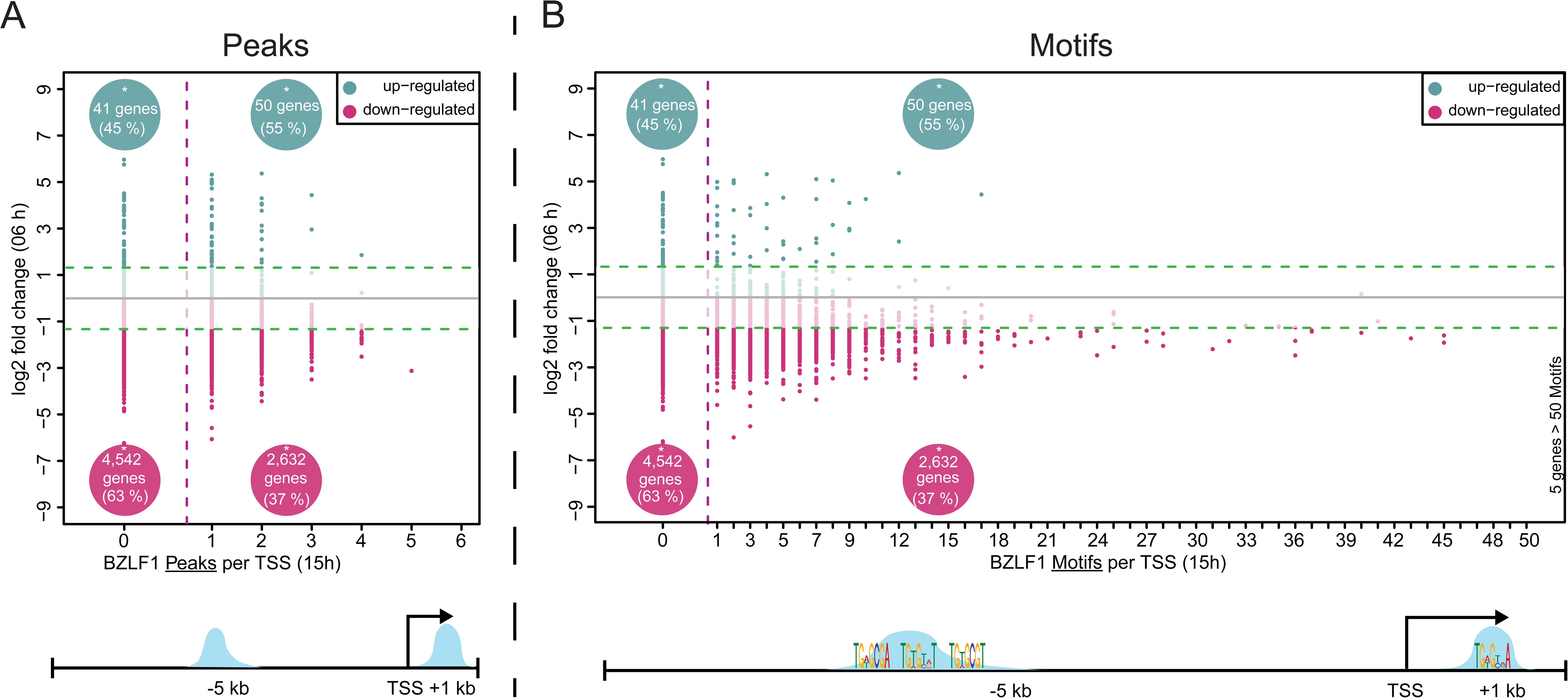
Regulated genes and their association with BZLF1 ChIP-seq peaks and binding motifs in promoter regions. **(A)** The numbers of ChIP-seq defined peaks are plotted on the x-axis versus the magnitude of gene regulation expressed as ‘log2 fold change’ on the y-axis after doxycycline induced expression of full-length BZLF1 in Raji iBZLF1 cells. The analysis includes the promoter proximal region −5 kb/ +1 kb of the transcriptional start sites (TSS) as indicated in the cartoon below. Forty-one genes without BZLF1 peaks within their defined promoter region were upregulated whereas about 4,500 gene were downregulated (upper and lower green lines, respectively). Fifty and about 2,600 genes that had BZLF1 peaks in their promoters were found to be up-or downregulated, respectively. **(B)** The number of BZLF1 motifs as defined in panel C of Figure 2 and Supplementary Figure S4 panel C are plotted on the x-axis versus the magnitude of gene regulation on the y-axis as in panel A. BZLF1 motifs downstream and upstream of the TSS entered the analysis. The distribution of regulated genes (y-axis) with or without BZLF1 binding sites follows the scheme in panel A. Below, the cartoon depicts one peak upstream and downstream of TSS with three and one BZLF1 motifs, respectively, illustrating the basics of this analysis and the location of the motifs.

We also searched for a possible correlation between the absolute numbers of BZLF1 motifs within the promoter regions and the magnitude of gene regulation. Similar to the results shown in Figure 6A, we did not find an obvious correlation. Certain promoters with few BZLF1 binding motifs were more profoundly regulated than promoters with multiple binding motifs (Fig. 6B). Few cellular genes seemed to be exceptions to this rule in Figure 6B. A very small number of genes contained more than 50 and up to 142 BZLF1 binding motifs within their promoter regions but they were barely regulated upon BZLF1 expression.

Together, the many identified BZLF1 binding motifs in promoter regions of cellular genes did not provide a clear function that could be ascribed to BZLF1. BZLF1 was characterized as a transcription factor of viral promoters, initially, but BZLF1 has also been proposed to act as an enhancer factor (Ramasubramanyan et al., 2015) similar to its cellular homolog AP-1 (Lee et al., 1987; Chavanas et al., 2008). The cellular genome is about eighteen thousand times larger than the EBV genome, contains up to 26,000 genes and an estimated number of up to 9,000,000 regulatory regions (International, 2004; Nguyen et al., 2018). We asked if, upon induction of the lytic phase, EBV governs these regulatory regions at the level of chromatin interactions. We also asked if the frequent binding of BZLF1 to cellular DNA together with the remodeling of silent chromatin may have a discernable effect. Towards this end we selected several up-, down-, and non-regulated cellular genes to look at their interacting chromatin regions, which BZLF1 might regulate to control the expression of cellular genes.

### Induction of EBV’s lytic cycle reduces cellular chromatin interactions

The regulation of cellular transcription results from an interplay between transcription factors, histone and DNA marks, and chromatin interactions. The latter govern the spatial proximity between enhancers (or repressors) and promoter regions and regulate the activity of genes. The likelihood of functional interactions increases when distant regions are organized into loops by chromatin organizing proteins such as CTCF and cohesin.

We applied the Capture-C technique (Hughes et al., 2014) to study physical chromatin interactions of selected promoters as a function of BZLF1 levels. From our RNA-seq experiment with Raji iBZLF1 cells (Fig. 5C) we chose promoters of genes that were up-or down-regulated but we also included genes that were not affected when BZLF1 was induced. With synthetic single-stranded RNA probes that hybridize to regions of approximately 5 kb up-and downstream of the transcriptional start sites (TSS) we focused on 53 selected genes and pulled-down their promoter regions.

After data normalization we identified about 1.3 million interactions between the pulled regions and proximal or distal flanking DNA fragments that had been generated by DpnII cleavage following the Capture-C protocol. For about 170,000 DpnII fragments a fold change could be calculated in a pair-wise comparison of non-induced Raji iBZLF1 cells with cells induced for 6 or 15 hours (0 versus 6 h; 0 versus 15 h). Figure 7A shows a slight increase of total interactions 6 h after induction of EBV’s lytic cycle. About 1,000 DpnII fragments showed a 2-fold increase of the number of interactions, whereas about 300 DpnII fragments lost more than half of their interactions. After 15 h the total number of interactions clearly decreased (Fig. 7B). About 4,000 DpnII fragments pairs had lost more than half of their interactions while only 74 DpnII fragment pairs showed a more than 2-fold gain of interactions.

**Figure 7.**
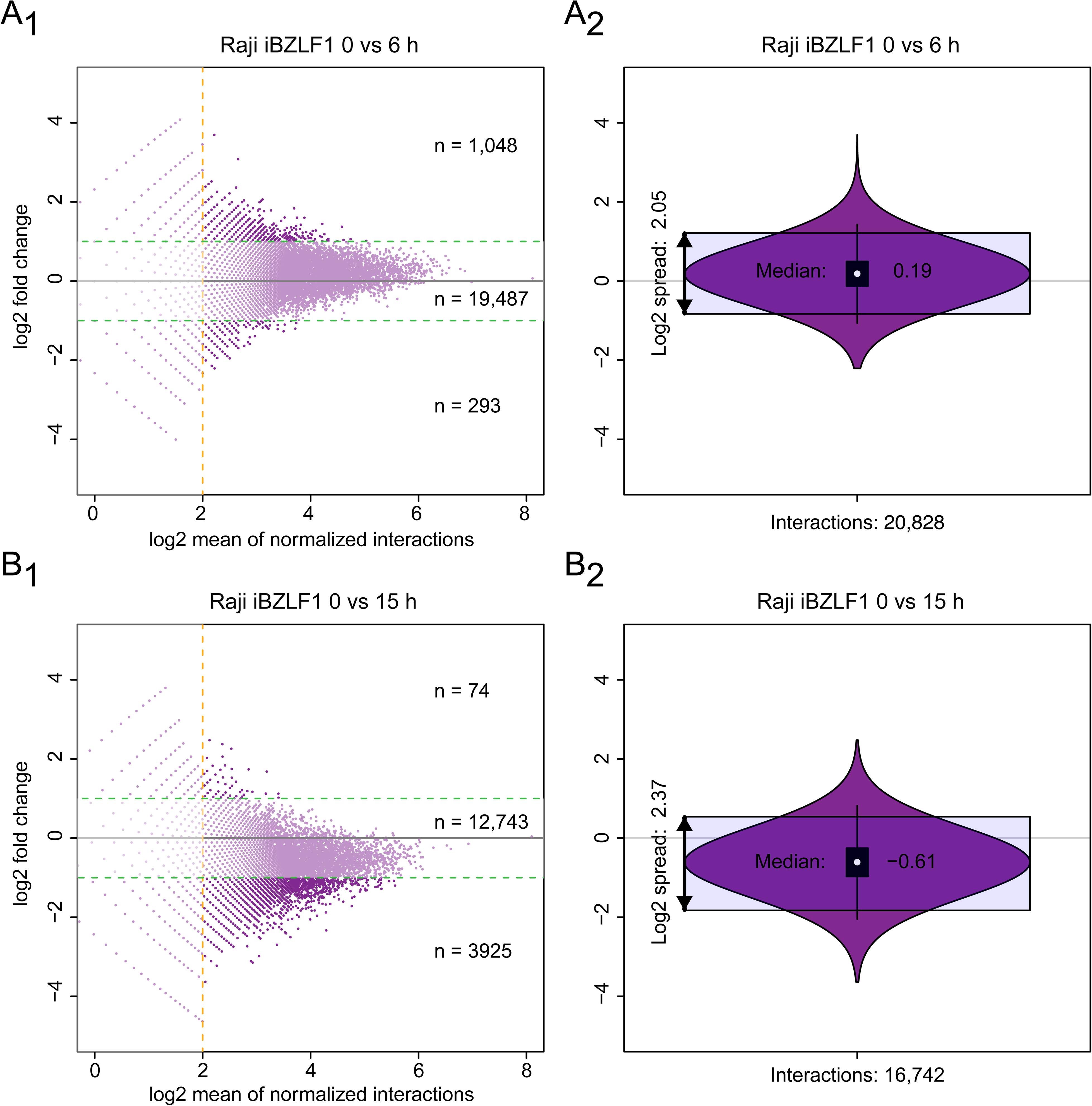
Gain and loss of chromatin interactions in Capture-C experiments 6 and 15 hours after induction of EBV’s lytic cycle. The two MA plots summarize the dynamics of interactions between the promoter regions of 53 analyzed genes and their captured distal DpnII fragments. The x-axis shows the number of identified interactions in log2 scale, the y-axis shows the log2 fold change between the paired time points. **(A1)** After induction of the lytic cycle for six hours a slight global increase of the about 21,000 chromatin interactions is visible. Only fragments with more than 4 interactions were taken into consideration (vertical orange line). About 1,000 interactions showed a more than twofold increase (upper green horizontal line), while about 300 interactions were reduced by at least half (lower green horizontal line). **(A2)** The violin plot summarizes A1 and shows median and population spread. **(B1)** After 15 hours of doxycycline-induced BZLF1 expression almost 4,000 fragments lose more than half of their interactions (lower green horizontal line) compared with the status prior to induction of the lytic phase. Conversely, 74 DpnII fragments show a more than twofold increase of interactions (upper green horizontal line). In total only about 17,000 reach or exceed the threshold of 4 interactions. **(B2)** In comparison with panel A2 the violin plot shows a clear reduction of the median and a slight increase of the population spread.

After bioinformatic analysis and data normalization we visualized the close, medium and distant physical chromatin interactions of these promoter regions in a range of 400,000 nucleotides each on both flanks of the TSS (Fig. 8, Supplementary Figs. S15-S22). The y-axes of the graphs indicate the number of chromatin interactions prior to the induction of BZLF1 (0 h, grey bars) or six hours (orange bars) and 15 hours (blue bars) after the addition of doxycycline. In addition, we plotted BZLF1 binding sites at low and high levels in Raji iBZLF1 cells.

**Figure 8.**
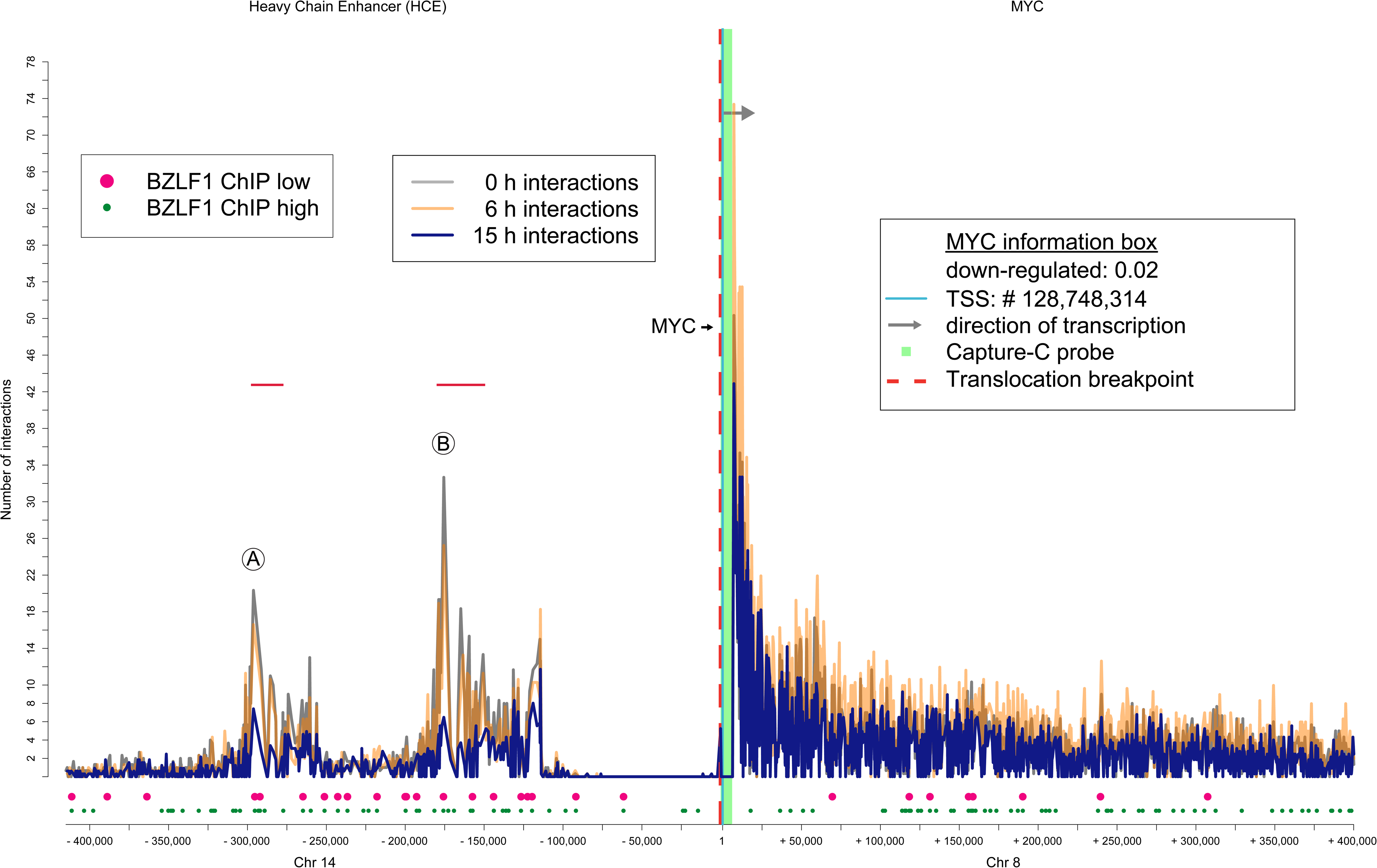
Loss of chromatin interactions between the *MYC* locus and the heavy chain enhancer. On the x-axis, individual chromatin interactions are shown as thin vertical lines of different heights that indicate their interaction frequencies (plotted on the y-axis as “number of interactions”) with the *MYC* promoter region (approximate −5/+5 kb of the TSS). The promoter region is depicted as a green bar in the center of the plot, a grey arrow head points in the direction of transcription, and the light blue line at position “1” indicates the TSS. Grey vertical lines indicate the chromatin interactions prior to BZLF1 induction (0 h), orange and blue lines enumerate interactions 6 and 15 hours after adding doxycycline, respectively. The bottom part of the graph shows the positions of BZLF1 binding sites prior to (big, pink dots) and 15 hours after adding doxycycline (small, green dots). The x-axis indicates the relative nucleotide coordinates (hg19 genome reference) encompassing two flanking regions 400 kb up-and downstream of the TSS. Due to a chromosomal translocation (vertical dashed red line), the heavy chain enhancer (HCE) on chromosome 14 drives the expression of the *MYC* locus on chromosome 8 in Raji cells. Prior to induction of BZLF1, the HCE (A and B) makes frequent contacts (grey vertical lines) with the *MYC* locus. Chromatin interactions decrease after 6 h (orange lines) and are further reduced to about 20 % (blue lines) 15 h after BZLF1 induction. Both HCE interacting A and B loci colocalize with numerous BZLF1 ChIP-seq peaks. In Raji cells, 6 h after BZLF1 induction the level of *MYC* transcripts is reduced to 2 % compared with the non-induced state.

We visualized the actual situation for several genes to inspect the individual gain and loss of chromatin interactions. Supplementary Figure S15 shows the *CD68* gene, which is not regulated (0.99-fold) six hours after induction of the lytic cycle. The number of locus interactions of the promoter region with flanking chromatin modestly increased by about 7 % six hours after induction of the lytic cycle relative to the interactions identified prior to its induction. 15 hours after induction the interactions are reduced to about 63 % relative to the non-induced state. A very similar pattern of altered chromatin interactions was found in the locus of the strongly downregulated *BTG2* gene (regulated by a factor of 0.02; Supplementary Fig. S16) and the upregulated *COL2A1* gene (19-fold upregulated; Supplementary Fig. S17). The number of interactions at the *BTG2* and *COL2A1* loci slightly increased by 12 % and 15 %, respectively, six hours post induction, but decreased by 30 % 15 hours after induction following the general trend shown in Figure 7.

The altered numbers of interactions were particular impressive when the strong, long-range interactions of the *CXCR4* (Supplementary Fig. S18) and *MIR155HG* (Supplementary Fig. S19) genes were considered, which are both down-regulated by factors of 0.04 and 0.33, respectively. The interactions of the *CXCR4* gene remained unchanged on average (101 %) while they increased by about 12 % for the *MIR155HG* gene six hours after induction of BZLF1. After 15 hours, the long-range interactions of the *CXCR4* and *MIR155HG* loci were reduced by 39 % and 34 %, respectively.

All analyzed genes but one followed the observed kinetics and showed a considerable loss of chromatin interactions over time. The exception is *KCNQ5*. Here, the interactions of the flanking region upstream but not downstream of the *KCNQ5* gene remained stable or even slightly intensified 15 hours after BZLF1’s induction (Supplementary Fig. S20).

Additional examples include *E2F2* and *ID3* as well as *TLR6* and *TLR10*, which constitute two separate gene clusters as shown in Supplementary Figures S21 and S22 panels A and B. Again, these genes are in line with the global reduction of chromatin interactions as a late consequence of BZLF1 expression. Interestingly, the two neighboring genes in both examples share interactions but each individual gene is also engaged in gene-specific chromatin-chromatin contacts.

The *MYC* gene, which showed a reduction of its steady state transcript levels by a factor of 50 is an extreme example that demonstrates the reduction of chromatin interactions. In the Burkitt lymphoma cell line Raji the *MYC* gene is not regulated by its genuine *cis*-acting elements, but the gene is under the control of the heavy-chain immunoglobulin enhancer due to a translocation between the chromosomes 8 and 14. The x-axis in Figure 8 is discontinuous and a vertical red dashed line upstream of the *MYC* promoter indicates the chromosomal breakpoint. The two DpnII fragments within the heavy chain enhancer, which make the most frequent contacts with the *MYC* promoter regions are indicated by A and B (Fig. 8). Both interacting sites lose about 20 % of their contacts with the *MYC* promoter locus already six hours after induction of EBV’s lytic cycle. After 15 hours, the number of chromatin contacts at the interacting sites A and B are reduced by 60 % and 80 %, respectively. The two distinct DpnII fragments, A and B, include a 323 nt long repeat (Supplementary Fig. S23, panel A) that comprises 18, partially overlapping BZLF1 sequence motifs (Supplementary Fig. S23, panel B). In our ChIP experiments, both A and B fragments were found to be bound by BZLF1 in Raji iBZLF1 cells (Supplementary Fig. S24, panels A and B).

In summary, all analyzed loci show reduced local and distal chromatin interactions 15 hours after induction of EBV’s lytic cycle. The binding of BZLF1 to certain distant interacting sites is often apparent and occasionally impressive as in the example of the *MYC* gene suggesting that BZLF1 binding to these distal regions directly or indirectly reduces the frequency of chromatin interactions, but this apparent linkage is not supported by our further analysis. DpnII fragments bound by BZLF1 according to our ChIP-seq data and captured in the Capture-C experiment did not differ from captured DpnII fragments to which BZLF1 does not bind (Supplementary Fig. S25 panels A and B). It thus appears as if EBV uses so far unknown mechanisms to reduce cellular chromatin interactions and alter the 3D structure of its host cell chromatin in the lytic phase.

## DISCUSSION

After infection and a short period of pre-latency, EBV establishes its latent phase, which results in a very stable host-virus relationship. ln this phase, only very few proteins are expressed *in vivo* probably to avoid the recognition and elimination of virally infected cells by cellular immunity. Upon escape from latency the immediate early viral protein BZLF1 is the first protein of the lytic phase that is expressed. It induces the synthesis of additional viral proteins important for the complete transcriptional activation of the lytic phase, viral factors for the autonomous viral DNA replication and viral genes that fend off the antiviral immunity of the host organism. Therefore, the host cell has to be manipulated and reorganized to free macromolecules, energy, cellular machines, and transport capacity to support directly viral transcription, DNA replication, and viral morphogenesis, processes that also require space and occupy nuclear compartments to allow EBV’s reproduction. Since BZLF1 shows a strong structural homology with members of the cellular AP-1 protein family (Petosa et al., 2006) and binds their sequence motifs (Urier et al., 1989) genome wide, we not only examined global changes in the cellular genome and transcriptome, but also asked if BZLF1 is directly involved in these changes.

### BZLF1 reaches a high protein level in the lytic phase of B95-8 cells

In a first step we investigated the BZFL1 protein levels in the fraction of B95-8 cells that are in the lytic phase of infection. We found very high molecule numbers per cell (Supplementary Fig. S1B) that are also achieved upon doxycycline-mediated induction in our Raji cell model. In this model, the conditional BZLF1 allele has a basal leaky expression, but at this level BZLF1 was incapable of inducing EBV’s lytic cycle in Raji iBZLF1 cells (Woellmer et al., 2012) and in HEK 293 2089 iBZLF1 cells (Fig. 1, Supplementary Fig. S1C). With this knowledge we assumed that our model truly reflects the functions of BZLF1 in latently infected and lytically induced B cells.

### BZLF1 binds two major motifs in the cellular genome with high frequency and specificity

At high level BZLF1 binds to several hundred thousand sites in the Raji cell genome, which is in agreement with the diffuse nuclear distribution of BZLF1 (Park et al., 2008). We identified two major sequence motifs, one of which is found in 96 % of all called BZLF1 peaks. The high number of differently composed DNA sequence motifs led us to rank them according to their presumed relative affinities of BZLF1 binding (Fig. 3). Their composition partially confirms previous findings of Ramasubramanyan et. al. (Ramasubramanyan et al., 2015), who, very much in contrast to our study, identified 5,000 BZLF1 binding sites in cellular genomic DNA, only. Our study now extends this set of BZLF1 binding motifs and reveals that BZLF1 also binds closely related motifs that terminate with a thymidine residue (Fig. 3).

The many cellular BZLF1 binding motifs might have an obvious function supporting EBV’s life style. A cell that incompletely activates EBV’s lytic phase is likely prone to be eliminated by immediate antiviral T cell responses, because the many lytic immunoevasive viral genes are not timely and properly expressed to fend off cytotoxic CD4^+^ and CD8^+^ effector T cells (Ressing et al., 2015). It is therefore plausible that the transition from the latent to the lytic phase operates as a dichotomous switch and not as a rheostat. During a perhaps erratic expression of low BZLF1 levels the cellular genome with its abundant number of high-affinity CpG-ZRE binding motifs might act as a sink abrogating the activation of a subset of viral lytic promoters by chance. Only a strong inducing signal leads to BZLF1 levels sufficient to bind both cellular and viral CpG containing high-affinity motifs (TGWGCGA, Fig. 2C and TGWGCGT, Supplementary Fig. S4C) and, subsequently, the two canonical non-CpG motifs (TGWGYVA, Fig. 2C and TGWGYVT, Supplementary Fig. S4C). We know that BZLF1 must bind certain low affinity binding sites in the viral lytic origin of DNA replication to support efficient viral DNA amplification (Bergbauer et al., 2010; Schepers et al., 1996). Thus, the diverse and abundant BZLF1 binding motifs in cellular DNA (>5×10^5^) together with the 85 identified viral binding motifs likely cooperate to implement a molar switch that only flips when a certain threshold concentration of BZLF1 protein is reached.

### The presumed functions of cellular BZLF1 binding sites

BZLF1’s role in cellular chromatin is not obvious, but it might act at different levels. (i) BZLF1 could induce the expression of a few cellular factors that serve important functions in EBV’s lytic phase similar to its role on viral genomic DNA. For example, BZFL1 has to induce the BMLF1-BSLF2 gene (Kenney et al., 1989) such that they together can activate certain early EBV promoters (Chevallier-Greco et al., 1986). BZLF1 acts as a replication factor together with the viral factors BALF2, BALF5, BBLF2/3, BBLF4, BMRF1, and BSLF1 to initiate lytic viral DNA replication (Hammerschmidt and Sugden, 2013), but BZLF1 also induces the expression of these genes mainly via meZRE motifs (Fig. 2C) with methylated CpG dinucleotides (Bergbauer et al., 2010) and probably together with BRLF1. Similarly, BZLF1 has been reported to induce (or repress) numerous cellular genes (Flemington and Speck, 1990; Cayrol and Flemington, 1995; Lu et al., 2000; Morrison et al., 2001; Morrison et al., 2004; Chang et al., 2006; Jones et al., 2007; Hsu et al., 2008; Tsai et al., 2009; Heather et al., 2009; Bristol et al., 2010; Zuo et al., 2011), but a clear picture depicting how BZLF1 regulates cellular targets is lacking. (ii) BZLF1 is known to induce a DNA damage response (DDR), which seems to be needed for the full expression of BZLF1 and other downstream viral factors such as BMRF1. The induction of a DDR requires that BZLF1 binds DNA (Wang’ondu et al., 2015). Thus, cellular BZLF1 binding sites might contribute to DDR induction, but how a DDR could functionally support EBV’s lytic phase is uncertain. Lastly, (iii) the many cellular BZLF1 binding sites might be non-functional similar to 46-99 % of all binding sites of 59 different cellular transcription factors (Cusanovich et al., 2014). The binding of transcription factors to these sites did not lead to a measurable change in transcription levels of putative target genes (Cusanovich et al., 2014), but in the case of BZLF1 the many binding sites in cellular chromatin might act as a sink to prevent the erratic induction of EBV’s lytic.

### BZLF1 induces open chromatin at its cellular binding sites but global chromatin accessibility is reduced in EBV’s pre-replicative, lytic phase.

Regions of open chromatin allow the binding of cellular factors, protein complexes and large nuclear machines. Open chromatin is a prerequisite for chromatin-chromatin interactions that contribute to the formation of topologically associating domains (TADs), chromosome territories and, ultimately, the nuclear architecture. Our genome-wide ATAC-seq results indicate that the global accessibility of cellular chromatin is considerably reduced upon induction of EBV’s lytic cycle (Fig. 4B). Loss of open chromatin is generally linked with transcriptional repression, which is in line with our transcriptomic data (Fig. 5). To our knowledge, only during embryonic development the loss of chromatin-chromatin interactions is a noted feature, which is seen in conjunction with the formation of new TADs (Kaaij et al., 2018). Alternatively, an early and general loss or even collapse of discrete chromatin-chromatin interactions might prepare for a subsequent spatial regulation of cellular chromatin including the increase in volume of the nucleus of herpesvirus infected cells, when replication compartments or amplification factories form (Monier et al., 2000; Chiu et al., 2013).

Very much in contrast to the global loss of open cellular chromatin, the binding of BZLF1 to silent chromatin increases its local accessibility. BZLF1 is a pioneer factor (Schaeffner et al., 2019) and shares this function with other pioneer factors that bind to repressed and generally inaccessible sequence motifs and recruit chromatin remodelers to mobilize or even evict nucleosomes at and in close proximity of their cognate binding sites. Why BZLF1 opens chromatin at its many cellular binding sites is currently unclear.

### EBV’s lytic cycle induces a massive reduction of cellular transcripts

Upon induction of BZLF1 in Raji cells, only few cellular genes are upregulated, while the majority of transcripts (>7,000) are strongly reduced (Fig. 5C). Conversely, viral transcripts are massively upregulated to accumulate to >140,000 reads per viral gene (Fig. 1).

The reduction of global transcripts depends on BZLF1’s transactivation domain. It could cause a restructuring of the cellular chromatin architecture as seen in Figures 4B, 7, and 8 with a concomitant decrease of cellular transcription or it might induce additional viral genes that indirectly contribute to this reduction. A likely viral candidate is the BGLF5 gene that is highly expressed (87,000 reads, 170-fold induced) already 6 hours after the activation of EBV’s lytic phase. BGLF5 is an RNA exonuclease (Buisson et al., 2009) that destroys viral as well as cellular mRNAs nonspecifically (Horst et al., 2012) causing the so-called host-shutoff. Additionally, BGLF5 was described to potentially contribute to immune evasion (Rowe et al., 2007; Zuo et al., 2008) and to support the nuclear translocation of the PABPC protein together with BZLF1. A nuclear accumulation of PABPC leads to hyperadenylated transcripts blocking their nuclear export and translation (Kumar and Glaunsinger, 2010; Park et al., 2014).

Ramasubramanyan et al. also analyzed the transcriptome of Akata cells, an EBV-positive Burkitt lymphoma cell line (Ramasubramanyan et al., 2015), but did not see a global reduction of cellular transcripts in the lytic phase. The authors identified 2,300 genes of which ¾ were up-and ¼ down-regulated 24 hours after lytic induction of Akata cells. In contrast, we found 99 % of cellular genes down-and 1 % up-regulated in Raji iBZLF1 cells 6 hours post induction (Fig. 5C). The discrepancy between both studies can be explained by the chosen threshold levels of bioinformatic analysis and mainly by the method of normalization. Based on the published literature (Buisson et al., 2009; Horst et al., 2012; Kumar and Glaunsinger, 2010; Park et al., 2014) we expected the mRNA content between non-induced and induced cells to differ substantially. Therefore, we added known molar amounts of polyadenylated ERCC spike-in RNAs to total RNA samples from identical numbers of viable cells. Conventional data processing and normalization approaches mostly rely on average principles and center the mean of the fold change around zero independently of the RNA amount. We know from a conventional analysis of our data that we would have missed the profound global transcriptomic modifications in this model of viral activation had we chosen this bioinformatic approach.

### Cellular gene regulation and promoter-binding of BZLF1 are not necessarily linked

BZLF1 is known to be a promoter factor in the viral genome and was also described to up-or downregulate cellular genes in different cells. (Flemington and Speck, 1990; Cayrol and Flemington, 1995; Lu et al., 2000; Morrison et al., 2001; Morrison et al., 2004; Chang et al., 2006; Jones et al., 2007; Hsu et al., 2008; Tsai et al., 2009; Heather et al., 2009; Bristol et al., 2010; Zuo et al., 2011). A recent report even suggested that BZLF1 might directly or indirectly regulate about 10% of all protein-encoding genes in B cells (Ramasubramanyan et al., 2015). When we correlated cellular gene regulation and the occurrence of BZLF1 sites in promoter regions, many genes with BZLF1 binding sites were not regulated. This finding is in accordance with previous reports in which transcription factors were found to regulate only a subset of genes with which they are spatially associated (Cusanovich et al., 2014; Gao et al., 2004; Hu et al., 2007). In Raji cells, BZLF1 binding sites could be found in 37-55 % of apparently regulated genes, only (Fig. 6). This correlative finding contradicts the idea that BZLF1 binds to cellular promoters and induces cellular gene expression. No BZLF1 binding sites were identified in promoters of 45-63 % of genes that are regulated after induced expression of BZLF1. Clearly, at the level of the lytically induced cells, BZLF1 does not follow the rules found in the viral genome, where BZLF1 binds in close proximity to the transcriptional start sites of its regulated genes (Bergbauer et al., 2010).

### Upon induction of EBV’s lytic cycle, cellular chromatin connections detach

Using the Capture-C technique we studied the chromatin structure of 53 selected cellular genes and asked if BZLF1’s local binding contributed to their regulation. In latently infected cells we identified many distal regions that contact the promoters’ captured regions −5kb/+5kb of the transcriptional start sites. Uniformly, the number of chromatin interactions rises slightly 6 hours post induction but is substantially reduced at 15 hours irrespective of whether the analyzed genes were up-, down-, or non-regulated.

It is very unlikely that BZLF1 is directly involved in the wide-spread loss of chromatin interactions, because BZLF1 binding and the loss of chromatin interactions do not seem to correlate (Supplementary Fig. S25). The regulation of the *MYC* locus could be an exception, because BZLF1 might be directly involved in the reduction of chromatin-chromatin contacts between the heavy chain enhancer *HCE* and the *MYC* locus (Fig. 8). No BZLF1 binding sites are located in the promoter region of the *MYC* gene, but it forms strong contacts with *HCE*, which is rich in BZLF1 binding sites. Already six hours after induction, *HCE-MYC* chromatin interactions decrease considerably and the *MYC* mRNA level is reduced to two percent, only (Fig. 8).

A BZLF1 regulated lytic viral gene such as BGLF5 is a possible candidate for this task, since it is not only an RNAse but also a DNAse that might affect genomic stability (Baylis et al., 1989). BGLF5 acts as an endonuclease (Zhang et al., 1987; Stolzenberg and Ooka, 1990; Lin et al., 1995) and exonuclease (Lin et al., 1995) and was found to induce DNA damage in the cellular genome and instability of microsatellites in epithelial cells (Wu et al., 2010). A second viral candidate that might contribute to the loss of chromatin interactions is BALF3, which induces DNA strand breaks, meditates genome instability (Chiu et al., 2014a), and supports efficient virion synthesis (Chiu et al., 2014b).

In conclusion, upon induction of EBV’s lytic phase we found that cellular chromatin-chromatin interactions are lost and the transcriptome of the cell is profoundly reduced. Regions of accessible, hence open chromatin lose their accessibility, but, conversely, BZLF1 binding sites in repressed cellular chromatin gain accessibility. These alterations in cellular chromatin are massive and global suggesting that they contribute to the success of lytic induction and viral *de novo* synthesis. We speculate that these alterations help to provide space for lytic viral DNA replication (Monier et al., 2000; Chiu et al., 2013) and/or for capsid morphogenesis and mobility (Bosse et al., 2015) and redirect basic cellular functions including transcription, mRNA transport and export such that these resources and functions become available for the synthesis of viral transcripts and viral DNA and additional virus-related downstream processes.

## ACCESSION NUMBERS AND DATA AVAILABILITY

NGS files have been deposited on ArrayExpress (Kolesnikov et al., 2015) using the web-based submission tool Annotare 2.0 (https://github.com/arrayexpress/annotare2). The data files of interest can be downloaded via https://www.ebi.ac.uk/arrayexpress/ using the following accession numbers:

anti-BZLF1ChIP-seq Raji iBZLF1: http://www.ebi.ac.uk/arrayexpress/experiments/E-MTAB-7788 anti-BZLF1 ChIP-seq parental DG75: to be added.

ATAC-seq: http://www.ebi.ac.uk/arrayexpress/experiments/E-MTAB-7789

RNA-seq: to be added.

Capture-C: to be added.

## SUPPLEMENTARY DATA

Supplementary Data are available in separate files and include 25 Supplementary Figures, their figure legends, and three Supplementary Tables.

## ACKNOWLEDGEMENT

We thank Alexander Graf (Laboratory for Functional Genome Analysis, Gene Center, Ludwig Maximilians University Munich, Munich, Germany) for the introduction to the Galaxy suit and the initial analysis of RNA-seq data, Sylvia Mallok (Laboratory for Functional Genome Analysis, Gene Center, Ludwig Maximilians University Munich, Munich, Germany) for the help with the preparation of the RNA-seq and Capture-C libraries, Tamas Schauer (Molecular Biology Division, Biomedical Center, Faculty of Medicine, Ludwig-Maximilians University Munich, Planegg-Martinsried, Germany) for the code for ERCC spike-in normalization, Thomas Schwarzmayr (Institute of Human Genetics, Helmholtz Zentrum München, Germany) for the code for the visualization of ERCC spike-in NGS sequencing reads, and James Davies (Trinity College, Bristol, UK) for supporting our understanding of the Capture-C analysis.

## AUTHORS CONTRIBUTIONS

A.B., P.M.-G., S.K., H.B., F.M.C., D.P., T.S., and W.H. designed and performed experiments and analyzed data. A.B., G.S., T.S., and W.H. designed and conceived the project. A.B. and W.H. wrote the manuscript.

