## Supplementary Figures S1-S25 for "Epstein-Barr virus inactivates the transcriptome and disrupts the chromatin architecture of its host cell in the first phase of lytic reactivation"

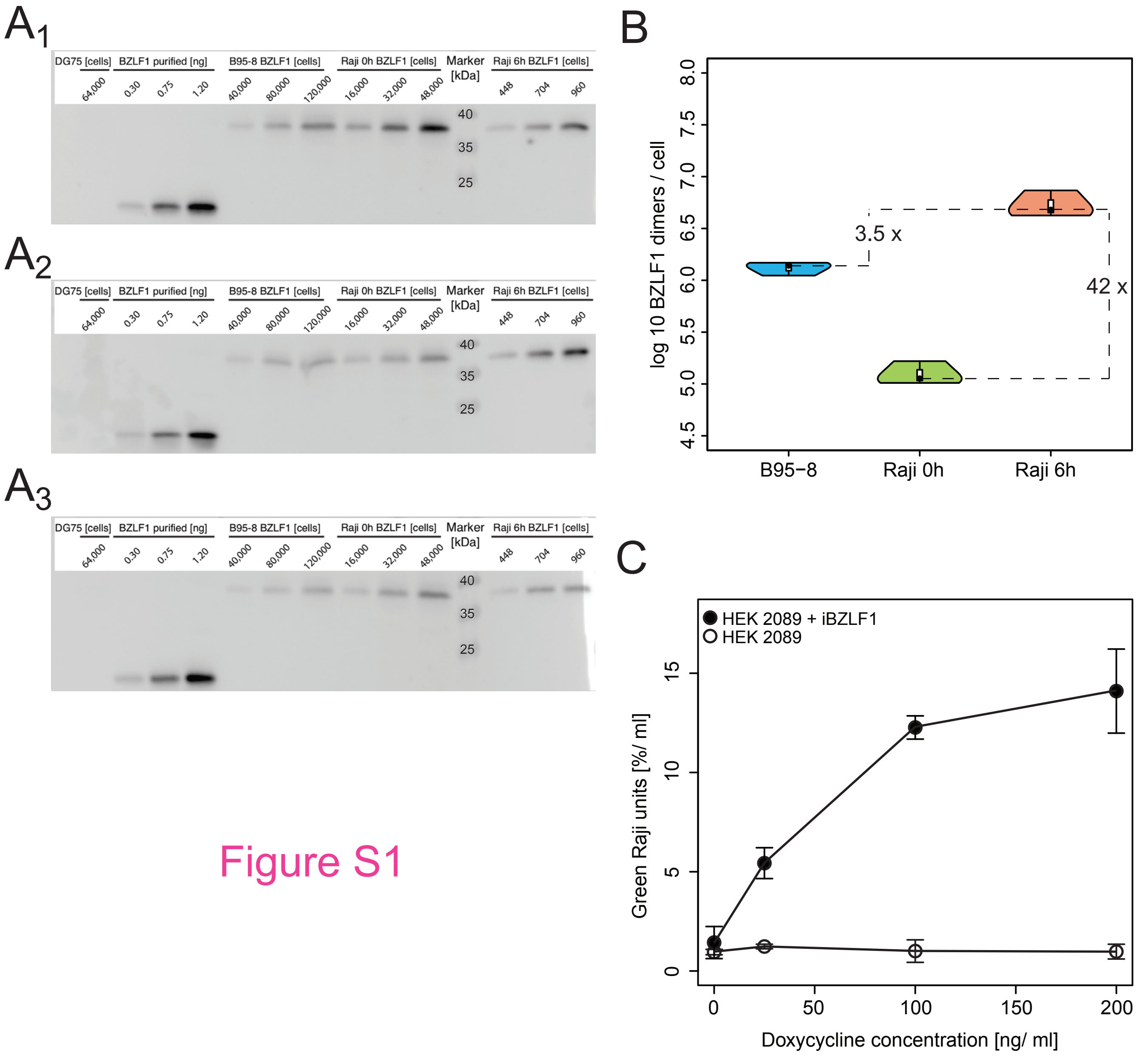

A1
A2
A3
DG75 [cells]
BZLF1 purified [ng]
B95-8 BZLF1 [cells]
Raji 0h BZLF1 [cells]
Marker [kDa]
Raji 6h BZLF1 [cells]
64,000
0.30
0.75
1.20
40,000
80,000
120,000
16,000
32,000
48,000
448
704
960
40
35
25
B
log 10 BZLF1 dimers / cell
8.0
7.5
7.0
6.5
6.0
5.5
5.0
4.5
3.5 x
42 x
B95−8
Raji 0h
Raji 6h
C
HEK 2089 + iBZLF1
HEK 2089
Green Raji units [%/ ml]
15
10
5
0
0
50
100
150
200
Doxycycline concentration [ng/ ml]


Figure S1

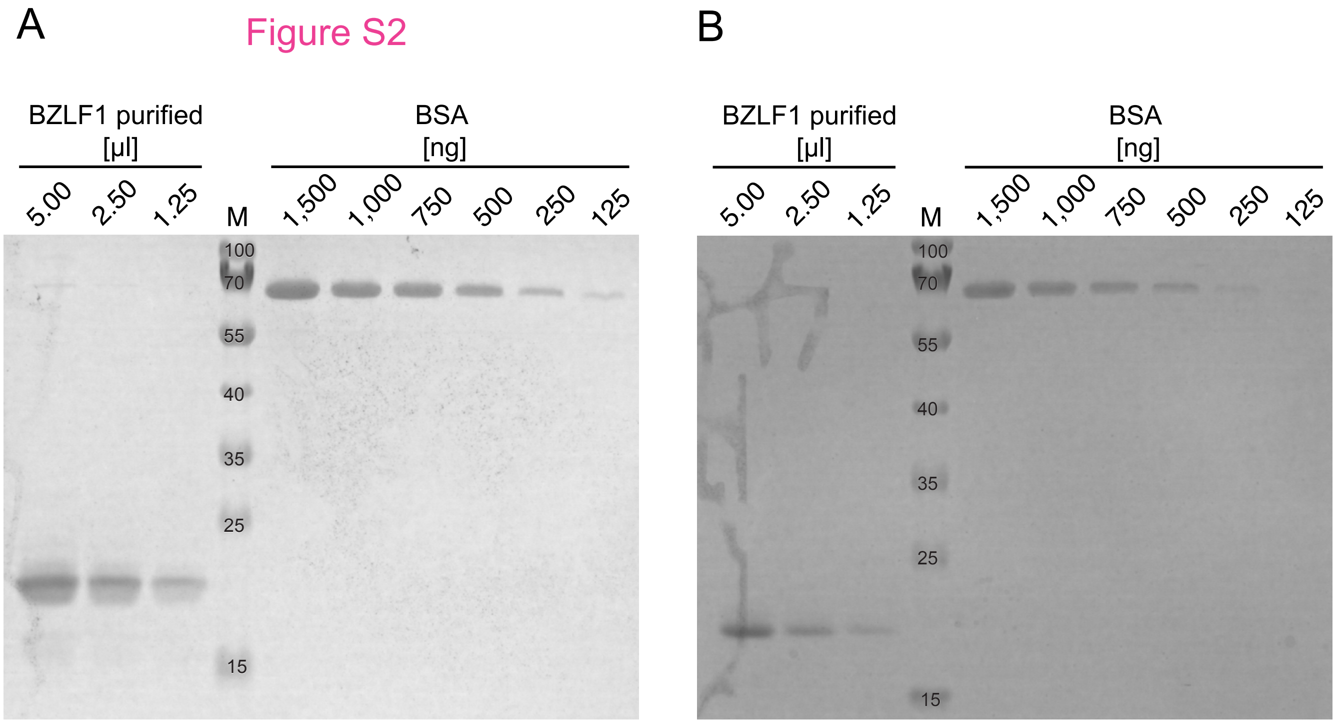
A
Figure S2
B
BZLF1 purified [µl]
5.00
2.50
1.25
M
BSA [ng]
1,500
1,000
750
500
250
125
100
70
55
40
35
25
15
BZLF1 purified [µl]
5.00
2.50
1.25
M
BSA [ng]
1,500
1,000
750
500
250
125
100
70
55
40
35
25
15

### Figure S3

#### Raji iBZLF1

**A**

##### low level

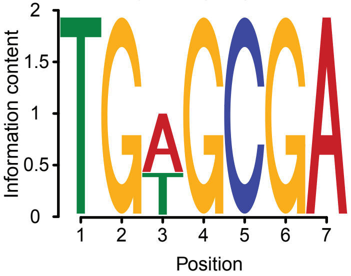


within 30346 sequences

| Motif | RC Motif | Positive | Negative | P-value | E-value |
|---|---|---|---|---|---|
| TCGCWCA | <u>TGWGCGA</u> | 21387 | 1543 | 7.3e-6832 | 3.4e-6827 |
| TCGCTCA | <u>TGAGCGA</u> | 12716 | 775 | 1.6e-3438 | 7.3e-3434 |
| TCGCACA | <u>TGTGCGA</u> | 9211 | 803 | 5.9e-2107 | 2.7e-2102 |

**B**

##### high level

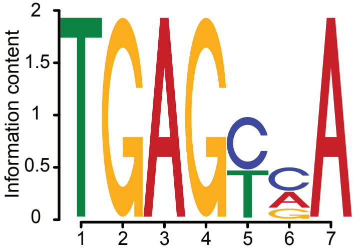


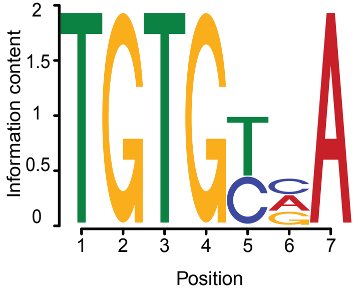


within 145544 sequences

| Motif | RC Motif | Positive | Negative | P-value | E-value |
|---|---|---|---|---|---|
| TBRCTCA | <u>TGAGYVA</u> | 110167 | 50296 | 1.8e-11160 | 8.3e-11156 |
| TGACTCA | <u>TGAGTCA</u> | 38251 | 11185 | 2.9e-4064 | 1.4e-4059 |
| TGGCTCA | <u>TGAGCCA</u> | 40353 | 13528 | 1.2e-3696 | 5.8e-3692 |
| TTGCTCA | <u>TGAGCAA</u> | 33772 | 13527 | 3.6e-2312 | 1.7e-2307 |
| TCGCTCA | <u>TGAGCGA</u> | 16240 | 3333 | 5.3e-2149 | 2.5e-2144 |
| TTACTCA | <u>TGAGTAA</u> | 22306 | 6824 | 9.0e-2084 | 4.2e-2079 |
| TCACTCA | <u>TGAGTGA</u> | 17785 | 12185 | 3.3e-257 | 1.5e-252 |

| Motif | RC Motif | Positive | Negative | P-value | E-value |
|---|---|---|---|---|---|
| TBRCACA | <u>TGTGYVA</u> | 80012 | 50325 | 1.2e-2681 | 5.5e-2677 |
| TTGCACA | <u>TGTGCAA</u> | 24280 | 12999 | 1.0e-864 | 4.7e-860 |
| TCGCACA | <u>TGTGCGA</u> | 10454 | 3468 | 8.5e-838 | 3.9e-833 |
| TTACACA | <u>TGTGTAA</u> | 16914 | 7956 | 7.8e-785 | 3.6e-780 |
| TGACACA | <u>TGTGTCA</u> | 22354 | 12890 | 2.0e-637 | 8.9e-633 |
| TGGCACA | <u>TGTGCCA</u> | 20046 | 14474 | 5.5e-225 | 2.5e-220 |
| TCACACA | <u>TGTGTGA</u> | 16530 | 13985 | 8.1e-054 | 3.7e-049 |

### Figure S4

A

Raji iBZLF1 cells

| BZLF1 level | low | high |
|---|---|---|
| ChIP peaks | 30,346 | 145,544 |
| peaks with ≥ 1 major BZLF1 motifs | 1,930 (6 %) | 84,546 (58 %) |

Peaks

B

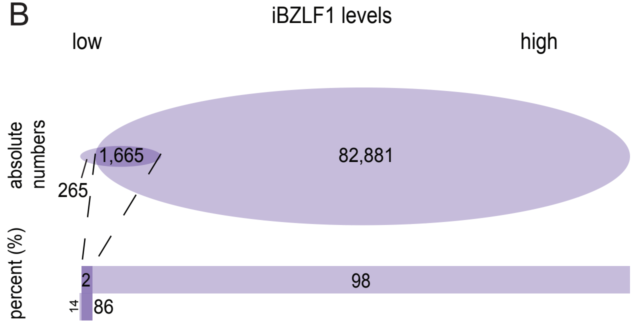


C

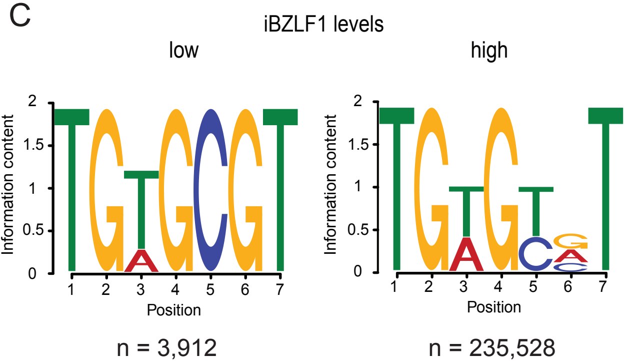


Motifs

D

iBZLF1 levels

| low | high |
|---|---|
| 2.0 motifs/peak (3,912 motifs/1,930 peaks) | 2.8 motifs/peak (235,528 motifs/84,546 peaks) |

Ratio

### Figure S5

Raji iBZLF1

**A**

low level

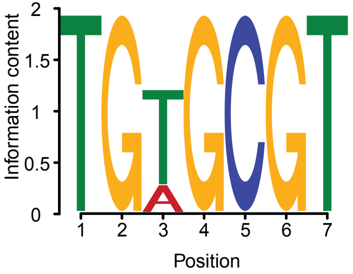


within 1930 sequences

| Motif | RC Motif | Positive | Negative | P-value | E-value |
|---|---|---|---|---|---|
| ACGCWCA | TGWGCGT | 1930 | 505 | 1.1e-622 | 5.1e-618 |
| ACGCACA | TGTGCGT | 1371 | 445 | 3.7e-204 | 1.7e-199 |
| ACGCTCA | TGAGCGT | 693 | 95 | 1.1e-138 | 4.9e-134 |

**B**

high level

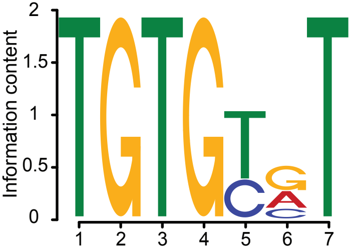


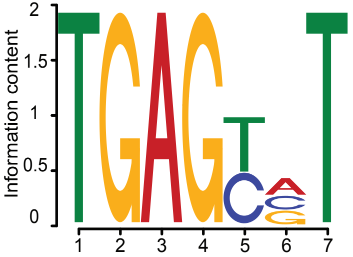


within 84546 sequences

| Motif | RC Motif | Positive | Negative | P-value | E-value |
|---|---|---|---|---|---|
| ABRCACA | TGTGYVT | 59709 | 31367 | 4.3e-4238 | 2.0e-4233 |
| ACACACA | TGTGTGT | 24979 | 10810 | 1.4e-1582 | 6.6e-1578 |
| ACGCACA | TGTGCGT | 10412 | 3239 | 1.4e-935 | 6.6e-931 |
| ATACACA | TGTGTAT | 14621 | 6374 | 3.1e-825 | 1.5e-820 |
| ATGCACA | TGTGCAT | 18375 | 9517 | 4.9e-745 | 2.3e-740 |
| AGACACA | TGTGTCT | 14427 | 8420 | 3.1e-403 | 1.4e-398 |
| AGGCACA | TGTGCCT | 13975 | 8758 | 2.8e-305 | 1.3e-300 |

| Motif | RC Motif | Positive | Negative | P-value | E-value |
|---|---|---|---|---|---|
| ABRCTCA | TGAGYVT | 39890 | 26970 | 7.1e-904 | 3.2e-899 |
| ATACTCA | TGAGTAT | 6912 | 3458 | 1.7e-273 | 7.8e-269 |
| ATGCTCA | TGAGCAT | 11292 | 7140 | 3.2e-232 | 1.4e-227 |
| ACGCTCA | TGAGCGT | 4025 | 1769 | 4.5e-205 | 2.0e-200 |
| AGGCTCA | TGAGCCT | 8209 | 6137 | 1.5e-073 | 6.7e-069 |
| ACACTCA | TGAGTGT | 9097 | 7357 | 1.4e-046 | 6.4e-042 |
| AGACTCA | TGAGTCT | 7507 | 6213 | 5.0e-031 | 2.2e-026 |

### Figure S6

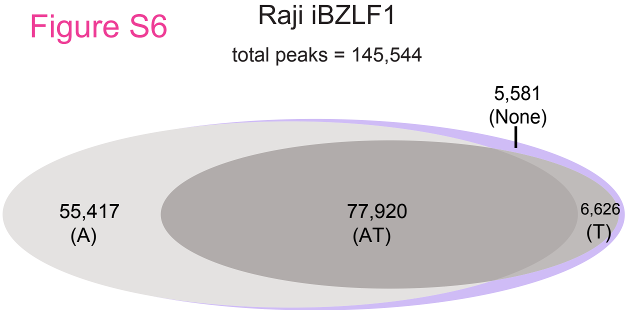
Raji iBZLF1
total peaks = 145,544
5,581
(None)
55,417
(A)
77,920
(AT)
6,626
(T)

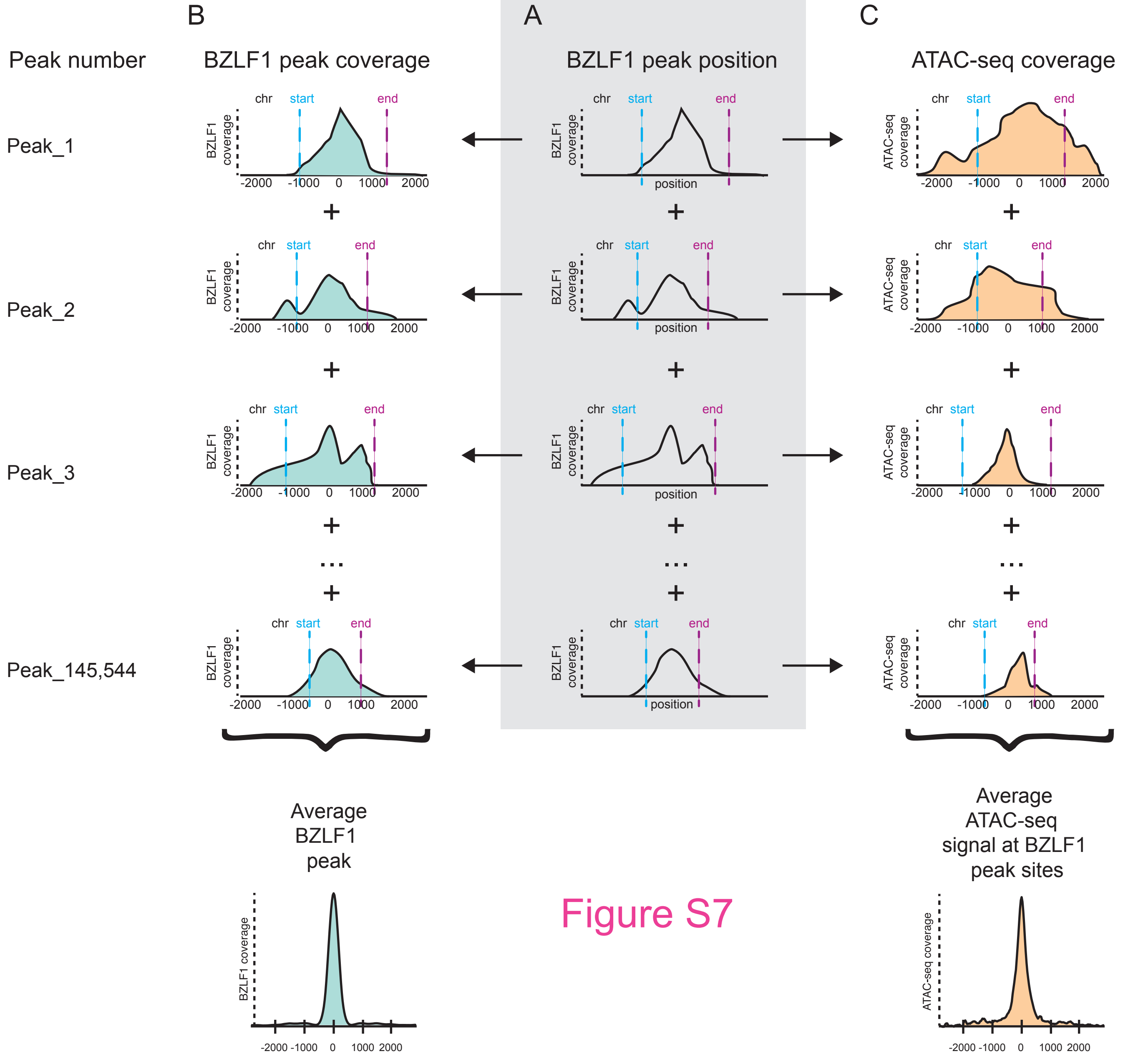

B
A
C
Peak number
BZLF1 peak coverage
BZLF1 peak position
ATAC-seq coverage
Peak_1
Peak_2
Peak_3
Peak_145,544
chr
start
end
BZLF1 coverage
ATAC-seq coverage
position
-2000
-1000
0
1000
2000
+
...
Average BZLF1 peak
Average ATAC-seq signal at BZLF1 peak sites


Figure S7

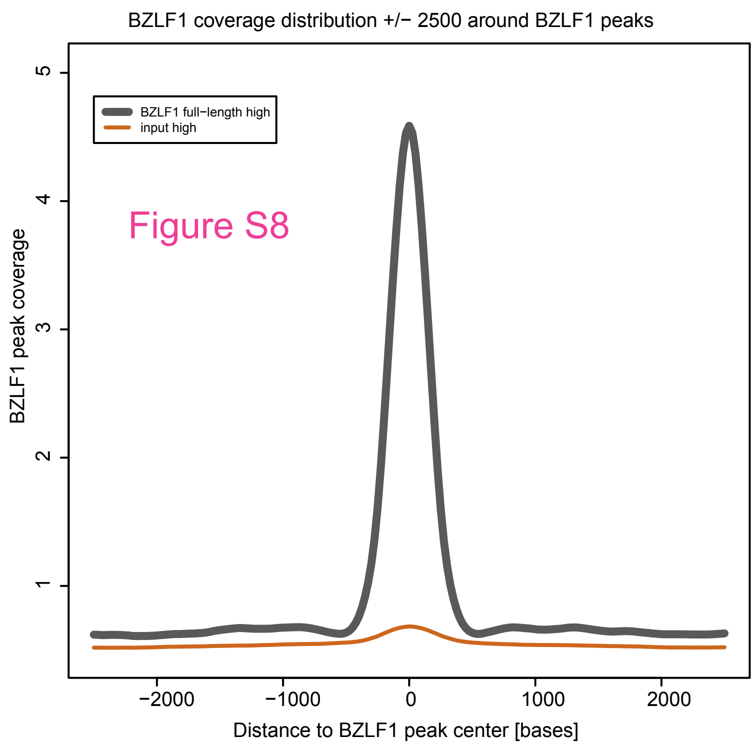
BZLF1 coverage distribution +/− 2500 around BZLF1 peaks
BZLF1 full−length high
input high
Figure S8
BZLF1 peak coverage
1
2
3
4
5
−2000
−1000
0
1000
2000
Distance to BZLF1 peak center [bases]

### Figure S9

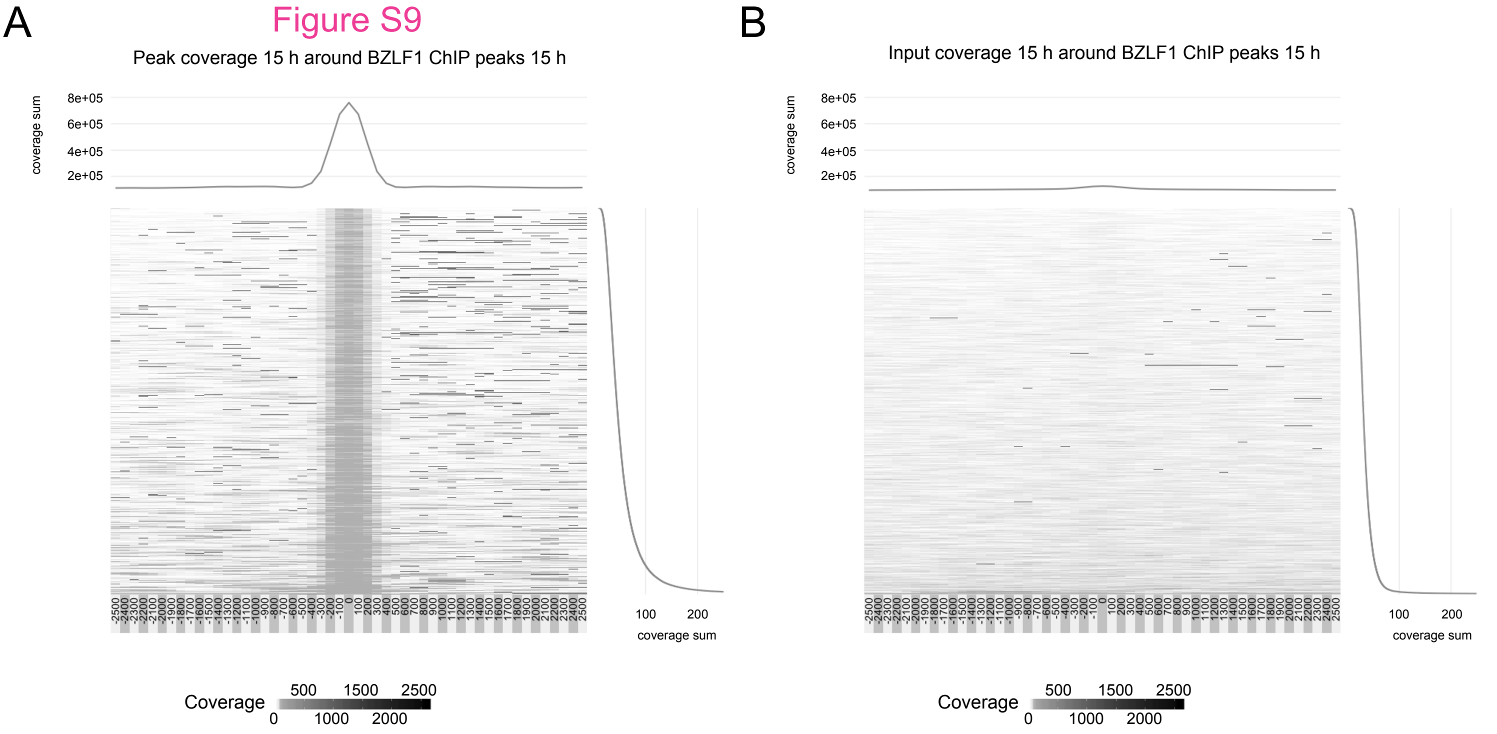
A
Peak coverage 15 h around BZLF1 ChIP peaks 15 h
coverage sum
8e+05
6e+05
4e+05
2e+05
100
200
coverage sum
Coverage
500 1500 2500
0 1000 2000
B
Input coverage 15 h around BZLF1 ChIP peaks 15 h
coverage sum
8e+05
6e+05
4e+05
2e+05
100
200
coverage sum
Coverage
500 1500 2500
0 1000 2000

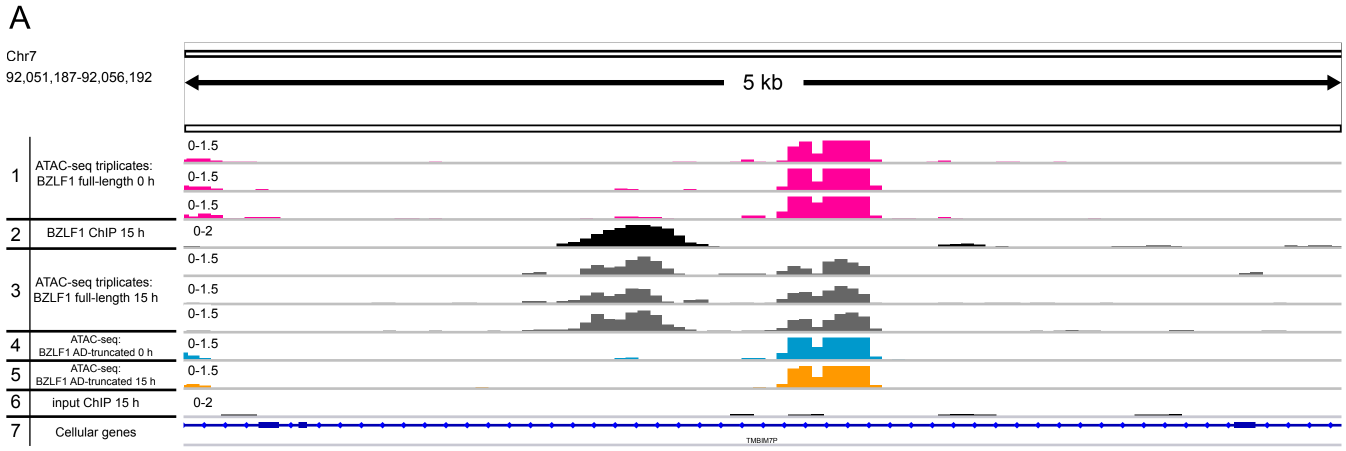
A
Chr7
92,051,187-92,056,192
5 kb
1 ATAC-seq triplicates: BZLF1 full-length 0 h
0-1.5
0-1.5
0-1.5
2 BZLF1 ChIP 15 h
0-2
3 ATAC-seq triplicates: BZLF1 full-length 15 h
0-1.5
0-1.5
0-1.5
4 ATAC-seq: BZLF1 AD-truncated 0 h
0-1.5
5 ATAC-seq: BZLF1 AD-truncated 15 h
0-1.5
6 input ChIP 15 h
0-2
7 Cellular genes
TMBIM7P

### Figure S10

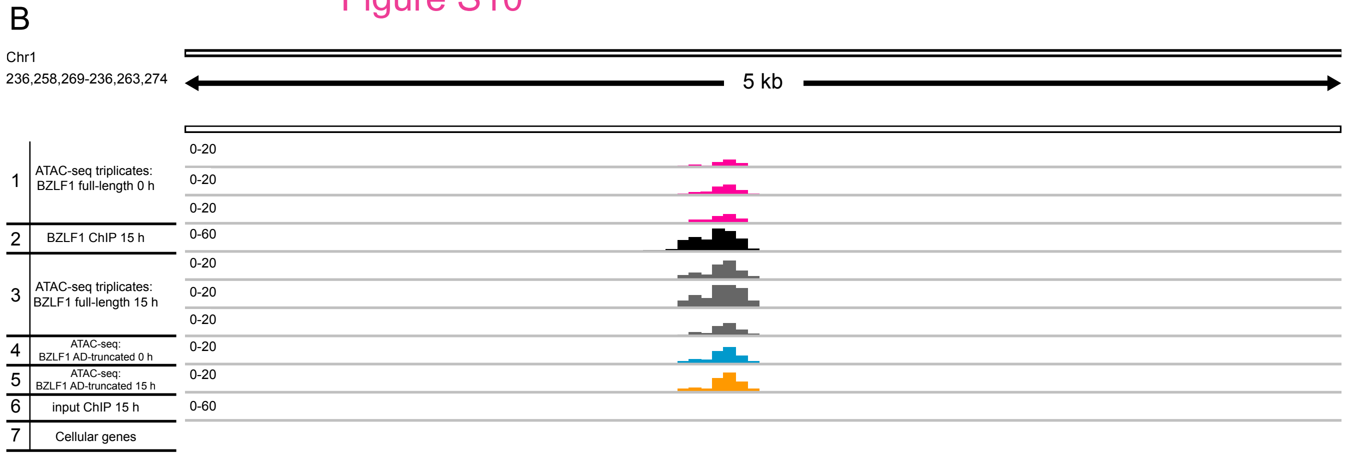
B
Chr1
236,258,269-236,263,274
5 kb
1 ATAC-seq triplicates: BZLF1 full-length 0 h
0-20
0-20
0-20
2 BZLF1 ChIP 15 h
0-60
3 ATAC-seq triplicates: BZLF1 full-length 15 h
0-20
0-20
0-20
4 ATAC-seq: BZLF1 AD-truncated 0 h
0-20
5 ATAC-seq: BZLF1 AD-truncated 15 h
0-20
6 input ChIP 15 h
0-60
7 Cellular genes

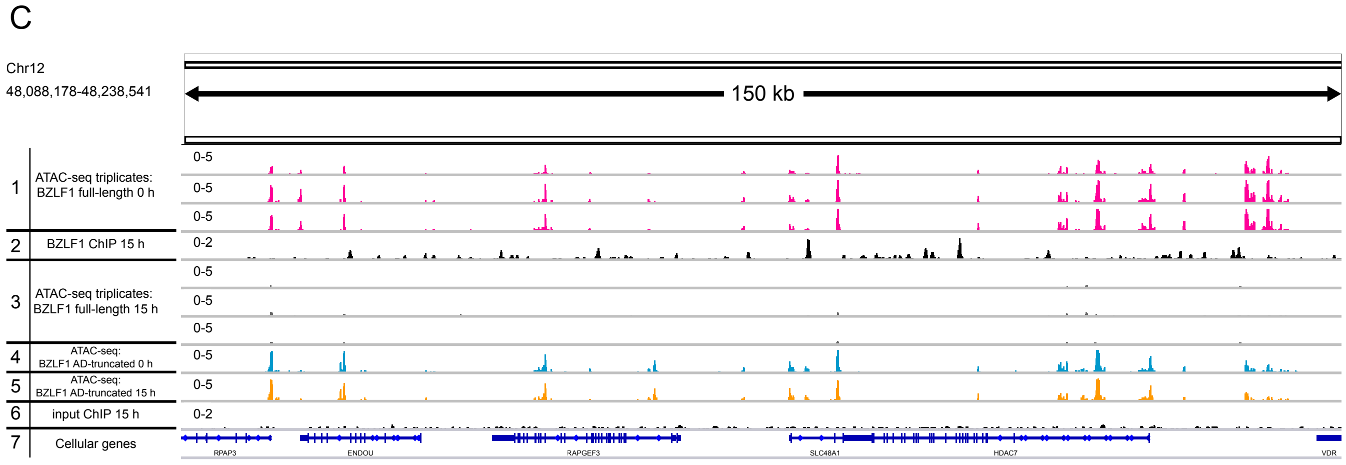
C
Chr12
48,088,178-48,238,541
150 kb
1 ATAC-seq triplicates: BZLF1 full-length 0 h
0-5
0-5
0-5
2 BZLF1 ChIP 15 h
0-2
3 ATAC-seq triplicates: BZLF1 full-length 15 h
0-5
0-5
0-5
4 ATAC-seq: BZLF1 AD-truncated 0 h
0-5
5 ATAC-seq: BZLF1 AD-truncated 15 h
0-5
6 input ChIP 15 h
0-2
7 Cellular genes
RPAP3
ENDOU
RAPGEF3
SLC48A1
HDAC7
VDR

Figure S11

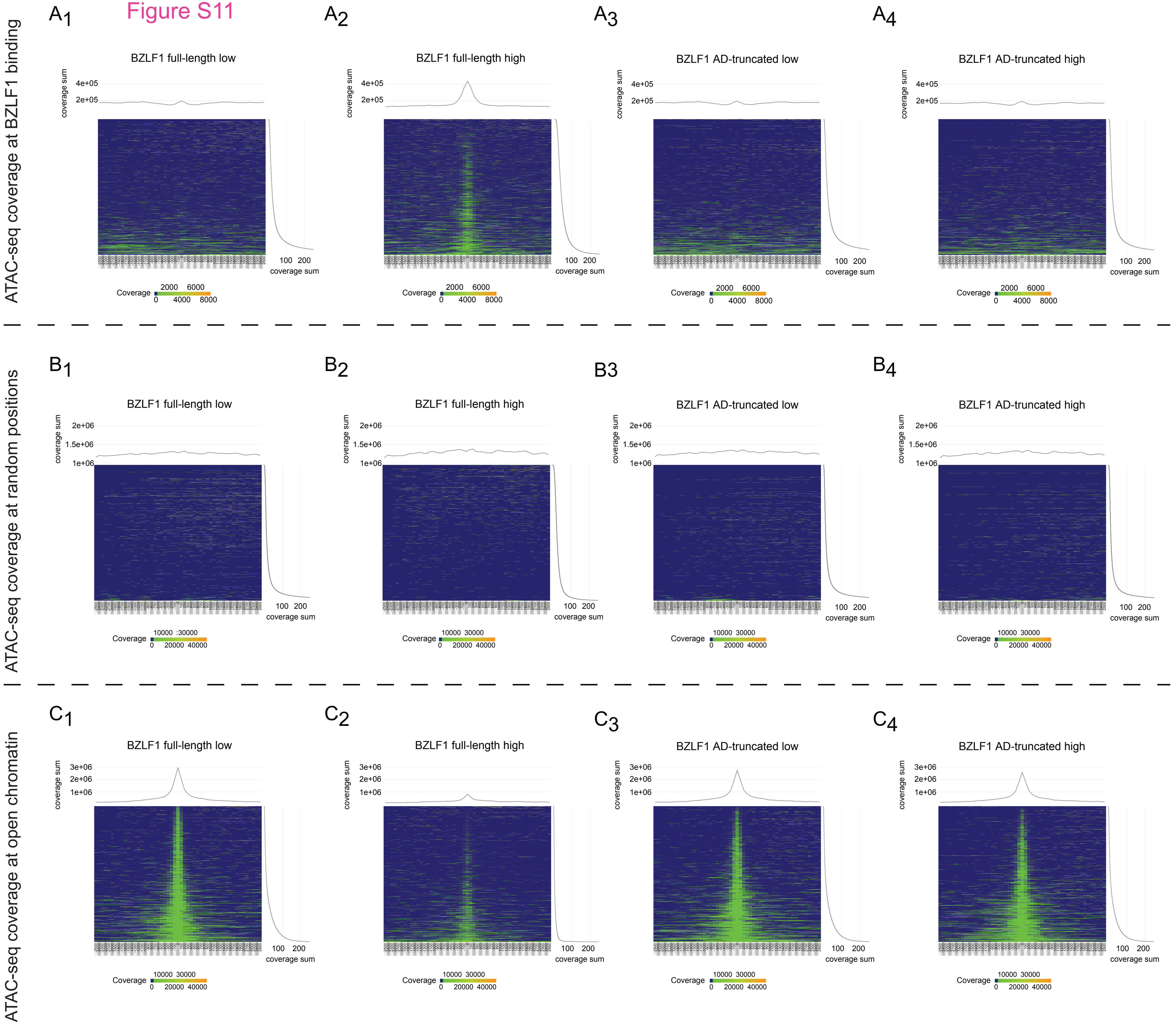
ATAC-seq coverage at BZLF1 binding
A1
BZLF1 full-length low
A2
BZLF1 full-length high
A3
BZLF1 AD-truncated low
A4
BZLF1 AD-truncated high
coverage sum
4e+05
2e+05
100 200
coverage sum
Coverage
2000 6000
0 4000 8000
ATAC-seq coverage at random positions
B1
BZLF1 full-length low
B2
BZLF1 full-length high
B3
BZLF1 AD-truncated low
B4
BZLF1 AD-truncated high
2e+06
1.5e+06
1e+06
10000 30000
0 20000 40000
ATAC-seq coverage at open chromatin
C1
BZLF1 full-length low
C2
BZLF1 full-length high
C3
BZLF1 AD-truncated low
C4
BZLF1 AD-truncated high
3e+06
2e+06
1e+06

### Figure S12

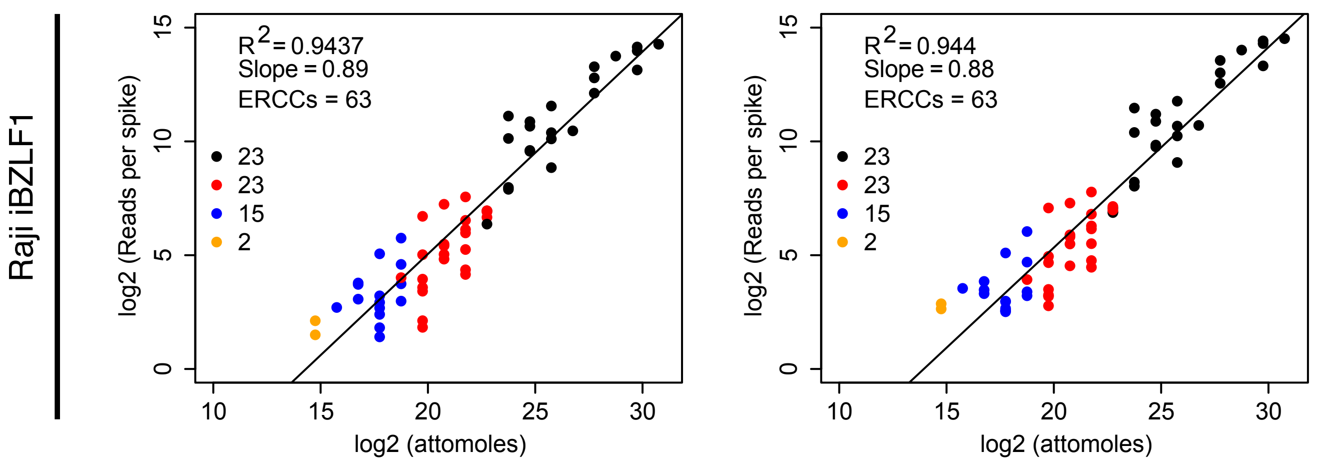
Raji iBZLF1
0 h
6 h
R2= 0.9437
Slope = 0.89
ERCCs = 63
R2= 0.944
Slope = 0.88
ERCCs = 63
23
23
15
2
log2 (Reads per spike)
log2 (attomoles)
0
5
10
15
10
15
20
25
30

### Figure S13

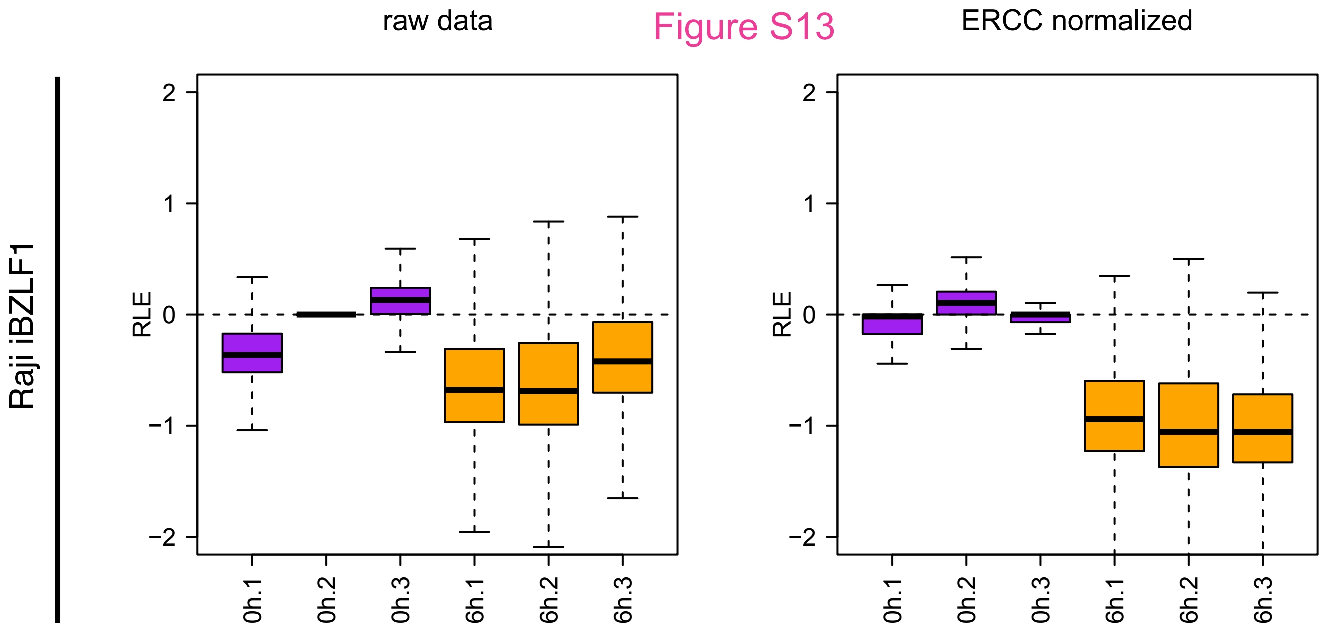
raw data
ERCC normalized
Raji iBZLF1
RLE
2
1
0
−1
−2
0h.1
0h.2
0h.3
6h.1
6h.2
6h.3

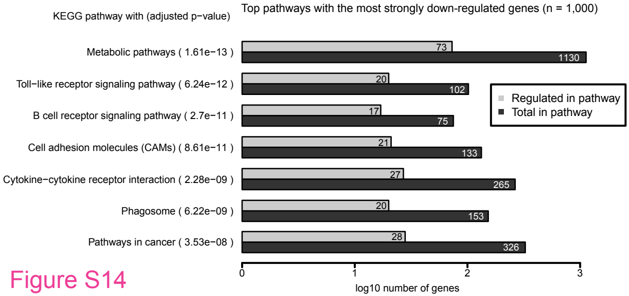
KEGG pathway with (adjusted p−value)
Top pathways with the most strongly down-regulated genes (n = 1,000)
Metabolic pathways ( 1.61e−13 )
73
1130
Toll−like receptor signaling pathway ( 6.24e−12 )
20
102
B cell receptor signaling pathway ( 2.7e−11 )
17
75
Cell adhesion molecules (CAMs) ( 8.61e−11 )
21
133
Cytokine−cytokine receptor interaction ( 2.28e−09 )
27
265
Phagosome ( 6.22e−09 )
20
153
Pathways in cancer ( 3.53e−08 )
28
326
Regulated in pathway
Total in pathway
0
1
2
3
log10 number of genes

Figure S14

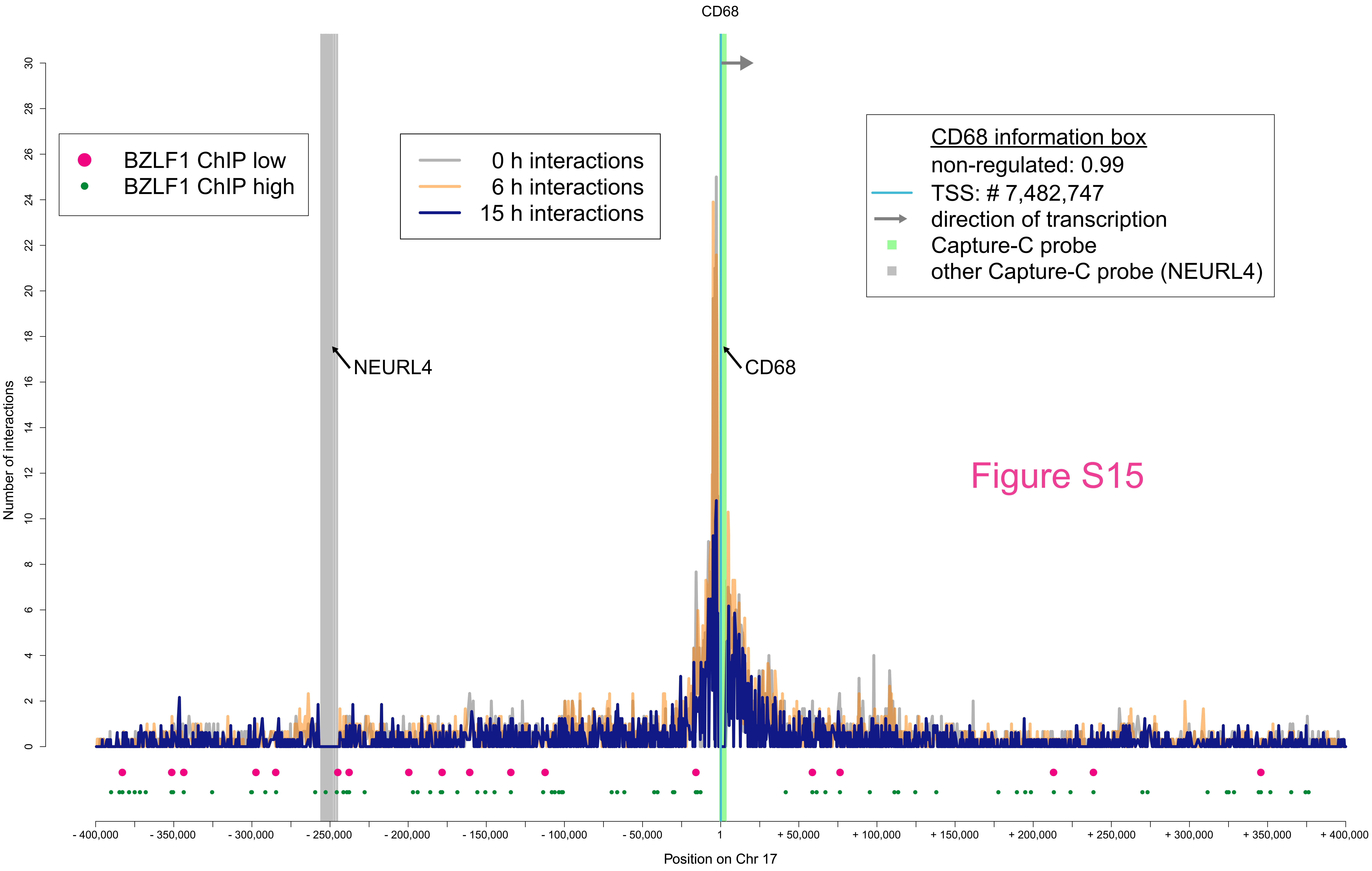
CD68
BZLF1 ChIP low
BZLF1 ChIP high
0 h interactions
6 h interactions
15 h interactions
CD68 information box
non-regulated: 0.99
TSS: # 7,482,747
direction of transcription
Capture-C probe
other Capture-C probe (NEURL4)
NEURL4
CD68
Number of interactions
0
2
4
6
8
10
12
14
16
18
20
22
24
26
28
30
- 400,000
- 350,000
- 300,000
- 250,000
- 200,000
- 150,000
- 100,000
- 50,000
1
+ 50,000
+ 100,000
+ 150,000
+ 200,000
+ 250,000
+ 300,000
+ 350,000
+ 400,000
Position on Chr 17

Figure S15

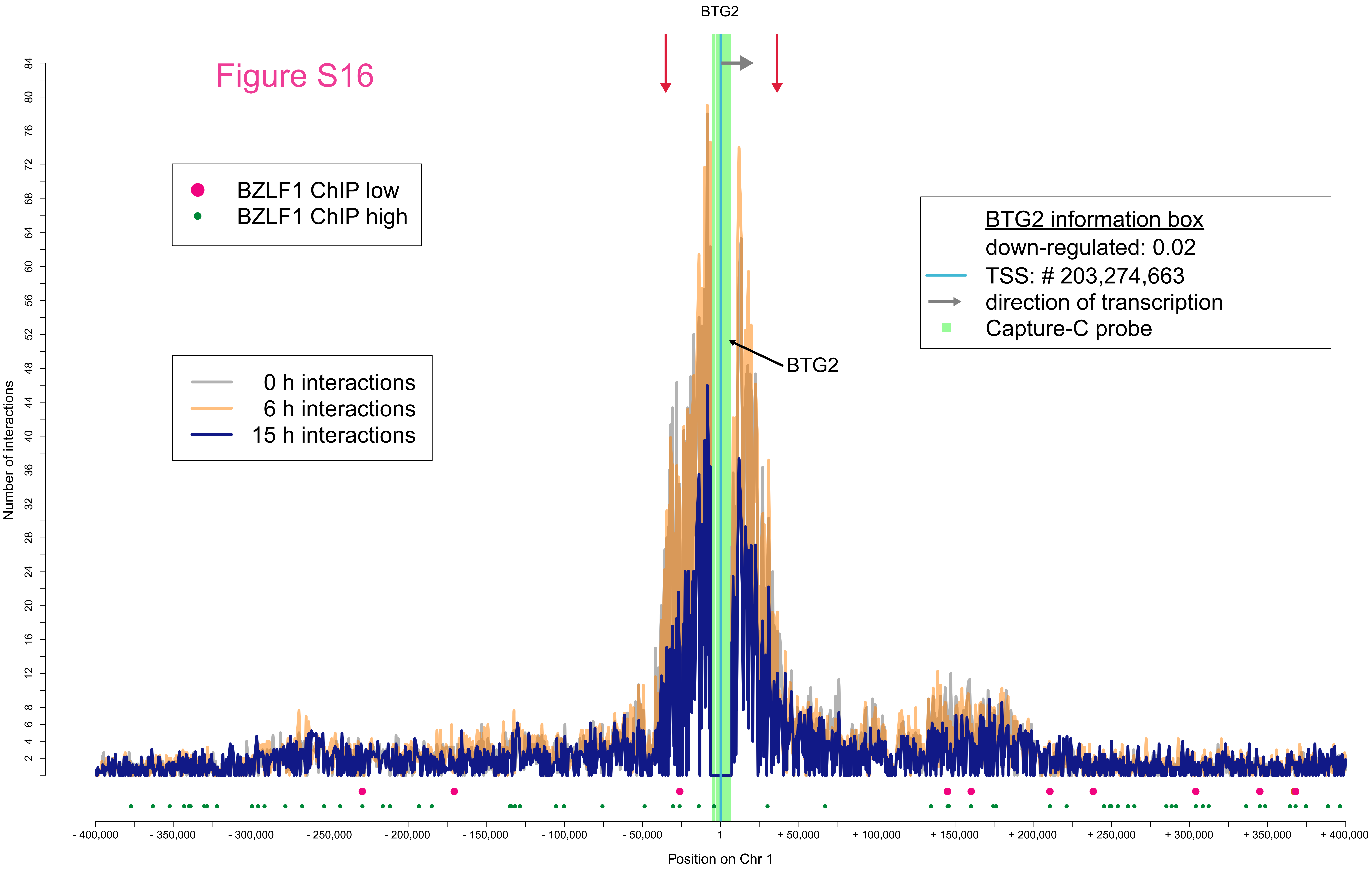
Figure S16
BTG2
BZLF1 ChIP low
BZLF1 ChIP high
0 h interactions
6 h interactions
15 h interactions
BTG2 information box
down-regulated: 0.02
TSS: # 203,274,663
direction of transcription
Capture-C probe
BTG2
Number of interactions
Position on Chr 1
- 400,000
- 350,000
- 300,000
- 250,000
- 200,000
- 150,000
- 100,000
- 50,000
1
+ 50,000
+ 100,000
+ 150,000
+ 200,000
+ 250,000
+ 300,000
+ 350,000
+ 400,000

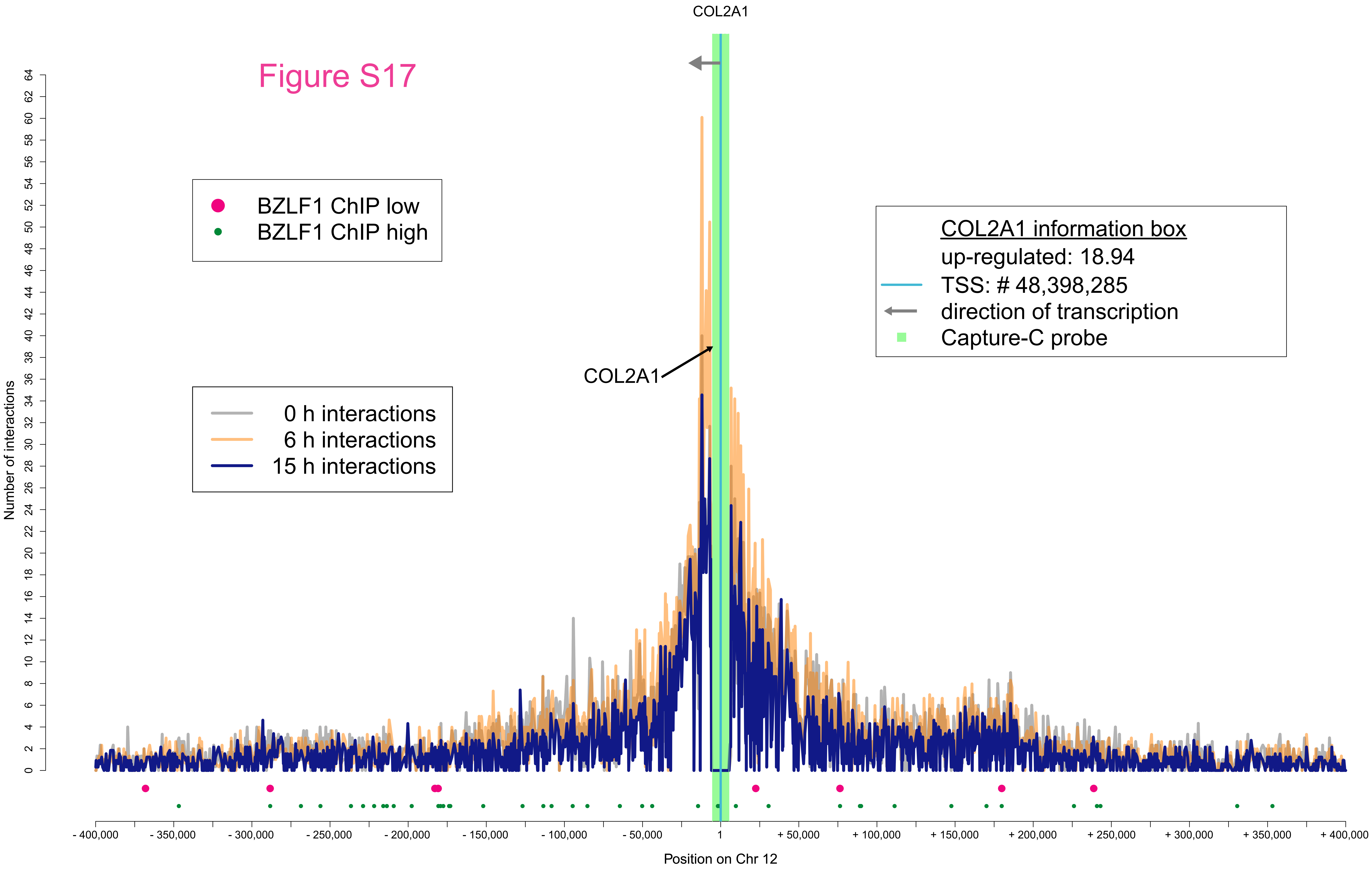

Figure S17
COL2A1
BZLF1 ChIP low
BZLF1 ChIP high
0 h interactions
6 h interactions
15 h interactions
COL2A1 information box
up-regulated: 18.94
TSS: # 48,398,285
direction of transcription
Capture-C probe
Number of interactions
Position on Chr 12
- 400,000
- 350,000
- 300,000
- 250,000
- 200,000
- 150,000
- 100,000
- 50,000
1
+ 50,000
+ 100,000
+ 150,000
+ 200,000
+ 250,000
+ 300,000
+ 350,000
+ 400,000

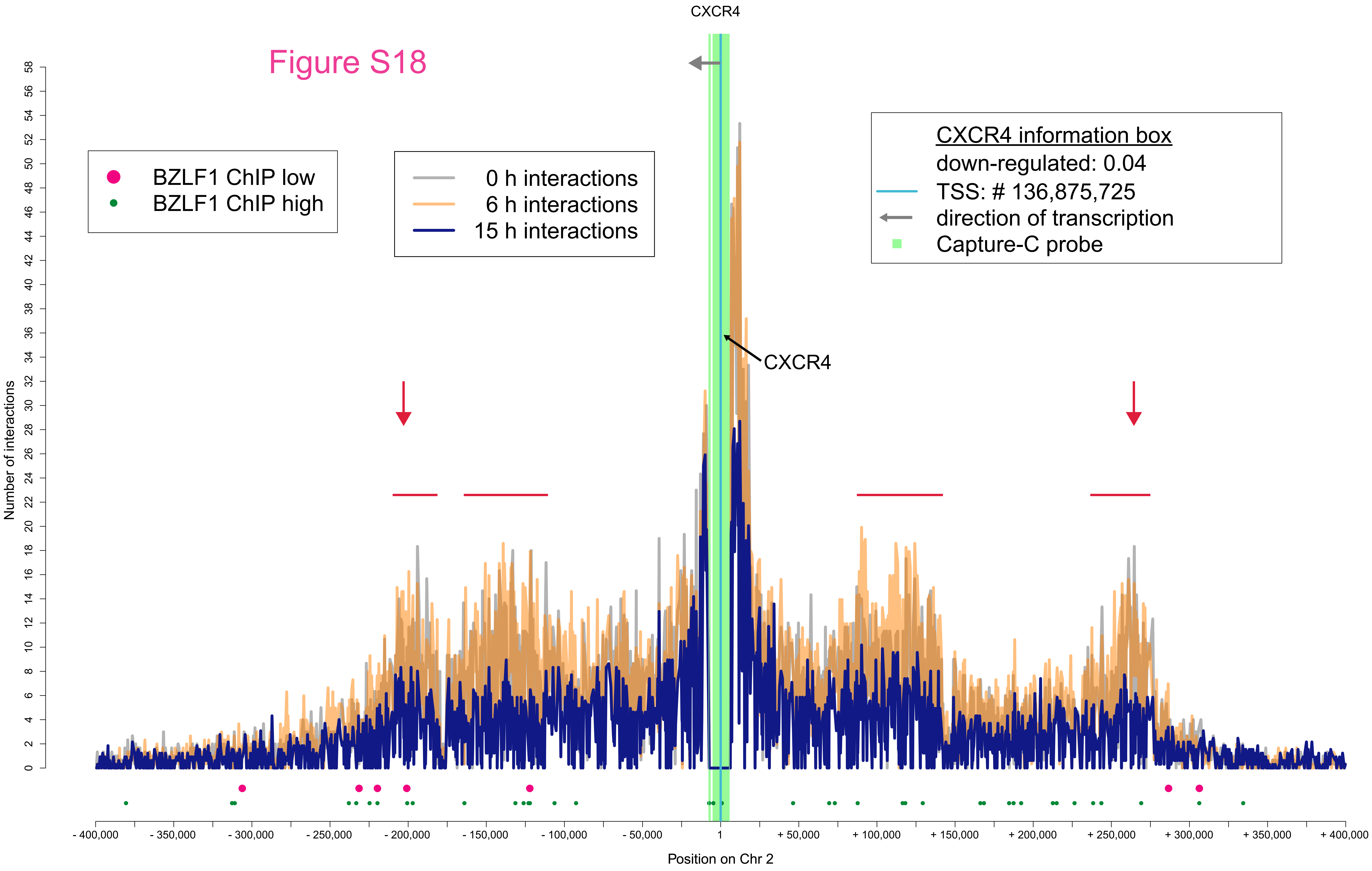

CXCR4
Figure S18
BZLF1 ChIP low
BZLF1 ChIP high
0 h interactions
6 h interactions
15 h interactions
CXCR4 information box
down-regulated: 0.04
TSS: # 136,875,725
direction of transcription
Capture-C probe
CXCR4
Number of interactions
0
2
4
6
8
10
12
14
16
18
20
22
24
26
28
30
32
34
36
38
40
42
44
46
48
50
52
54
56
58
- 400,000
- 350,000
- 300,000
- 250,000
- 200,000
- 150,000
- 100,000
- 50,000
1
+ 50,000
+ 100,000
+ 150,000
+ 200,000
+ 250,000
+ 300,000
+ 350,000
+ 400,000
Position on Chr 2

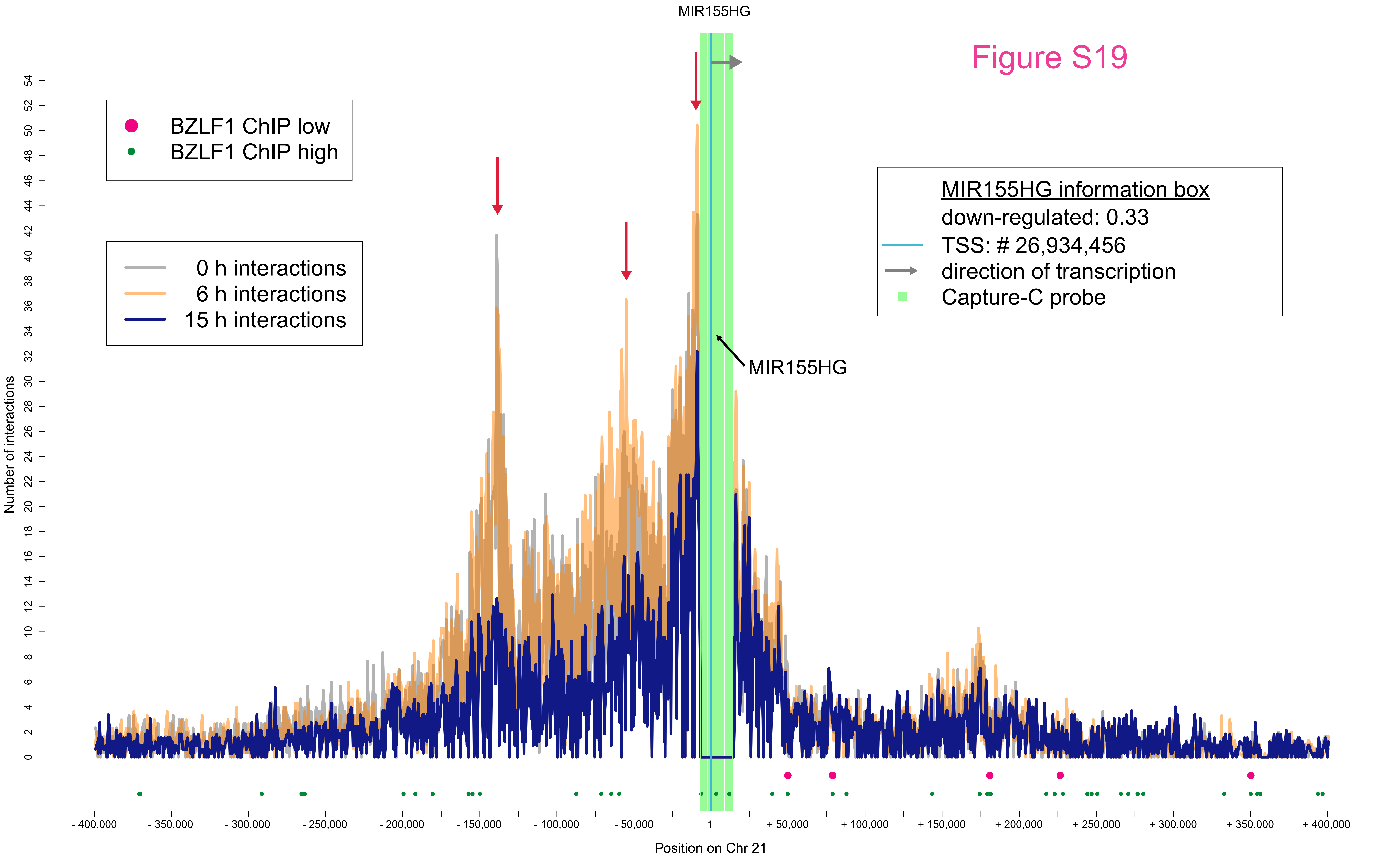

MIR155HG
Figure S19
BZLF1 ChIP low
BZLF1 ChIP high
0 h interactions
6 h interactions
15 h interactions
MIR155HG information box
down-regulated: 0.33
TSS: # 26,934,456
direction of transcription
Capture-C probe
MIR155HG
Number of interactions
0
2
4
6
8
10
12
14
16
18
20
22
24
26
28
30
32
34
36
38
40
42
44
46
48
50
52
54
- 400,000
- 350,000
- 300,000
- 250,000
- 200,000
- 150,000
- 100,000
- 50,000
1
+ 50,000
+ 100,000
+ 150,000
+ 200,000
+ 250,000
+ 300,000
+ 350,000
+ 400,000
Position on Chr 21

### Figure S20

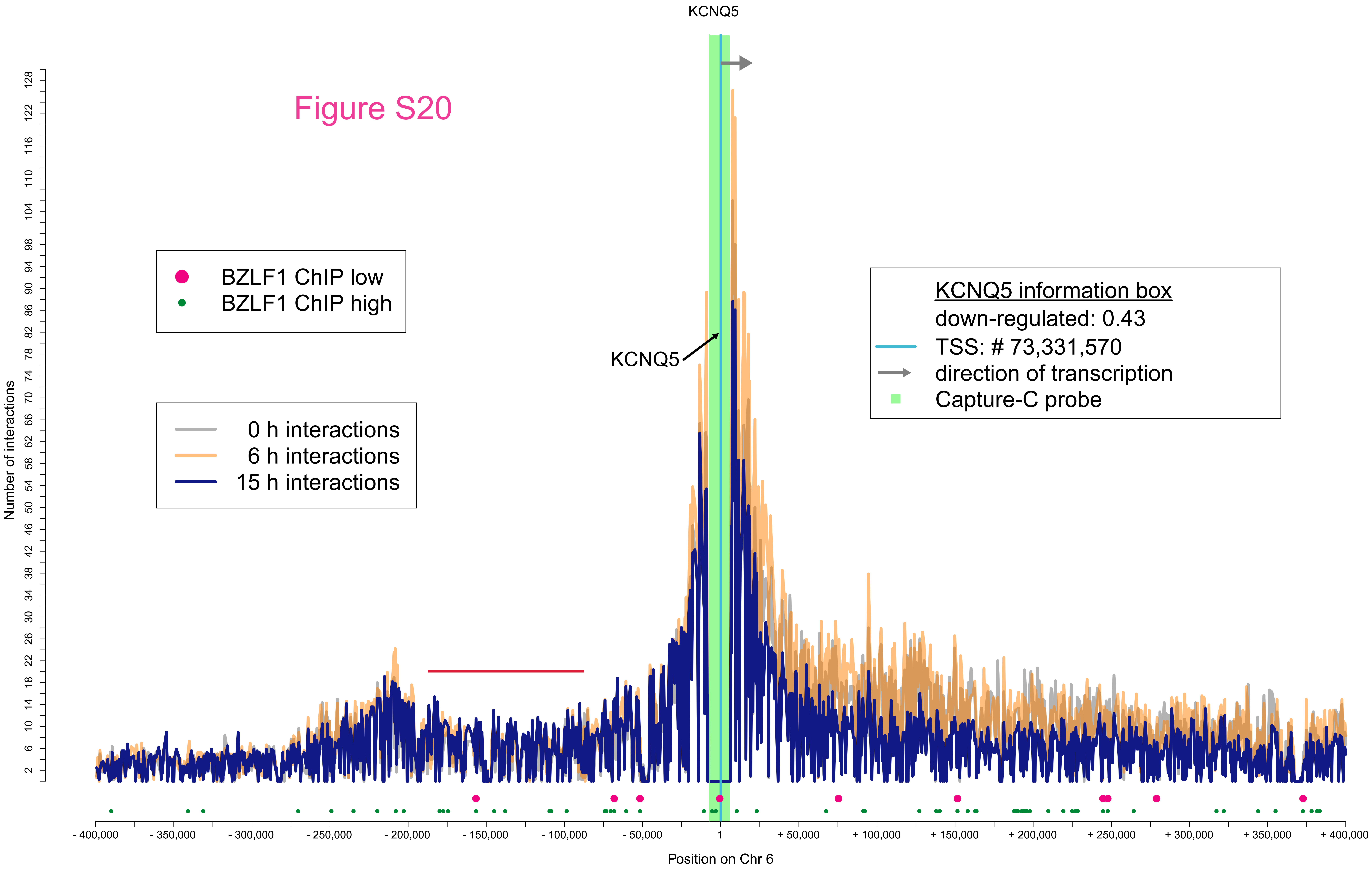

KCNQ5
BZLF1 ChIP low
BZLF1 ChIP high
0 h interactions
6 h interactions
15 h interactions
KCNQ5
KCNQ5 information box
down-regulated: 0.43
TSS: # 73,331,570
direction of transcription
Capture-C probe
Number of interactions
Position on Chr 6
- 400,000
- 350,000
- 300,000
- 250,000
- 200,000
- 150,000
- 100,000
- 50,000
1
+ 50,000
+ 100,000
+ 150,000
+ 200,000
+ 250,000
+ 300,000
+ 350,000
+ 400,000

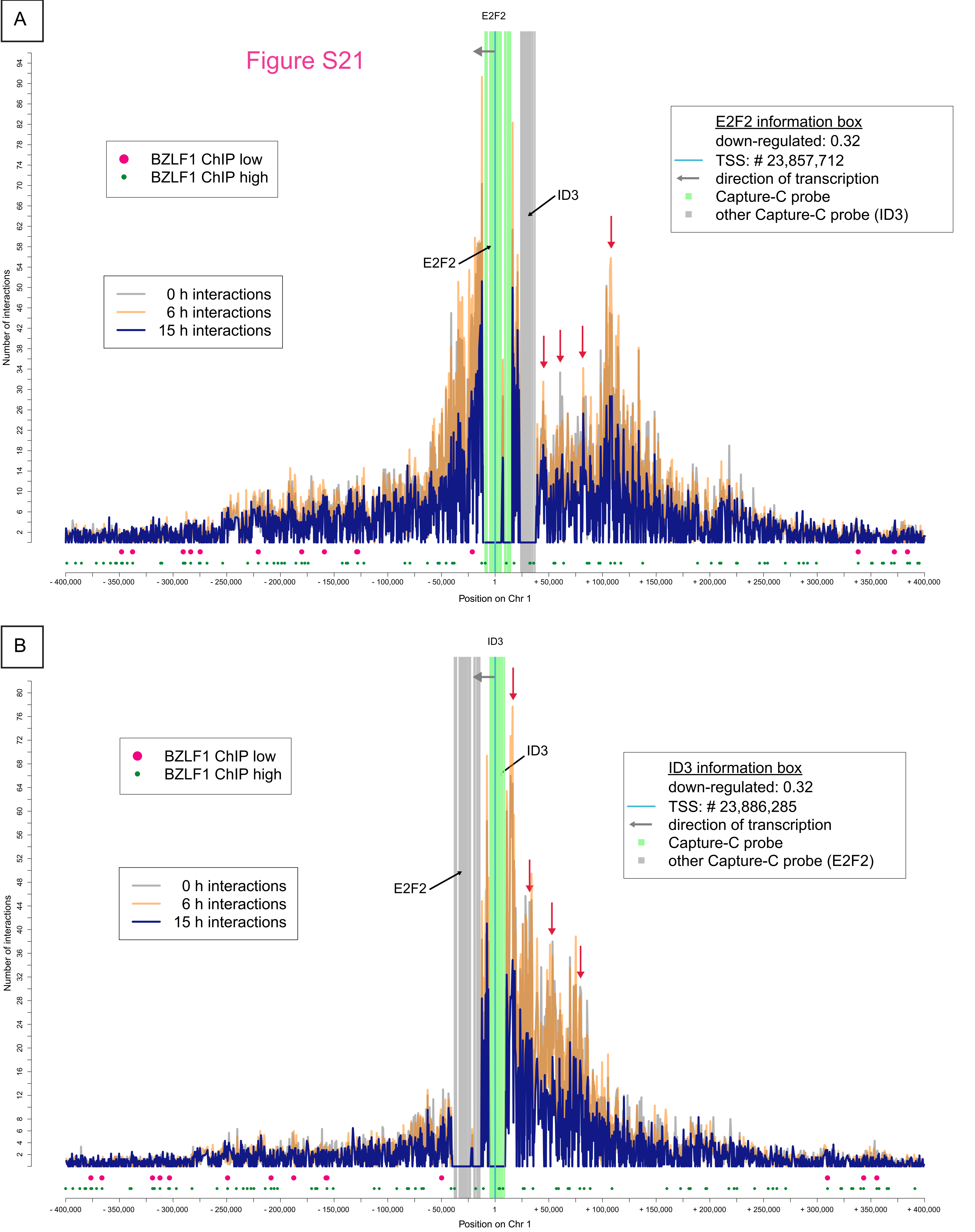

A
Figure S21
E2F2
BZLF1 ChIP low
BZLF1 ChIP high
0 h interactions
6 h interactions
15 h interactions
E2F2 information box
down-regulated: 0.32
TSS: # 23,857,712
direction of transcription
Capture-C probe
other Capture-C probe (ID3)
ID3
E2F2
Number of interactions
Position on Chr 1
B
ID3
BZLF1 ChIP low
BZLF1 ChIP high
ID3 information box
down-regulated: 0.32
TSS: # 23,886,285
direction of transcription
Capture-C probe
other Capture-C probe (E2F2)
ID3
E2F2
0 h interactions
6 h interactions
15 h interactions
Number of interactions
Position on Chr 1

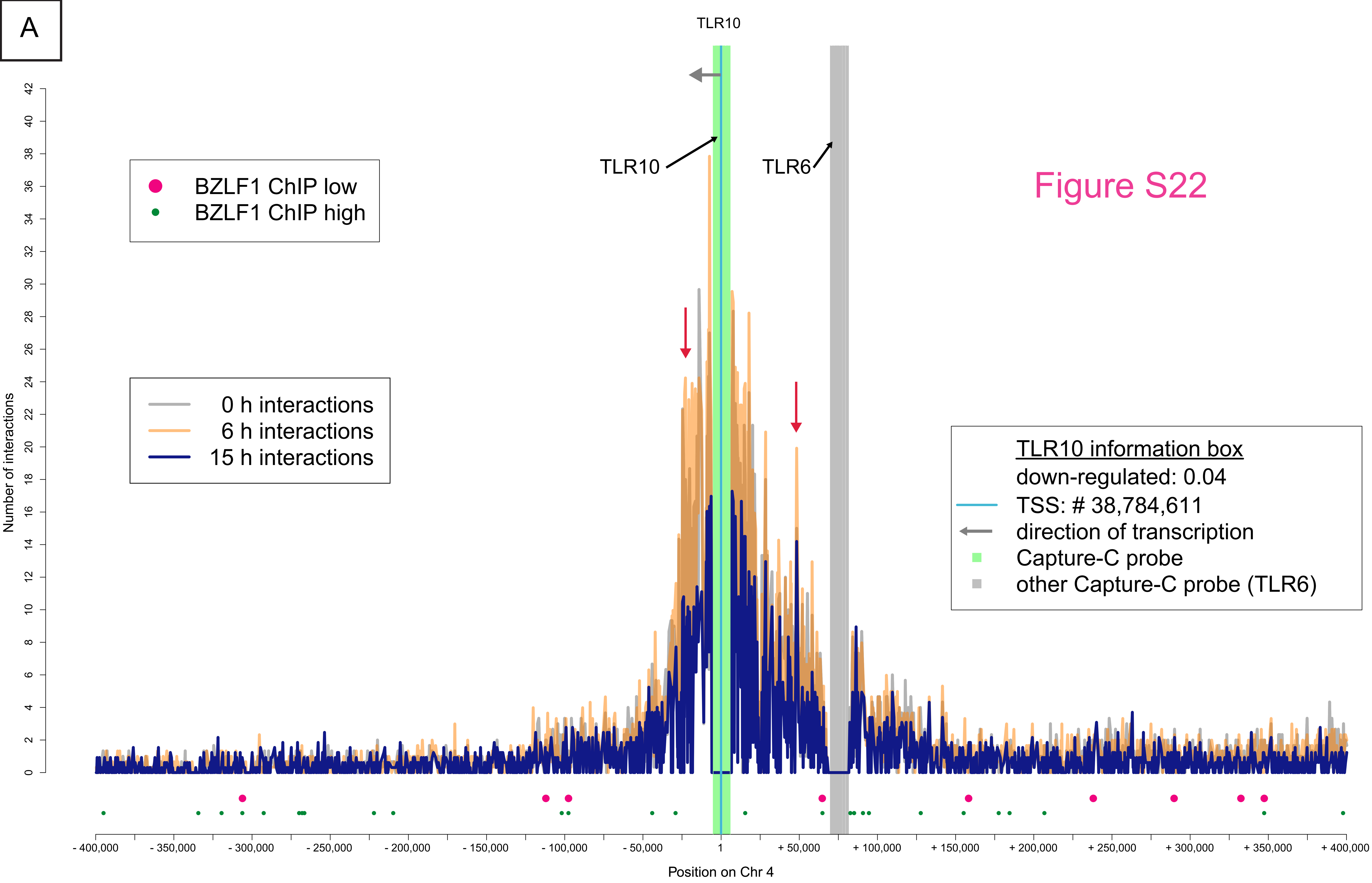
A
TLR10
TLR10
TLR6
Figure S22
BZLF1 ChIP low
BZLF1 ChIP high
0 h interactions
6 h interactions
15 h interactions
TLR10 information box
down-regulated: 0.04
TSS: # 38,784,611
direction of transcription
Capture-C probe
other Capture-C probe (TLR6)
Number of interactions
Position on Chr 4

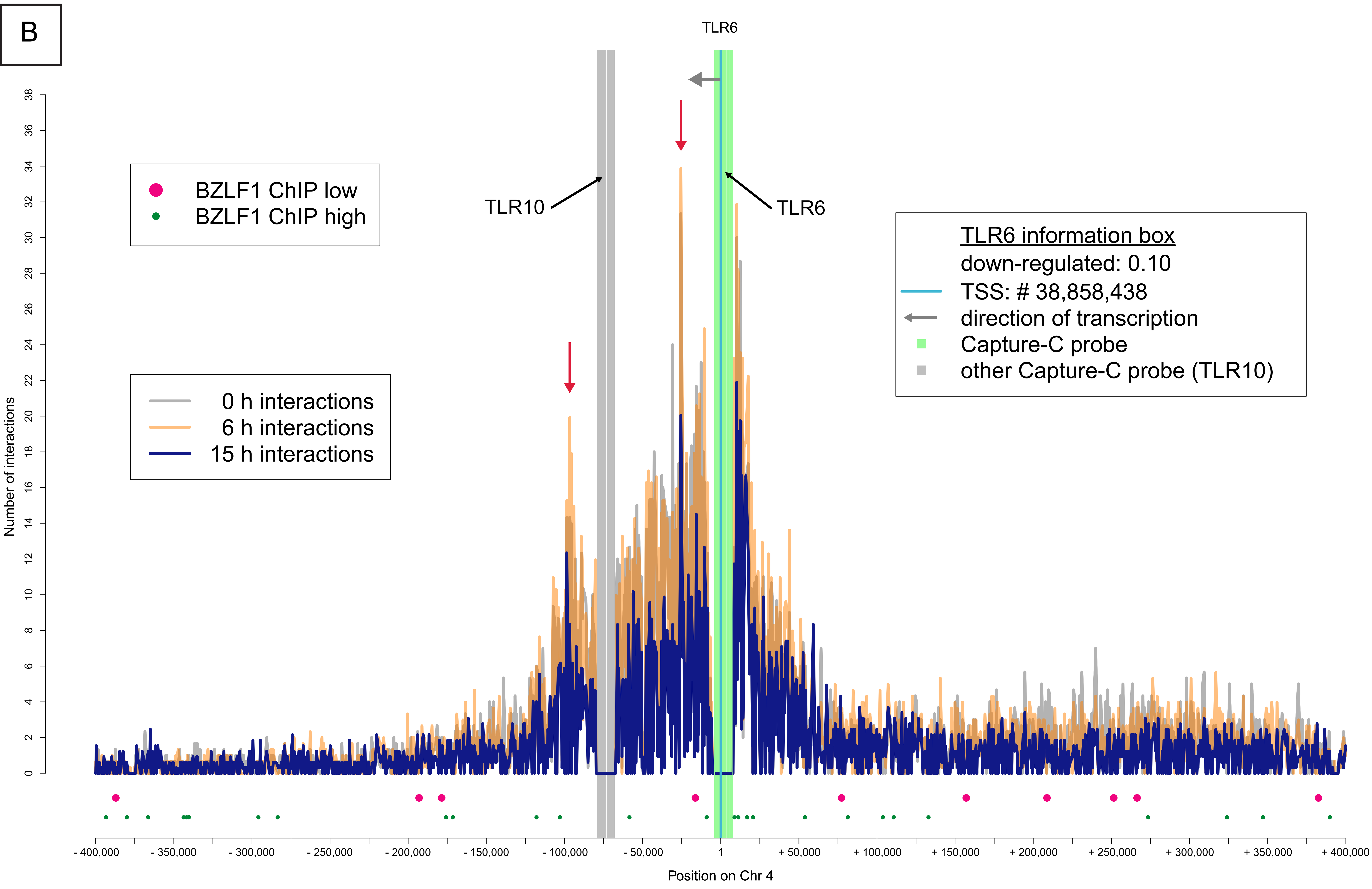
B
TLR6
TLR10
TLR6
BZLF1 ChIP low
BZLF1 ChIP high
0 h interactions
6 h interactions
15 h interactions
TLR6 information box
down-regulated: 0.10
TSS: # 38,858,438
direction of transcription
Capture-C probe
other Capture-C probe (TLR10)
Number of interactions
Position on Chr 4

A

Figure S23

ctcatggacccccggacaccagtcccacccTGTGTCTgtgATGCTCActctcTGAGCCTtccctg
accccaTGTGCCAaagcctggcccagcaatgccacgTGTGCATGTGTGAGCGTgagcttggc
agcgggtgggagaatgTGTGCGTcagTGTGTGAGCGTgagcacggcctgtgcttgtctggctgtgg
gactgggggcctctgtgacgctgtgagcccttgtggaccTGAGTGTgggccgtgtgagctgtgtgcacgtgg
ctgtggcTGAGTGTGAGCCTcTGAGTGTGTGTGTCTgtgccgtgtacctcTGTGTGT

B

| | CpG | | Non-CpG | | |
|---|---|---|---|---|---|
| | 1 x | 2 x | 1 x | 2 x | 3 x |
| A-ending | | | TGTGCCA | TGTGTGA | |
| T-ending | TGTGCGT | TGAGCGT | TGAGCAT | TGAGCCT | TGAGTGT |
| | | | TGTGCAT | TGTGTCT | TGTGTGT |

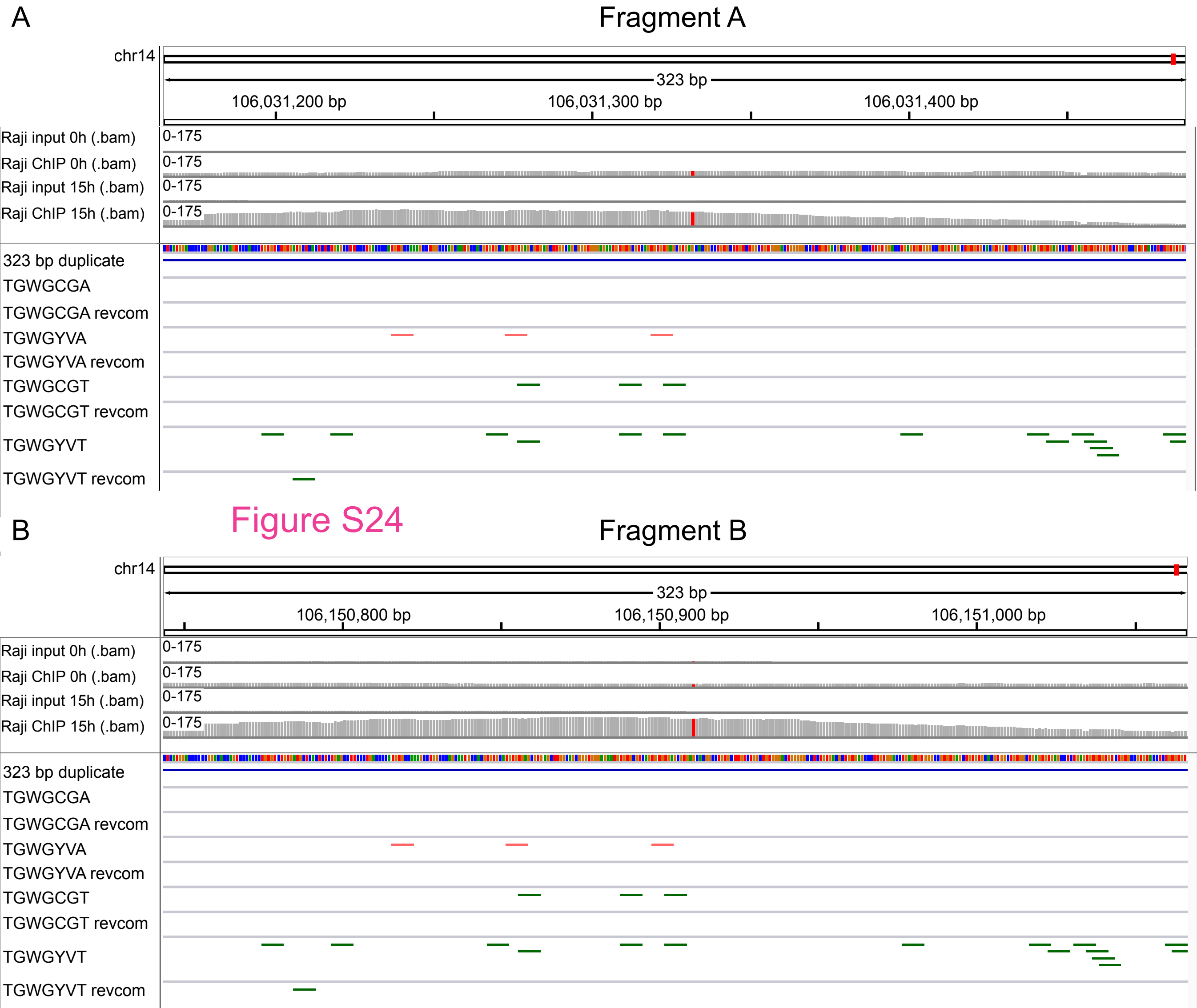
A
Fragment A
chr14
323 bp
106,031,200 bp
106,031,300 bp
106,031,400 bp
Raji input 0h (.bam)
Raji ChIP 0h (.bam)
Raji input 15h (.bam)
Raji ChIP 15h (.bam)
0-175
323 bp duplicate
TGWGCGA
TGWGCGA revcom
TGWGYVA
TGWGYVA revcom
TGWGCGT
TGWGCGT revcom
TGWGYVT
TGWGYVT revcom
B
Fragment B
chr14
323 bp
106,150,800 bp
106,150,900 bp
106,151,000 bp
Raji input 0h (.bam)
Raji ChIP 0h (.bam)
Raji input 15h (.bam)
Raji ChIP 15h (.bam)
0-175
323 bp duplicate
TGWGCGA
TGWGCGA revcom
TGWGYVA
TGWGYVA revcom
TGWGCGT
TGWGCGT revcom
TGWGYVT
TGWGYVT revcom

Figure S24

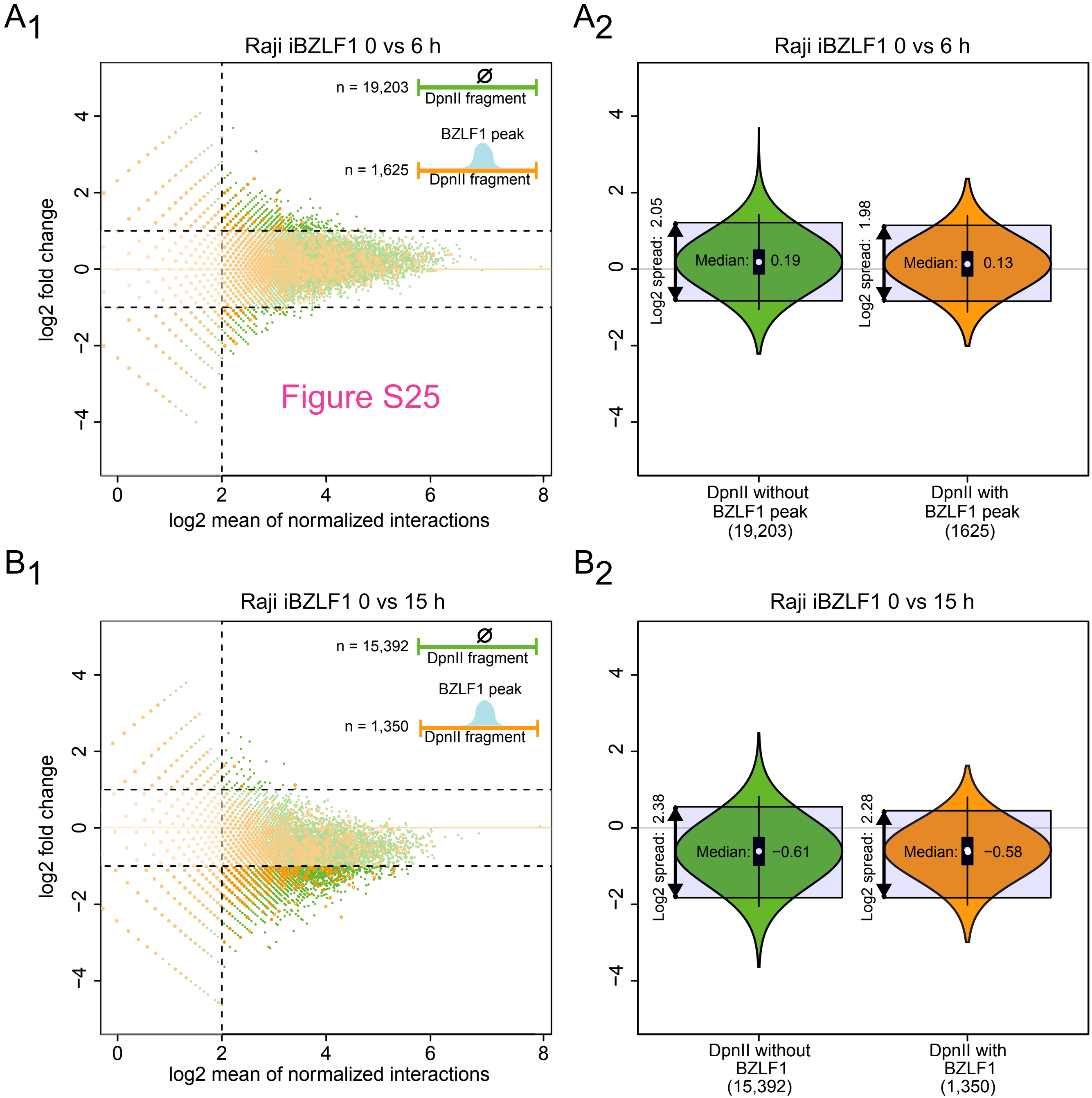
A1
Raji iBZLF1 0 vs 6 h
n = 19,203
Ø
DpnII fragment
BZLF1 peak
n = 1,625
DpnII fragment
Figure S25
log2 fold change
log2 mean of normalized interactions
A2
Raji iBZLF1 0 vs 6 h
Log2 spread: 2.05
Median: 0.19
Log2 spread: 1.98
Median: 0.13
DpnII without BZLF1 peak (19,203)
DpnII with BZLF1 peak (1625)
B1
Raji iBZLF1 0 vs 15 h
n = 15,392
Ø
DpnII fragment
BZLF1 peak
n = 1,350
DpnII fragment
log2 fold change
log2 mean of normalized interactions
B2
Raji iBZLF1 0 vs 15 h
Log2 spread: 2.38
Median: −0.61
Log2 spread: 2.28
Median: −0.58
DpnII without BZLF1 (15,392)
DpnII with BZLF1 (1,350)
