## Supplementary Tables S1-S3 for "Epstein-Barr virus inactivates the transcriptome and disrupts the chromatin architecture of its host cell in the first phase of lytic reactivation"

Supplementary Table S1

**Viral gene expression prior to and after induction of EBV's lytic cycle.**

| Gene_symbol [a] | log2FoldChange [b] | baseMean [c] | lfcSE [d] | padj [e] | PriorDox [f] | PostDox [g] |
|---|---|---|---|---|---|---|
| **EBER-1** | -0.34 | 47509 | 0.2 | 2.90E-01 | 53081 | 41937 |
| **EBNA-LP** | -1.08 | 118 | 0.3 | 1.00E-03 | 160 | 76 |
| **EBNA1** | -1.48 | 1876 | 0.3 | 1.8e-06 | 2762 | 990 |
| **EBNA2** | -1.83 | 437 | 0.3 | 8.3e-11 | 682 | 192 |
| **BXRF1** | 4.69 | 24 | 0.7 | 6.7e-10 | 2 | 46 |
| **RPMS1** | 3.56 | 70 | 0.4 | 8.00E-16 | 11 | 129 |
| **BDLF2** | 0.94 | 19 | 0.5 | 7.7e-2 | 13 | 25 |
| **BDLF1** | 0.92 | 8 | 0.6 | 2.80E-01 | 6 | 10 |
| **EBER-2** | 0.44 | 439 | 0.3 | 2.90E-01 | 373 | 505 |
| **BVLF1** | 4.34 | 366 | 0.5 | 3.7e-19 | 34 | 698 |
| **BILF2** | 3.94 | 569 | 0.5 | 2.7e-17 | 70 | 1068 |
| **BALF3** | 3.93 | 209 | 0.4 | 3.5e-21 | 26 | 392 |
| **BPLF1** | 3.84 | 409 | 0.5 | 3.1e-14 | 53 | 765 |
| **BcLF1** | 2.43 | 198 | 0.5 | 1.4e-06 | 62 | 334 |
| **BTRF1** | 6.42 | 377 | 0.4 | 3.3e-56 | 9 | 745 |
| **BRRF2** | 5.99 | 697 | 0.3 | 2.1e-80 | 22 | 1372 |
| **BLRF1** | 5.94 | 468 | 0.4 | 6.8e-48 | 15 | 921 |
| **BARF0** | 5.72 | 1302 | 0.4 | 1.4e-57 | 48 | 2556 |
| **BLRF2** | 5.72 | 537 | 0.4 | 5.1e-57 | 20 | 1054 |
| **BBRF2** | 5.42 | 279 | 0.4 | 9.4e-43 | 13 | 545 |
| **BSRF1** | 5.41 | 1348 | 0.3 | 6.5e-55 | 62 | 2634 |
| **LMP2** | 5.3 | 726 | 0.6 | 2.8e-17 | 36 | 1416 |
| **BVRF2** | 5.3 | 2167 | 0.4 | 5.8e-41 | 107 | 4227 |
| **BVRF1** | 5.24 | 761 | 0.4 | 4.5e-31 | 39 | 1483 |
| **BBRF1** | 4.78 | 537 | 0.3 | 5.8e-63 | 38 | 1036 |
| **BaRF1** | 4.64 | 1904 | 0.5 | 7.4e-22 | 147 | 3661 |
| **BDLF3.5** | 8.18 | 1579 | 0.3 | 1.3e-127 | 11 | 3147 |
| **BBLF3-BBLF2** | 7.97 | 24994 | 0.3 | 4.00E-159 | 199 | 49789 |
| **BDLF4** | 7.78 | 622 | 0.5 | 6.2e-59 | 6 | 1238 |
| **BBLF1** | 7.76 | 5347 | 0.5 | 1.8e-65 | 49 | 10645 |
| **BZLF1** | 7.7 | 28019 | 0.3 | 1.7e-168 | 268 | 55770 |
| **BGLF4** | 7.56 | 17984 | 0.3 | 1.5e-192 | 190 | 35778 |
| **BLLF3** | 7.52 | 957 | 0.4 | 3.2e-89 | 10 | 1904 |
| **BBLF4** | 7.45 | 19055 | 0.3 | 9.4e-102 | 217 | 37893 |
| **BGLF5** | 7.4 | 43750 | 0.4 | 2.2e-96 | 515 | 86985 |
| **BGLF3.5** | 7.31 | 8438 | 0.2 | 1.3e-274 | 106 | 16770 |
| **BLLF1-BLLF2** | 7.28 | 4401 | 0.3 | 3.1e-134 | 56 | 8746 |

| | | | | | | |
|---|---|---|---|---|---|---|
| **BFLF1** | 7.06 | 11569 | 0.3 | 4.00E-98 | 172 | 22966 |
| **BGLF2** | 7.02 | 2092 | 0.3 | 8.00E-95 | 32 | 4152 |
| **BKRF2** | 6.96 | 6110 | 0.3 | 1.00E-127 | 97 | 12123 |
| **LF1** | 6.81 | 6597 | 0.4 | 2.7e-69 | 117 | 13077 |
| **BGLF3** | 6.8 | 2066 | 0.3 | 1.2e-110 | 37 | 4095 |
| **BILF1** | 6.8 | 10214 | 0.4 | 4.7e-59 | 182 | 20246 |
| **BALF5** | 6.78 | 25395 | 0.4 | 5.2e-74 | 458 | 50332 |
| **BKRF3** | 6.76 | 8630 | 0.3 | 3.3e-91 | 158 | 17102 |
| **BFRF1A** | 6.63 | 9032 | 0.3 | 1.5e-113 | 181 | 17883 |
| **BKRF4** | 6.63 | 10577 | 0.3 | 5.8e-93 | 211 | 20943 |
| **BRRF1** | 6.58 | 2114 | 0.4 | 2.7e-62 | 44 | 4184 |
| **BFRF3** | 6.57 | 7214 | 0.4 | 1.8e-64 | 150 | 14278 |
| **BORF1** | 6.56 | 36895 | 0.5 | 1.8e-46 | 774 | 73016 |
| **BORF2** | 6.53 | 56450 | 0.5 | 5.6e-44 | 1209 | 111691 |
| **BFRF1** | 6.45 | 6153 | 0.4 | 1.1e-68 | 139 | 12167 |
| **BRLF1** | 6.44 | 6881 | 0.4 | 1.9e-58 | 157 | 13605 |
| **BFRF2** | 6.39 | 9496 | 0.4 | 3.8e-60 | 224 | 18768 |
| **LF2** | 6.3 | 12876 | 0.4 | 1.2e-52 | 323 | 25429 |
| **BBRF3** | 6.27 | 3659 | 0.3 | 3.6e-118 | 94 | 7224 |
| **BGRF1/BDRF1** | 6.17 | 3591 | 0.3 | 3.4e-121 | 98 | 7084 |
| **BALF4** | 6.13 | 4510 | 0.4 | 3.9e-54 | 127 | 8893 |
| **BMRF2** | 6.13 | 59620 | 0.5 | 2.4e-40 | 1679 | 117561 |
| **BMLF1-BSLF1** | 6.12 | 63992 | 0.4 | 2.7e-48 | 1814 | 126170 |
| **BMRF1** | 5.94 | 46694 | 0.4 | 3.2e-50 | 1497 | 91891 |
| **LF3** | 5.9 | 7020 | 0.4 | 1.4e-46 | 231 | 13809 |
| **BXLF1** | 5.84 | 26640 | 0.4 | 3.5e-40 | 914 | 52366 |
| **BXLF2** | 5.8 | 24431 | 0.4 | 1.3e-39 | 862 | 48000 |
| **BNLF2b** | 5.13 | 5796 | 0.4 | 9.00E-40 | 322 | 11270 |
| **BHLF1** | 5.05 | 72803 | 0.3 | 1.5e-51 | 4266 | 141340 |
| **BNLF2a** | 5.04 | 11081 | 0.4 | 2.2e-32 | 654 | 21508 |
| **BHRF1** | 4.85 | 38594 | 0.3 | 2.1e-65 | 2587 | 74601 |

[a] Acronyms of EBV genes.

[b] Log2 transformed comparison between 0 and 6 hours of BZLF1 induction.

[c] Average expression level.

[d] Log2 transformed fold change standard error.

[e] Adjusted p-value.

[f] Expression level prior to doxycycline induction.

[g] Expression level post doxycycline induction.

Supplementary Table S2

**Cellular genes analyzed in Capture-C experiments, their annotations and features.**

| Gene_symbol [a] | Capture.start [b] | Capture.end [c] | TSS [d] | Chr [e] | Gene.start [f] | Gene.end [g] |
|---|---|---|---|---|---|---|
| AMPD3 | 10,468,690 | 10,481,379 | 10,476,665 | chr11 | 10,472,224 | 10,529,126 |
| APBB3 | 139,938,429 | 139,955,586 | 139,944,189 | chr5 | 139,937,853 | 139,944,189 |
| BTG2 | 203,269,112 | 203,281,166 | 203,274,663 | chr1 | 203,274,664 | 203,278,729 |
| CCDC103 | 42,971,726 | 42,982,744 | 42,977,079 | chr17 | 42,977,080 | 42,981,047 |
| CCR7 | 38,714,766 | 38,734,803 | 38,721,736 | chr17 | 38,710,022 | 38,721,736 |
| CD68 | 7,482,068 | 7,486,280 | 7,482,747 | chr17 | 7,482,805 | 7,485,429 |
| CD79B | 62,002,864 | 62,013,170 | 62,009,704 | chr17 | 62,006,098 | 62,009,704 |
| CEBPB | 48,795,327 | 48,815,340 | 48,807,119 | chr20 | 48,807,120 | 48,809,227 |
| COL2A1 | 48,393,035 | 48,403,596 | 48,398,285 | chr12 | 48,366,748 | 48,398,285 |
| CXCR4 | 136,868,097 | 136,881,255 | 136,875,725 | chr2 | 136,871,919 | 136,875,725 |
| CXCR5 | 118,749,270 | 118,759,566 | 118,754,474 | chr11 | 118,754,475 | 118,766,980 |
| CXCR7 | 237,470,373 | 237,486,379 | 237,478,379 | chr2 | 237,478,380 | 237,490,994 |
| DDR1 | 30,845,183 | 30,863,047 | 30,856,166 | chr6 | 30,856,465 | 30,867,933 |
| DPCR1 | 30,903,809 | 30,914,373 | 30,908,776 | chr6 | 30,908,777 | 30,921,998 |
| E2F2 | 23,847,186 | 23,872,472 | 23,857,712 | chr1 | 23,832,920 | 23,857,712 |
| GPR68 | 91,705,428 | 91,720,645 | 91,710,852 | chr14 | 91,698,876 | 91,710,852 |
| HARS | 140,066,416 | 140,077,508 | 140,071,312 | chr5 | 140,053,490 | 140,071,312 |
| HDAC9 | 18,529,944 | 18,546,880 | 18,535,368 | chr7 | 18,535,885 | 19,036,992 |
| HLA-DQA1 | 32,599,773 | 32,613,124 | 32,605,182 | chr6 | 32,605,183 | 32,611,429 |
| HLA-DRB1 | 32,551,289 | 32,562,299 | 32,557,614 | chr6 | 32,546,547 | 32,557,613 |
| HN1L | 1,722,326 | 1,734,521 | 1,728,277 | chr16 | 1,728,278 | 1,752,073 |
| ID3 | 23,881,142 | 23,895,242 | 23,886,285 | chr1 | 23,884,421 | 23,886,285 |
| IL21R | 27,409,957 | 27,420,674 | 27,413,482 | chr16 | 27,413,483 | 27,460,604 |
| IL7R | 35,851,269 | 35,862,828 | 35,856,976 | chr5 | 35,856,977 | 35,879,705 |
| KANSL2 | 49,071,127 | 49,081,764 | 49,076,035 | chr12 | 49,046,995 | 49,076,008 |
| KCNQ5 | 73,324,319 | 73,337,153 | 73,331,570 | chr6 | 73,331,571 | 73,908,573 |
| KRBA2 | 8,274,263 | 8,287,497 | 8,280,029 | chr17 | 8,271,973 | 8,274,858 |
| LOC100128288 | 8,257,983 | 8,269,101 | 8,263,858 | chr17 | 8,261,731 | 8,263,859 |
| LPIN1 | 11,876,164 | 11,887,089 | 11,881,474 | chr2 | 11,817,705 | 11,967,533 |
| MAPK1 | 22,216,728 | 22,227,300 | 22,221,970 | chr22 | 22,113,947 | 22,221,970 |
| MIR155HG | 26,927,573 | 26,948,545 | 26,934,456 | chr21 | 26,934,457 | 26,947,480 |
| MYC | 128,742,847 | 128,754,169 | 128,748,314 | chr8 | 128,748,315 | 128,753,680 |
| NCF2 | 183,555,420 | 183,567,667 | 183,560,056 | chr1 | 183,524,697 | 183,560,056 |
| NEURL4 | 7,226,693 | 7,237,733 | 7,232,639 | chr17 | 7,218,951 | 7,232,638 |
| NIPBL | 36,869,587 | 36,882,946 | 36,876,860 | chr5 | 36,876,861 | 37,065,921 |
| PKD1 | 2,180,587 | 2,189,279 | 2,185,899 | chr16 | 2,138,711 | 2,185,899 |
| RAB27A | 55,576,845 | 55,590,196 | 55,582,013 | chr15 | 55,495,164 | 55,563,107 |

| | | | | | | |
|---|---|---|---|---|---|---|
| RSPH3 | 159,411,773 | 159,427,250 | 159,421,198 | chr6 | 159,398,266 | 159,421,198 |
| SAAL1 | 18,122,421 | 18,133,014 | 18,127,638 | chr11 | 18,101,890 | 18,127,638 |
| SLC43A3 | 57,188,561 | 57,200,936 | 57,195,054 | chr11 | 57,174,427 | 57,195,053 |
| STRN4 | 47,245,264 | 47,255,897 | 47,250,251 | chr19 | 47,222,768 | 47,249,720 |
| SWAP70 | 9,678,299 | 9,688,213 | 9,685,627 | chr11 | 9,685,628 | 9,774,507 |
| TFEC | 115,791,535 | 115,815,105 | 115,799,950 | chr7 | 115,575,202 | 115,670,867 |
| TIFA | 113,201,569 | 113,213,332 | 113,207,059 | chr4 | 113,196,782 | 113,207,059 |
| TLR10 | 38,779,548 | 38,790,407 | 38,784,611 | chr4 | 38,773,860 | 38,784,611 |
| TLR6 | 38,854,456 | 38,866,131 | 38,858,438 | chr4 | 38,825,329 | 38,858,438 |
| TNF | 31,537,973 | 31,550,047 | 31,543,343 | chr6 | 31,543,344 | 31,546,112 |
| TRIM27 | 28,887,138 | 28,909,405 | 28,891,768 | chr6 | 28,870,779 | 28,891,768 |
| WHSC1L1 | 38,236,292 | 38,245,260 | 38,239,790 | chr8 | 38,132,561 | 38,239,790 |
| ZBTB37 | 173,832,343 | 173,843,173 | 173,837,492 | chr1 | 173,837,493 | 173,855,774 |
| ZNF143 | 9,477,248 | 9,486,800 | 9,482,511 | chr11 | 9,482,512 | 9,550,071 |
| ZNF165 | 28,042,087 | 28,054,278 | 28,048,481 | chr6 | 28,048,482 | 28,057,340 |
| ZSCAN12P1 | 28,055,260 | 28,102,764 | 28,058,584 | chr6 | 28,058,585 | 28,063,493 |

[a] Acronyms of EBV genes.

[b] Nucleotide coordinate of the start of the Capture-C RNA probes.

[c] Nucleotide coordinate of the end of the Capture-C probes.

[d] Transcriptional start sites of the captured genes.

[e] Chromosome of the Capture-C probes.

[f] Top strand nucleotide coordinate of the start of the gene (hg19 genome reference).

[g] Top strand nucleotide coordinate of the end of the gene (hg19 genome reference).

Supplementary Table S3

### Overview of the NGS datasets uploaded to ArrayExpress

| Sample Replicates [a] | ChIP-seq [b] | Accession numbers [c] |
|---|---|---|
| 1 | Raji iBZLF1 full-length ChIP low | E-MTAB-7788 |
| 2 | Raji iBZLF1 full-length ChIP low | E-MTAB-7788 |
| 1 | Raji iBZLF1 full-length ChIP high | E-MTAB-7788 |
| 2 | Raji iBZLF1 full-length ChIP high | E-MTAB-7788 |
| 1 | Raji iBZLF1 full-length input low | E-MTAB-7788 |
| 2 | Raji iBZLF1 full-length input low | E-MTAB-7788 |
| 1 | Raji iBZLF1 full-length input high | E-MTAB-7788 |
| 2 | Raji iBZLF1 full-length input high | E-MTAB-7788 |
| 1 | DG75 parental ChIP | To be added |
| 2 | DG75 parental ChIP | To be added |
| 1 | DG75 parental input | To be added |
| 2 | DG75 parental input | To be added |
| Sample Replicates | ATAC-seq | |
| 1 | Raji iBZLF1 full-length low | E-MTAB-7789 |
| 2 | Raji iBZLF1 full-length low | E-MTAB-7789 |
| 3 | Raji iBZLF1 full-length low | E-MTAB-7789 |
| 1 | Raji iBZLF1 full-length high | E-MTAB-7789 |
| 2 | Raji iBZLF1 full-length high | E-MTAB-7789 |
| 3 | Raji iBZLF1 full-length high | E-MTAB-7789 |
| 1 | Raji iBZLF1 truncated low | E-MTAB-7789 |
| 2 | Raji iBZLF1 truncated low | E-MTAB-7789 |
| 3 | Raji iBZLF1 truncated low | E-MTAB-7789 |
| 1 | Raji iBZLF1 truncated high | E-MTAB-7789 |
| 2 | Raji iBZLF1 truncated high | E-MTAB-7789 |
| 3 | Raji iBZLF1 truncated high | E-MTAB-7789 |
| Sample Replicates | RNA-seq | |
| 1 | Raji iBZLF1 full-length low | To be added |
| 2 | Raji iBZLF1 full-length low | To be added |
| 3 | Raji iBZLF1 full-length low | To be added |
| 1 | Raji iBZLF1 full-length high | To be added |
| 2 | Raji iBZLF1 full-length high | To be added |
| 3 | Raji iBZLF1 full-length high | To be added |
| 1 | Raji iBZLF1 truncated low | To be added |
| 2 | Raji iBZLF1 truncated low | To be added |
| 3 | Raji iBZLF1 truncated low | To be added |
| 1 | Raji iBZLF1 truncated high | To be added |
| 2 | Raji iBZLF1 truncated high | To be added |
| 3 | Raji iBZLF1 truncated high | To be added |
| 1 | Raji parental -dox | To be added |
| 2 | Raji parental -dox | To be added |
| 3 | Raji parental -dox | To be added |
| 1 | Raji parental +dox | To be added |
| 2 | Raji parental +dox | To be added |
| 3 | Raji parental +dox | To be added |
| Samples Sets | Capture-C | |
| 1 | Raji iBZLF1 full-length 0h | To be added |
| 2 | Raji iBZLF1 full-length 0h | To be added |
| 3 | Raji iBZLF1 full-length 0h | To be added |

| | | |
|---|---|---|
| 1 | Raji iBZLF1 full-length 6h | To be added |
| 2 | Raji iBZLF1 full-length 6h | To be added |
| 3 | Raji iBZLF1 full-length 6h | To be added |
| 1 | Raji iBZLF1 full-length 15h | To be added |
| 2 | Raji iBZLF1 full-length 15h | To be added |
| 3 | Raji iBZLF1 full-length 15h | To be added |

[a] Replicates of the experiments
[b] Experimental section of the manuscripts
[c] Accession numbers: https://www.ebi.ac.uk/arrayexpress/experiments/browse.html
