## Supplementary Data Text for "Epstein-Barr virus inactivates the transcriptome and disrupts the chromatin architecture of its host cell in the first phase of lytic reactivation"

Alexander Buschle[1], Paulina Mrozek-Gorska[1], Stefan Krebs[2], Helmut Blum[2], Filippo M. Cernilogar[3], Gunnar Schotta[3], Dagmar Pich[1], Tobias Straub[3], and Wolfgang Hammerschmidt[1]*

[1]Research Unit Gene Vectors
Helmholtz Zentrum München
German Research Center for Environmental Health and
German Center for Infection Research (DZIF), Partner site Munich, Germany
Marchioninistr. 25
D-81377 Munich, Germany

[2]Laboratory for Functional Genome Analysis (LAFUGA), Gene Center of the Ludwig-Maximilians-Universität (LMU) München, Munich, Germany

[3]Biomedical Center, Molecular, Biology, LMU Munich, 82152 Planegg-Martinsried, Germany

*To whom correspondence should be addressed.


### Supplementary Figures

#### Figure S1

**Quantitation of BZLF1 protein in B95-8 and Raji cells**

**(A)** BZLF1 protein levels in different cell lines. The protein levels of BZLF1 (37.5 kDa) were analyzed by Western blot immunostaining with protein lysates from B95-8 cells and Raji iBZLF1 cells prior to (0h) and after induction of BZLF1 (6h). Protein lysates corresponding to different cell numbers were loaded per lane as indicated (B95-8: 40,000 to 120,000 cells; Raji 0h: 16,000 to 48,000 cells; Raji 6h: 448 to 960 cells). A truncated BZLF1 protein transiently expressed in HEK 293 cells was highly purified and 0.30, 0.75, or 1.20 ng BZLF1 protein was loaded as calibration standards to allow for the absolute quantitation of BZLF1 in the protein lysates. DG75 cells are EBV-negative cells that do not express BZLF1 and serve as a negative control.

**(B)** Absolute quantitation of BZLF1 dimers. The absolute number of BZLF1 dimers per cell were calculated based on scanned Western Blot images shown in panel A. The values are depicted in log10 scale as violin plots. The log10 median of BZLF1 dimers per cell were 6.1, 5.1 and 6.7 for the lytic fraction of B95-8 cells, non-induced (0h) and induced (6h) Raji iBZLF1 cells, respectively. In Raji iBZLF1 cells doxycycline induced the level of BZLF1 protein 42-fold within six hours. The fraction of B95-8 cells that spontaneously supports EBV's lytic phase was 2.8 % as determined by intracellular FACS analysis with an Alexa Fluor 647 coupled antibody directed against BZLF1. This value served to estimate the concentration of BZLF1 dimers in the small fraction of B95-8 cells that supports EBV's lytic phase.

**(C)** Dose-dependent release of infectious EBV from HEK 293 2089 iBZLF1 cells upon doxycycline induction. HEK 293 2089 cells (Delecluse et al., 1998) were engineered to carry the conditional BZLF1 expression plasmid p4816. Three different doxycycline concentrations (25, 100, and 200 ng/ml) were used to induce BZLF1. Non-induced cells (0 ng/ml) served as a negative control. After three days, the supernatants from HEK 293 iBZLF1 cells were harvested and the concentration of infectious virus was measured by infecting Raji cells. Green Raji units provide a measure of the concentration of infectious EBV (Steinbrück et al., 2015). Parental HEK 293 2089 cells do not release EBV above background levels (~1 %) upon addition of doxycycline. Non-induced HEK 293 iBZLF1 cells are in the same range. Mean and standard deviation from three independent experiment are shown.

#### Figure S2

**Quantitation of highly purified BZLF1 protein.**

Different volume samples of purified BZLF1 protein were loaded onto a 14 % SDS page together with the indicated amounts of BSA protein standards (125-1,500 ng). Scanned images of the gels were used to calculate the concentrations of BZLF1 protein in the samples. Two Coomassie-stained gels are shown as examples.

### Figure S3

**BZLF1 binding motifs identified in Raji iBZLF1 cells prior to and after induction of BZLF1 for 15 h.**

The sequence motifs are shown as 'sequence logos' along with the statistics, p- and e-values (Bailey, 2011) of the sub-motifs that led to the definition of the logo sequence. The sub-motifs are shown in both orientations ('Motif' and 'RC Motif'), the orientation that corresponds to the 'logo' is underlined. Sub-motifs containing a CpG pair are shown in bold. The number of motifs identified in the data set are indicated (Positive) compared with the number of motifs identified after shuffling of the data set in question (Negative) (Bailey, 2011).

**(A)** Prior to induction, when BZLF1 was expressed at low levels the TGWGCGA motif was identified in 21,387 out of 30,346 peak sequences (70.5 %).

**(B)** After 15 h of BZLF1 induction in 145,544 identified peak sequences two defined motif sets were identified that merge into the TGWGVYA motif shown in Fig. 2C.

### Figure S4

**Identification of the BZLF1 binding motif TGWGYVT in Raji iBZLF1 cells.**

**(A)** The alternative TGWGYVT motif was identified in the set of ChIP-seq peaks that contained no other previously identified BZLF1 binding motif. A search confirmed the occurrence of this motif in ChIP-seq peaks identified in Raji iBZLF1 cells. About half of the peaks in cells with a high BZLF1 level contain the TGWGYVT motif, but the related TGWGCGT motif was infrequent in ChIP-seq peaks of cells with low BZLF1 levels.

**(B)** The intersections of the two peak sets (BZLF1 levels 'low' and 'high') with the TGWGYVT motif is shown comparing their frequencies at the two different BZLF1 levels in Raji iBZLF1 cells.

**(C)** At low BZLF1 levels the BZLF1 binding motif TGWGYVT is rarely found in ChIP peaks with the BZLF1 specific BZ1 antibody, but at high BZLF1 levels this motif is abundant.

**(D)** The panel provides the average frequencies of the TGWGCGT and TGWGYVT motifs per called peak.

### Figure S5

**Relative abundance of the TGWGYVA (A) and TGWGYVT (T) motifs and the intersection (AT) of the two sets within the identified ChIP-seq peaks at high BZLF1 levels.**

The Venn diagram shows the number of peaks that contain either the TGWGYVA (A) or the TGWGYVT (T) motifs, both (AT), or none (None). In Raji iBZLF1 cells at high BZLF1 levels, 54 % of the peaks contain both (A/T) motifs.

### Figure S6

**BZLF1 binding motifs with a terminal T residue were identified in Raji iBZLF1 cells prior to and after induction of BZLF1 for 15 h.**

The figure legend follows the basics shown of Figure S4. After exclusion of peaks with sequence motifs shown in Fig. S4, the remaining peaks were reanalyzed. New sequence logos were identified and mapped back to all peak sequences as shown in Fig. S5. The statistics of these motifs and their founding sub-motifs with a terminal T residue is provided. The identified peak sequences with two defined motif sets were merged into the TGWGVYT motif shown in Fig. S5C.

### Figure S7

**Workflow of the ATAC-seq coverage at peaks of BZLF1 binding.**

The schematic overview represents the three main steps of the bioinformatic analysis, which merges ChIP-seq data with BZLF1 peak information and ATAC-seq data.

**(A)** ChIP-seq data with a BZLF1 specific antibody were obtained from doxycycline induced Raji iBZLF1 chromatin. Within the mapped ChIP-seq data the peak caller MACS2 identified 145,544 BZLF1 peaks in cellular chromatin and determined their positions on each chromosome (“chr”) indicated by dashed lines (“start”, and “end”). Four BZLF1 peaks are schematically shown.

**(B)** The BZLF1 peak positions were used to calculate the average coverage based on 145,544 BZLF1 ChIP-seq peaks. The single metaplot below summarizes the average BZLF1 peak coverage.

**(C)** ATAC-seq data were aligned according to the called nucleotide coordinates of the individual BZLF1 peaks as depicted in panel A. The resulting metaplot below gives the average coverage of the ATAC-seq signals centered at the sites of BZLF1 binding.

### Figure S8

**Average BZLF1 peak coverage and input.**

The metaplot shows the average peak coverage of all 145,544 BZLF1 peaks in Raji iBZLF1 cell chromatin 15 hours post induction. In addition, the average coverage of the input is shown at the positions of the called BZLF1 peaks. Panels A and B of Figure S7 provide the details of the bioinformatic workflow.

### Figure S9

**Heatmap of BZLF1 peaks in Raji iBZLF1 cells and input control.**

The heatmaps show the centered coverage signals at BZLF1 peaks after ChIP-seq in induced Raji iBZLF1 cells and their flanking sequences (+/- 2500 bases) in 100 bp increments.

**(A)** The heatmap of BZLF1 peaks after ChIP-seq analysis is shown.

**(B)** The heatmap of input chromatin samples at the positions of the BZLF1 peaks is shown.

### Figure S10

**Snapshot of ATAC-seq data and BZLF1 ChIP-seq data.**

**(A)** A region of about 5 kb on chromosome 7 is shown (92,051,187 to 92,056,192) in Rajji iBZLF1 cells. The ATAC-seq signals are shown prior to and after induction of the lytic cycle (lanes 1 and 3, respectively). Upon lytic induction BZLF1 binds and forms a prominent peak (lane 2) compared with the situation in non-induced cells (lane 6). As a control the AD-truncated version of BZLF1 was used prior to and after induction (lanes 4 and 5, respecively). Additionally, cellular genes are shown (lane 7). While cellular chromatin loses its accessibility, BZLF1 binding sites become accessible in ATAC-seq experiments.

**(B)** The setup follows the scheme in panel A, but depicts a 5 kb fragment on chr 1 (236,258,269 to 236,263,274). Shown in an example of an accessible region (lane 1), which stays open upon BZLF1 binding (lane 3).

**(C)** A region of about 150 kb on chromosome 12 (48,088,178 to 48,238,541) is shown. After induction of EBV’s lytic phase cellular chromatin loses its open state.

### Figure S11

**Heatmaps of the ATAC-seq coverage centered at cellular BZLF1 peaks, at randomly chosen control sequences, or at sites of open chromatin in non-induced Raji iBZLF1 cells.**

**(A)** The ATAC-seq data were collected from Raji iBZLF1 cells that express full length or AD-truncated BZLF1 at high and low BZLF1 levels. The four heatmaps show the individual ATAC-seq coverage centered at the BZLF1 peaks and their flanking sequences (+/- 2500 bases) in 100 bp increments. The sum of peak signals per 100 bp increment is shown above the heatmaps, on the right the sum of the individual peak coverage signal is provided. Figure S7 visualizes the bioinformatic workflow. The ATAC-seq coverage is strongly increased 15 h after high level expression of full-length BZLF1 (panel $A_2$) at ChIP-seq BZLF1 peaks compared with the ATAC-seq coverage at low levels of BZLF1 expression (panel $A_1$). Low or high expression levels of AD-truncated BZLF1 (panels $A_3$ and $A_4$) do not lead to a chromatin opening at BZLF1 ChIP-seq peaks. See also Figure 4A.

**(B)** As a control, 1,455,440 random sequences were chosen to analyze the global distribution of the ATAC-seq coverage. The heatmaps ($B_{1-4}$) summarize individual, random sequences that reflect the average coverage depicted in the inset of Figure 4A.

**(C)** The heatmaps ($C_{1-4}$) summarize individual sequences of the average ATAC-seq coverage shown in Figure 4B. The chromatin is open and accessible in non-induced Raji iBZLF1 cells or AD-truncated iRaji cells in both induced and non-induced conditions, but open chromatin becomes non-accessible in induced Raji iBZLF1 cells ($C_2$).

### Figure S12

**RNA-seq reads of ERCC spike-in control RNAs.**

Shown is the distribution of the ERCC spike-in RNAs in samples of Raji iBZLF1 cells prior to (0 h) and post (6 h) doxycycline induced expression of BZLF1. The x-axes provide the amount of individual RNAs (attomoles, log2 scale) added to each sample, while the y-axes show the sequencing reads, which were mapped to the individual spike-in RNAs. The color coding indicates four groups with 23 spike-in RNAs each (92 RNAs in total) and their relative abundance from low (yellow) to high (black). Of the 92 spike-in RNAs added, 63 could be identified as indicated in the shown examples. The insets provide the coefficient of determination ($R^2$), the slope of the regression, and the total number of identified ERCC spike-in RNAs (ERCCs). The data analysis shows the mean of two sets (0 h, 6 h) with three independent experiments each.

### Figure S13

**Boxplots of individual RNA-seq experiments with Raji iBZLF1 cells prior to and 6 h post BZLF1 induction.**
Boxplots of six RNA-seq samples with the raw data (left panel) and after ERCC spike-in RNA-based normalization (right panel) are provided.

### Figure S14

**KEGG pathway analysis of the 1000 most strongly down-regulated genes in Raji iBZLF1 cells.**
The top 1000 repressed genes in Raji iBZLF1 cells 6 hours after BZLF1 induction belong to important metabolic pathways or pathways characteristic of immune cells and host immune responses. Pathways related to distinct diseases or pathways with fewer than 50 genes per pathway were excluded.

### Figure S15

**Chromatin interactions of the non-regulated *CD68* gene change 6 and 15 hours after BZLF1 induction.**
The annotation of the figure is identical to Figure 8. The regions closely upstream of *CD68* loses more than half of its interactions 15 h after BZLF1 induction. The narrow grey column at about nucleotide coordinate #7,232,747 indicates the promoter region of the *NEURL4* gene, which was also studied by the Capture-C technology (Supplementary Tab. S2). The data of the *CD68* gene are not shown.

### Figure S16

**The *BTG2* gene is strongly downregulated and loses its chromatin interactions 15 hours post induction.**
The annotation of the figure is identical to Figure 8. The *BTG2* genes is downregulated by a factor of 50 upon expression of BZFL1. Prior to induction, numerous close-range chromatin interactions exist in a region flanked by two CTCF clusters with three sites each (red arrows on top of the figure bracket the region) up- and downstream of *BTG2*'s TSS. Additionally, a prominent region with increased interactions is apparent 150,000 nt downstream of the TSS.

While the interactions remain stable 6 h after BZLF1 induction, most interactions are disrupted 15 hours after induction of BZLF1.

### Figure S17

**The BZLF1-induced *COL2A1* gene initially gains chromatin interactions, which are subsequently lost.**

The annotation of the figure is identical to Figure 8. *COL2A1* is upregulated 19-fold and gains short-range, mostly intragenic chromatin interactions 6 h after BZLF1 induction, which are strongly reduced after 15 h. The induced expression of BZLF1 leads to a substantial increase of ChIP-seq peaks downstream of the gene body.

### Figure S18

**The *CXCR4* gene loses its intense long-range interactions upon BZLF1 induction.**

The annotation of the figure is identical to Figure 8. The *CXCR4* gene, which is downregulated by a factor of 25 is bracketed by two CTCF clusters (red arrows). Within these boundaries four main chromatin interaction loci indicated by red bars constitute prominent long-range interactions, all of which contain numerous BZLF1 peaks. 15 h after BZLF1 induction the majority of the long-range interactions are disrupted.

### Figure S19

**Upon BZLF1 induction the *MIR155HG* gene loses its substantial long-range interactions.**

The annotation of the figure is identical to Figure 8. When BZLF1 is not induced, the *MIR155HG* gene is characterized by three prominent regions with numerous chromatin interactions (indicated by three red arrows). The gene is downregulated by a factor of 3 upon induction of BZLF1, which binds multiple times close to the two interaction peaks. Similar to the *CXCR4* gene (Fig. S18), the long-range interactions are strongly reduced 15 h after BZLF1 induction.

### Figure S20

**Upon BZLF1 expression the *KCNQ5* gene interacts more frequently with an upstream region.**

The annotation of the figure is identical to Figure 8. In the gene body, chromatin interactions are sequentially lost when BZLF1 is induced. In a larger upstream region (red bar) chromatin interactions increase 15 h after induction.

### Figure S21

**The two adjacent genes *E2F2* (panel A) and *ID3* (panel B) have very individual chromatin interactions.**

**(A)** The annotation of the figure is identical to Figure 8. The *E2F2* promoter region frequently interacts with chromatin in the gene body and downstream of it. Upstream contacts form four peaks indicated by red arrows.

**(B)** The annotation of the figure is identical to Figure 8. The promoter interactions of the *ID3* gene clearly differ from the interactions within the *E2F2* gene.

### Figure S22

**The *TLR10* (panel A) and *TLR6* (panel B) genes are located in a distance of about 125,000 bases and harbor individual as well as shared chromatin interactions.**

The annotation of the figure is identical to Figure 8. The *TLR6* and *TLR10* genes have many individual gene-specific chromatin interactions, but share common regions (red arrows).

### Figure S23

**Two copies of a repetitive DNA sequence in the heavy chain enhancer.**

**(A)** The 323 bp long DNA sequence identified in both fragments A and B within the HCE as depicted in Figure 8 is shown. Capital, bold letters indicate BZLF1 motifs identified within this sequence. Underlined nucleotides indicate overlapping motifs.

**(B)** The BZLF1 motifs identified in the single copy of the DNA sequence shown in panel A are sorted by occurrence as indicated.

### Figure S24

**Fragments A and B within the heavy-chain-enhancer make strong contacts with the promoter of the *MYC* gene.**

**(A)** The A version of the duplicated 323 bps fragment, which is present twice in the heavy-chain enhancer (Figure 8, Figure S23) is shown in the IGV browser. Raji iBZLF1 ChIP-seq results are shown prior to and 15 h after BZLF1 induction together with the ChIP seq input control. In addition, the positions of the different BZLF1 motifs are shown.

**(B)** The B version of the duplicated fragment is shown.

### Figure S25

**The gain or loss of chromatin interactions in Capture-C experiments does not correlate with BZLF1 binding in the interacting distal fragments.**

The MA plots show the same data as in Figure 7 of the main article. In addition, the supplementary figure visualizes the occurrence or absence of BZLF1 ChIP-seq peaks in the interacting DpnII fragments. A total of 20,828 interacting fragments (0 vs 6 h, panels A1 an A2) or 16,742 (0 vs 15 h, panels B) are depicted. The orange and green dots indicate DpnII fragments with and without identified BZLF1 peaks, respectively. In panels A2 (0 vs 6 h) and B2 (0 vs 15 h) the distribution of the DpnII fragments with and without BZLF1 fragments are visualized as violin plots following the same color scheme.
